# Live-cell NanoBRET assay to measure AKT inhibitor binding to conformational states of AKT

**DOI:** 10.1101/2025.03.20.644015

**Authors:** Jeremy W. Harris, Flávio Antônio de Oliveira Simões, Erin N. Ryerson, William M. Marsiglia

## Abstract

AKT is the main protein kinase of the PI3K-AKT pathway, interacting with over one hundred protein partners to facilitate cellular processes that allow cancer cells to survive and proliferate. It is an attractive target due to its control over many cellular outputs. However, ATP-competitive and allosteric AKT inhibitors have performed poorly in clinical trials. AKT inhibitor interactions with AKT are multi-faceted and influence the catalytic activity of AKT, its conformation, its ability to interact with binding partners, and its phosphorylation state. Therefore, a better understanding of how these inhibitors influence these parameters is needed, especially in a cellular context. Using a live-cell NanoBRET target engagement assay to query the binding of AKT inhibitors to all isoforms of AKT, we found that ATP-competitive inhibitors bind similarly across all three isoforms, and allosteric inhibitors bind more heterogeneously. Further, assaying gain-of-function pathological mutants and myristoylated active versions of all AKT isoforms revealed that T308 phosphorylation enhances the binding of ATP-competitive inhibitors. We found that this phosphorylation is a good indicator of cell viability sensitivity to ATP-competitive inhibitors when comparing effects on known resistant and sensitive triple-negative breast cancer cell lines. Taken together, this assay is useful for screening new AKT inhibitors, and these findings represent important considerations in developing the next generation of AKT inhibitors.

## INTRODUCTION

AKT is a critical protein kinase that interacts with over 100 binding partners to facilitate cellular growth, proliferation, survival, and metabolism. There are three isoforms of AKT (1-3), each containing an N-terminal pleckstrin homology (PH) domain, kinase domain, and C-terminal regulatory tail (note that the use of “AKT” refers to all three isoforms). In its inactive conformation, each isoform resides in the cytoplasm and other cellular compartments, including the mitochondria and nucleus. Here, its PH domain is associated with its kinase domain to prevent interaction with binding partners^1^. Upon the generation of PIP_3_ by PI3K, the PH domain of AKT translocates to the plasma membrane (**Figure 1A**). In this active confirmation, the kinase domain is exposed for phosphorylation at residue T308/T309/T305 (AKT1/2/3) by PDK1 and at S473/S474/S472 (AKT1/2/3) by the mTORC2 complex. These phosphorylation events fully activate AKT’s catalytic activity, allowing it to regulate binding partner activity through subsequent phosphorylation events.

**Figure 1.**
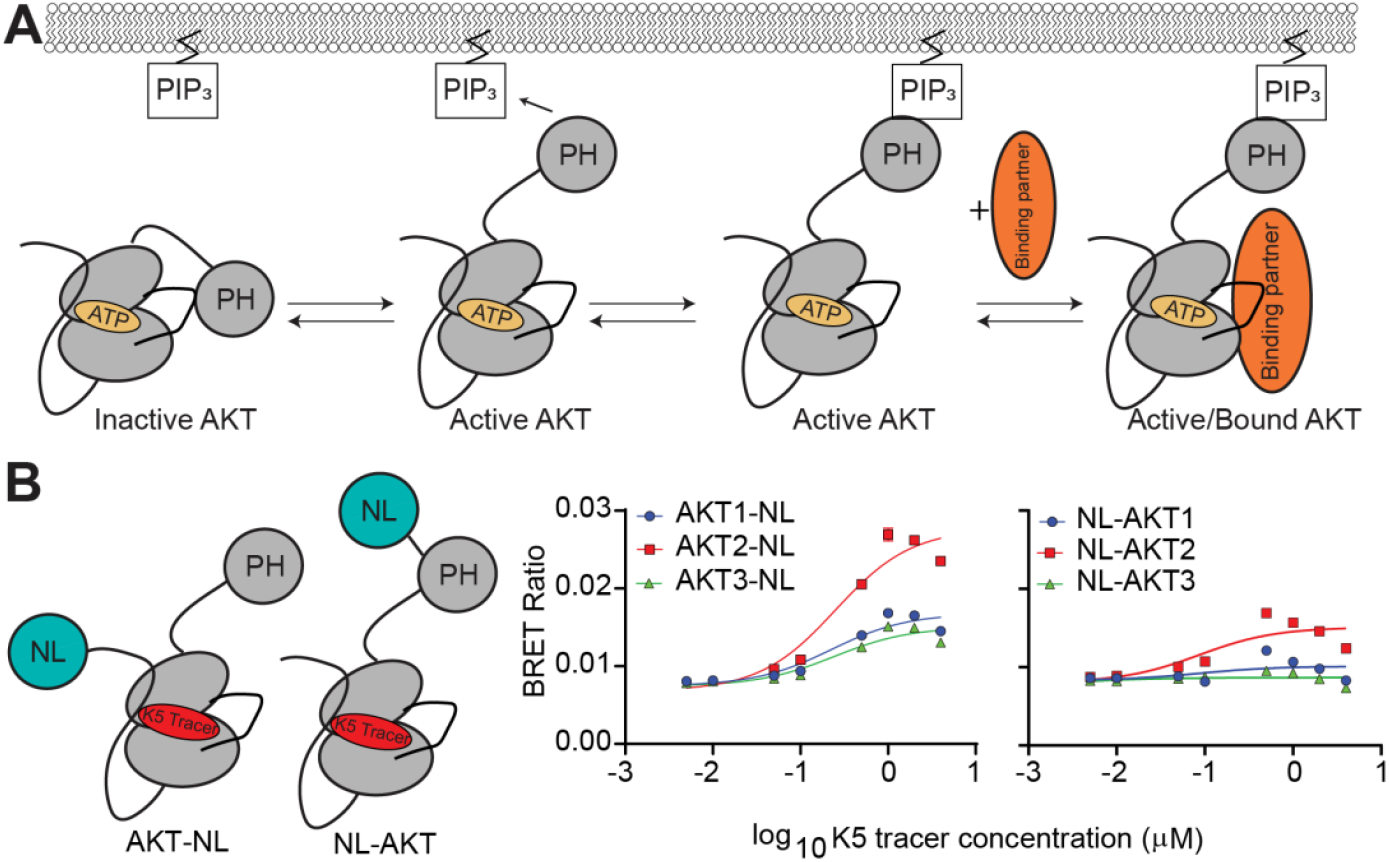
Optimum placement of NanoLuciferase on all AKT isoforms. **(A)** Schematic of AKT activation in the presence of PIP3. **(B)** Placement of Nanoluc on the C-terminus of all AKT isoforms yields a higher BRET signal than N-terminal placement. Each point for panels A, C, and D represents an average of three separate transfections (biological replicate). Each biological replicate has one technical replicate.

In cancer, AKT is aberrantly activated through PI3K-AKT pathway stimulation by upstream Receptor Tyrosine Kinases and/or PI3K/Ras mutations, and AKT gene amplification^2–5^. AKT mutations are less common but enhance the oncogenesis of breast, uterine, colorectal, and prostate cancers^6,7^. As such, AKT has stood as an attractive drug target. Whether directly targeting AKT to inhibit PI3K-AKT pathway stimulation or increased kinase activity imparted by AKT mutations, the ability to control AKT activity has promising implications for controlling oncogenic cellular processes. Two main classes of AKT kinase inhibitors are ATP-competitive (Type 1) and allosteric (Type 4). ATP-competitive inhibitors inhibit AKT through competition with ATP for AKT’s active site and putatively promote the active conformation of AKT upon binding^8^. A-443654 is a notable example that enhances membrane association and hyperphosphorylation of T308 and S473^9^. Allosteric inhibitors, such as MK-2206 and ARQ-092 (Miransertib), inhibit AKT by stabilizing its inactive conformation. Despite multiple modes of inhibition, AKT inhibitors have shown a lack of efficacy in treating cancer^10,11^.

A contributing factor underlying AKT inhibitor ineffectiveness relates to how kinase inhibitors are developed. These methods include cell viability assays that indirectly report on inhibitor binding and cell-free *in vitro* drug-binding assays using purified protein or lysates. Previous literature indicates that on-target binding in a cell-free system may not fully recapitulate binding in cells because of factors including membrane permeability, drug metabolism, target phosphorylation state etc^12,13^. In addition, since AKT inhibitors influence the conformational equilibrium of AKT between inactive and active conformations to affect its phosphorylation state indirectly and availability to binding partners, it is essential to know the affinity of AKT inhibitors towards these conformations. Obtaining this information in a cellular context is key to providing the most relevant insight for identifying and generating new inhibitors that target specific AKT isoforms and conformations prevalent in disease.

Here, we developed a NanoBRET assay to measure the binding affinity of ATP-competitive and allosteric inhibitors to different conformational states and activating mutants of AKT. We also explored the role of T308 and S473 phosphorylation on AKT inhibitor binding and the ability to use this phosphorylation as a predictor of cell viability in triple-negative breast cancer. These results represent a platform for developing future inhibitors that focus on inhibiting specific conformations of AKT.

## RESULTS AND DISCUSSION

### NanoBRET assay to measure AKT inhibitor binding across AKT isoforms

To quantify AKT inhibitor binding across all isoforms in a cellular context and correlate these results to clinical effectiveness, we performed NanoBRET competition assays. In this live-cell assay, Bioluminescence Resonance Energy Transfer (BRET) produced by the interaction of a transfected NanoLuciferase (NL)-tagged protein of interest and a protein-bound Bodipy-containing tracer molecule is dose-dependently competed out with a drug. To develop an assay for AKT, we first empirically determined if an N-terminal or C-terminal placement of NL on each AKT isoform would be optimal for BRET signal (i.e. orientation where the NL is closest to the tracer). These experiments were performed under serum-starved conditions in Opti-MEM to reduce background phosphorylation and promote an inactive population of each isoform. BRET signal buildup curves (tracer EC_50_) generated using the ATP-competitive K5 tracer from Promega indicated that the best placement (highest BRET signal) for the NL on AKT isoforms was the C-terminus (**Figure 1B and Figure S1A**). The highest BRET signal of the AKT isoforms was observed for AKT2, and the binding EC_50_ values for the K5 tracer were similar across all AKT isoforms (EC_50_: AKT1-NL 0.223 +/-0.006 μM; AKT2-NL 0.21 +/-0.02μM; AKT3-NL 0.19 +/-0.03μM).

We next characterized the binding of three representative ATP-competitive inhibitors (Ipatasertib, Capivasertib, and A-443654) and three representative allosteric inhibitors (MK-2206, Miransertib, and Inhibitor VIII) to AKT1-NL, AKT2-NL, and AKT3-NL (**Figure 2A and B, and Figure S1B-D**). We observed that the ATP-competitive inhibitors bound similarly across all three isoforms (Calculated K_i_: AKT1-NL_Ipatasertib_ 68.7 +/-14.8 nM, AKT2-NL_Ipatasertib_ 216.2 +/-55.7 nM, AKT3-NL_Ipatasertib_ 41.9 +/-10.6 nM; AKT1-NL_Capivasertib_76.0 +/-23.9 nM, AKT2-NL_Capivasertib_ 186.0 +/-39.2 nM, AKT3-NL_Capivasertib_ 87.8 +/-24.7 nM; AKT1-NL_A-443654_ 32.7 +/-14.1 nM, AKT2-NL_A-443654_ 108.2 +/-46.7 nM, AKT3-NL_A-443654_ 16.6+/-5.5 nM). MK-2206 and Miransertib bound better to AKT1 and AKT2 than AKT3 (Calculated K_i_: AKT1-NL_MK-2206_ 0.76 +/-0.09 nM, AKT2-NL_MK-2206_ 0.97 +/-0.25 nM, AKT3-NL_MK-2206_ no binding; AKT1-NL_Miransertib_ 0.27 +/-0.08 nM, AKT2-NL_Miransertib_ 2.1 +/-1.7 nM, AKT3-NL_Miransertib_ 10225 +/-7478 nM). Inhibitor VIII preferred AKT1 over AKT2 and AKT3 (Calculated K_i_: AKT1-NL_Inhibitor VIII_ 2.8 +/-0.7 nM, AKT2-NL_Inhibitor VIII_ 106.3 +/-6.6 nM, AKT3-NL_Inhibitor VIII_ 438.7 +/-72.3 nM). The reduced potency of allosteric inhibitors for AKT3 is consistent with literature showing that the upregulation of AKT3 is associated with MK-2206 resistance in breast cancer^14^, and higher kinase activity of AKT3 relative to AKT1 and 2 in multiple cell lines when exposed to MK-2206^15^. The isoform differences observed for the allosteric inhibitors could be explained by differences in the primary sequences among the isoforms that make up the inhibitor binding site^16,17^. It should be noted that the differences in calculated K_i_ values between AKT1 and AKT3 for allosteric inhibitors are orders of magnitude larger than values previously reported using other biochemical methods but follow the same potency trends^15,18^.

**Figure 2.**
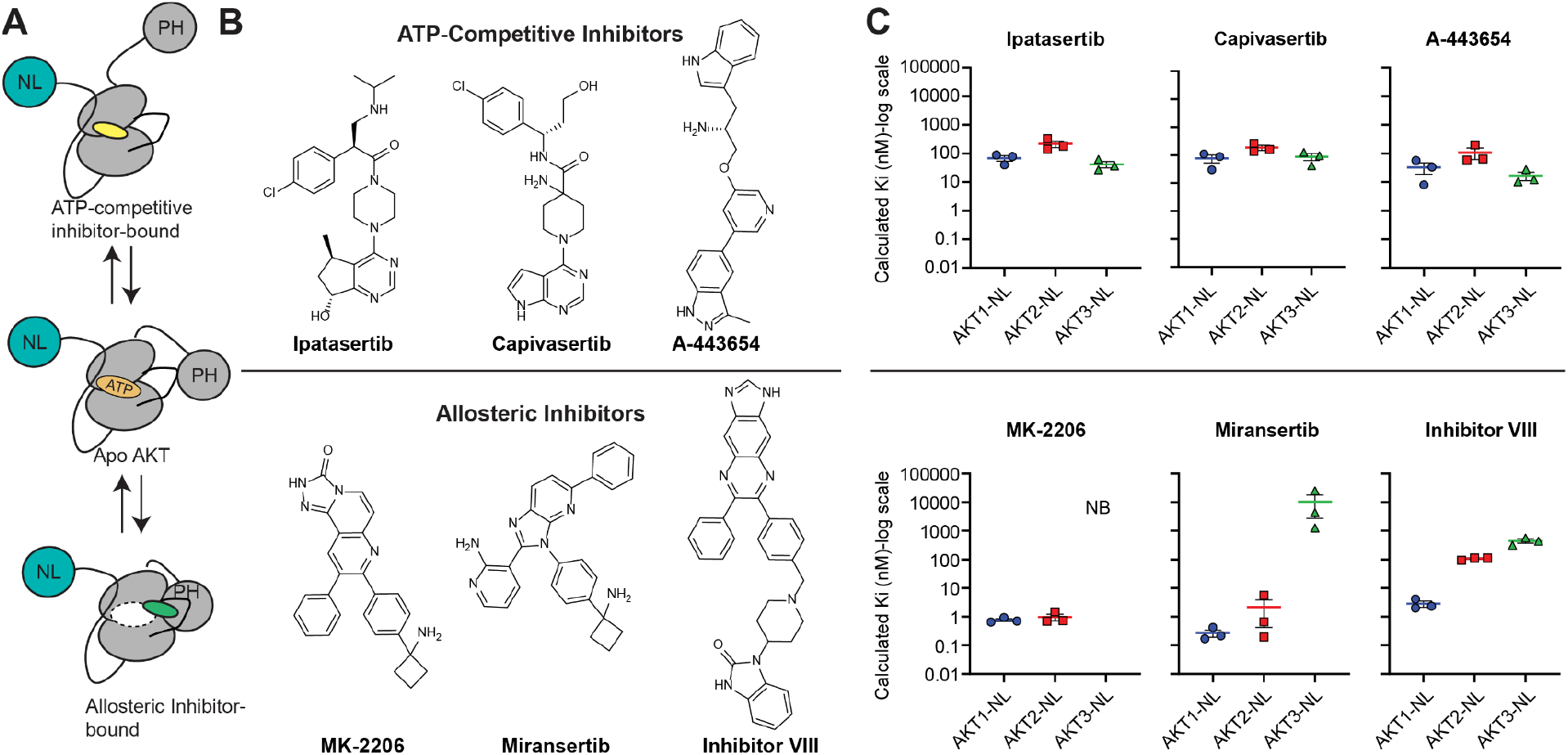
Characterization of ATP-competitive and Allosteric AKT inhibitors on all AKT isoforms. **(A and B)** Schematic of binding modes and structures for ATP-competitive (Ipatasertib, Capivasertib, and A443654) and Allosteric (MK-2206, Miransertib, and Inhibitor VIII) AKT inhibitors. The dotted oval in the allosteric-bound diagram represents the ATP-binding pocket. **(C)** Calculated Ki values for ATP-competitive (top) and allosteric inhibitors. Note that “NB” indicated no binding. Each point for panel C represents a separate transfection (biological replicate). Each biological replicate has one technical replicate.

### Characterization of AKT inhibitor binding to pathogenic AKT mutants

We were curious to see how the same set of inhibitors bound to pathological gain-of-function mutants that shift AKT toward a higher active population. Mutations of AKT are found in breast, prostate, colorectal, glioma, endometrial, bladder, lung, and melanoma cancers^19–22^. AKT1 has a greater frequency of mutations than AKT2 and AKT3^23^. These mutations are generally located near the interface between the PH and kinase domains (L52R and D323H), the PIP_3_ binding pocket (E17K), and the drug-binding pocket of allosteric inhibitors (Q79K and W80R) (**Figure 3A**). In the context of AKT1-NL, we quantified ATP-competitive and allosteric inhibitor binding to E17K, L52R, Q79K, W80R, and D323H. We also queried the E17K mutant in the context of AKT2-NL and AKT3-NL. Compared to wild-type (WT) AKT1-NL, we observed enhanced binding of ATP-competitive inhibitors to all mutants (**Figure 3B and Figure S2**). All allosteric inhibitors bound worse to all mutants relative to WT-AKT1-NL (**Figure 3C and Figure S2**). Notably, no binding of MK-2206 was observed to the Q79K mutant, a key residue that is part of the MK-2206 binding site. Miransertib and Inhibitor VIII, however, bound better to Q79K, highlighting inhibitor selectivity towards specific mutants. There was no significant difference in ATP-competitive inhibitor binding between WT and E17K versions of AKT2-NL and AKT3-NL (**Figure S3**). MK-2206 did not bind to AKT3-NL or AKT3-E17K-NL, and Miransertib did not bind to AKT3-E17K-NL. The weaker binding of allosteric inhibitors to AKT1 mutants can be explained through a combination of mutation location (i.e. near the inhibitor binding site) and destabilization of the kinase-PH domain interface that makes up the inhibitor binding site^19^.

**Figure 3.**
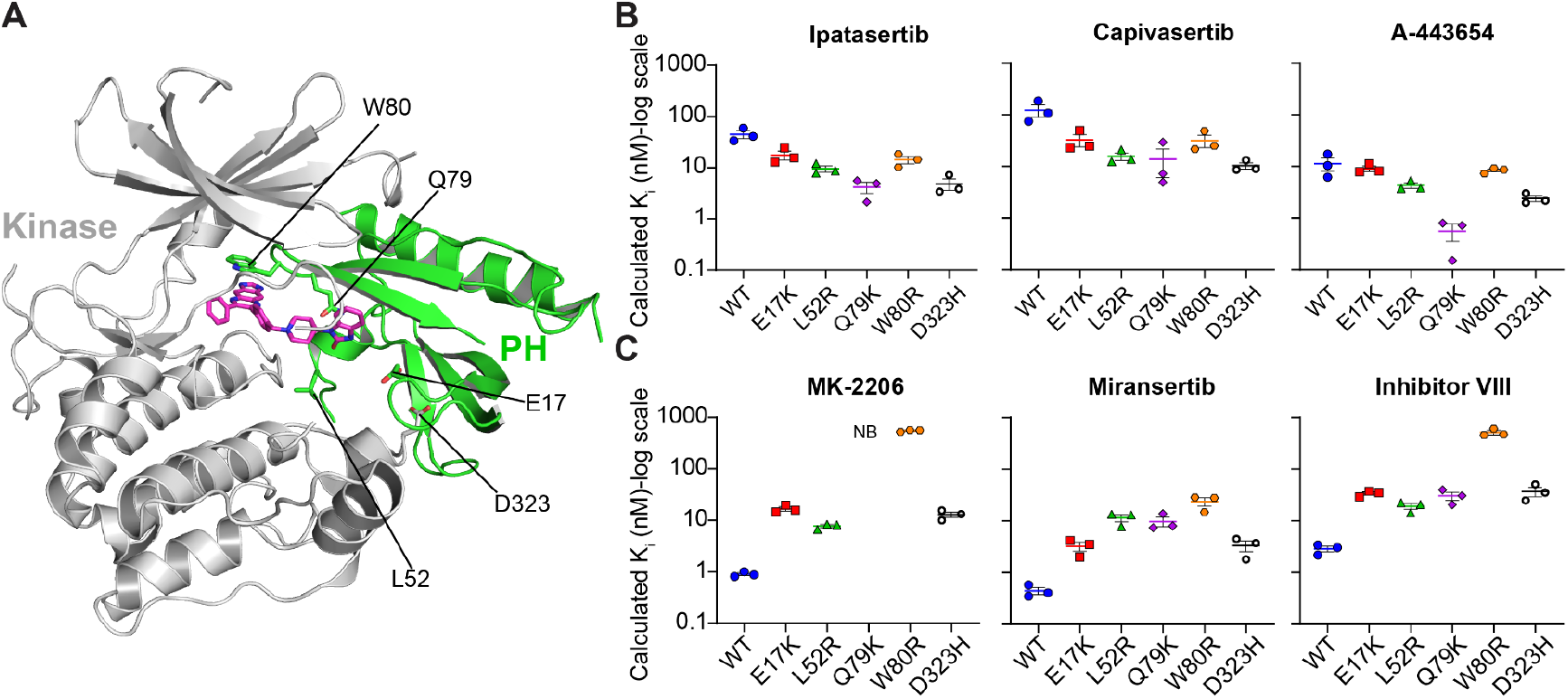
Comparison of AKT inhibitor binding on pathogenic mutations. **(A)** Structure of AKT2 bound to Inhibitor VIII (PDB ID:3O96) highlighting the location of pathogenic AKT pathogenic mutations. **(B)** Calculated Ki values for ATP-competitive inhibitors on pathogenic mutants. **(C)** Calculated Ki values for allosteric inhibitors for pathogenic mutants. Note that “NB” indicated no binding. Each point for panels B and C represents a separate transfection (biological replicate). Each biological replicate has one technical replicate.

### Residue T308 in AKT1 is important for the binding of ATP-competitive AKT inhibitors

It was puzzling why ATP-competitive inhibitors bound more tightly to some pathological mutants. Since these mutants have been shown to increase AKT activity by enhancing T308 and S473 phosphorylation^24^, we reasoned that changes in phosphorylation and conformation from inactive to active states may influence inhibitor binding. It was previously shown that unstimulated AKT primarily resides in the cytoplasm inactive and unphosphorylated until increased PIP_3_ levels are present^25^. To achieve an active AKT population characterized by increased phosphorylation and membrane association, we created N-terminally myristoylated AKT1-3-NL constructs (**Figure 4A**). AKT myristoylation is commonly used to keep AKT membrane-associated and available for phosphorylation by PDK1 and mTORC2^26,27^. K5 tracer binding was similar across isoforms and similar to the non-myristoylated versions of each isoform (K5 EC_50_: AKT1-NL 0.223 +/-0.006 μM; Myr-AKT1-NL 0.18 +/-0.02 μM; AKT2-NL 0.21 +/-0.02 μM; Myr-AKT2-NL 0.124 +/-0.007 μM; AKT3-NL 0.19 +/-0.03 μM; Myr-AKT3-NL 0.16 +/-0.01 μM) (**Figure S1A** and **Figure S4A**). We then measured the binding of the same ATP-competitive and allosteric inhibitors on non-myristoylated and myristoylated versions of AKT1-3-NL (**Figure 4B and Figure S4**). Relative to the non-myristoylated protein, we observed a ∼6-fold enhancement in the potency of ATP-competitive inhibitors to AKT1-NL vs. Myr-AKT1-NL on average. A smaller change between AKT2-NL and Myr-AKT2-NL (∼2-fold) and no significant change in AKT3-NL vs Myr-AKT3-NL were also observed. These results suggest that ATP-competitive inhibitors are isoform-selective in their preference for active and membrane-associated AKT.

**Figure 4.**
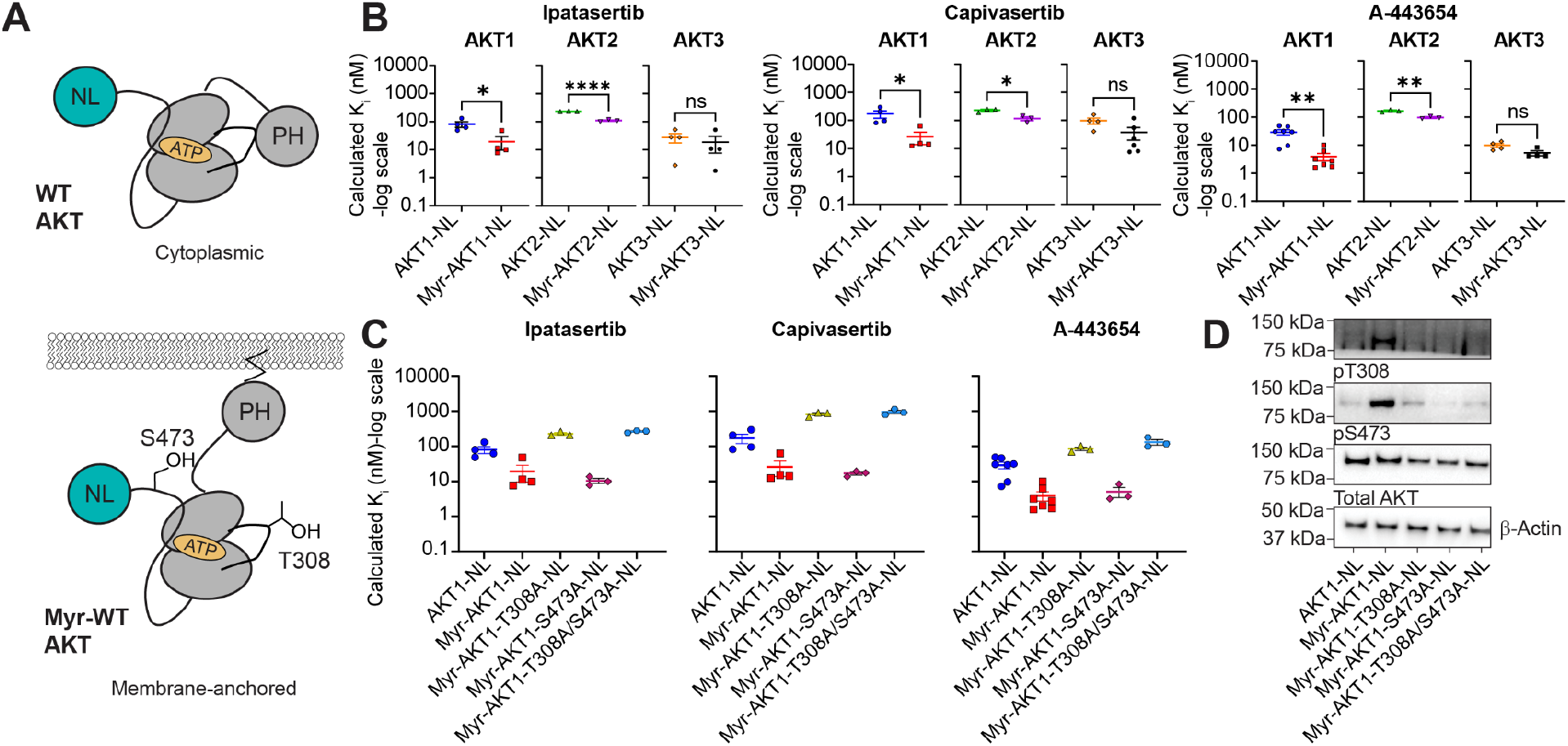
Influence of residues T308 and S473 on AKT inhibitor binding. **(A)** Schematic showing WT-AKT-NL (top) and Myristoylated AKT-NL constructs (bottom). Phosphorylatable residues T308 and S473 are shown using AKT1 numbering. **(B)** Calculated Ki comparison of ATP-competitive binding to WT and Myristoylated versions of each AKT isoform. **Ipatasertib**: AKT1-NL vs. Myr-AKT1-NL, *P*=0.286, *U*=0, two-tailed Mann-Whitney *U*-test; AKT2-NL vs. Myr-AKT2-NL, *P*=<0.001, *t*=20.41, *df*=4, two-tailed unpaired *t*-test; AKT3-NL vs. Myr-AKT3-NL, *P*=0.5749, *t*=0.5928, *df*=6, two-tailed unpaired *t*-test. **Capivasertib**: AKT1-NL vs. Myr-AKT1-NL, *P*=0.286, *U*=0, two-tailed Mann-Whitney *U*-test; AKT2-NL vs. Myr-AKT2-NL, *P*=0.0134, *t*=4.231, *df*=4, two-tailed unpaired *t*-test; AKT3-NL vs. Myr-AKT3-NL, *P*=0.1143, *U*=4, two-tailed Mann-Whitney *U*-test. **A-443654**: AKT1-NL vs. Myr-AKT1-NL, *P*=0.0023, *U*=2, two-tailed Mann-Whitney *U*-test; AKT2-NL vs. Myr-AKT2-NL, *P*=0.0033, *t*=6.294, *df*=4, two-tailed unpaired *t*-test; AKT3-NL vs. Myr-AKT3-NL, *P*=0.1143, *U*=2, two-tailed Mann-Whitney *U*-test. **(C)** Comparison of calculated Ki values of ATP-competitive inhibitor binding to WT-AKT1-NL and versions of AKT that cannot be phosphorylated at specific residues to determine the source of enhanced ATP-competitive inhibitor binding to Myr-AKT1-NL. Ki values were calculated using measured K5 EC50 values in Figure S1A and S4A, and apparent IC50 values in Figure S4A using the Cheng-Prusoff equation. Note that the data for AKT1-NL and Myr-AKT1-NL in panels B and C is the same due to how the data was collected. **(D)** Western blot analysis confirming the phosphorylation states of T308 and S473 in each construct. ^*^*P* < 0.05; ^**^*P* < 0.01; ^****^*P* < 0.0001. Each point for panels B and C represents a separate transfection (biological replicate). Each biological replicate has one technical replicate.

Since the AKT1 isoform showed a larger change in ATP-competitive inhibitor binding affinity than the other isoforms, we aimed to determine the influence of T308 and S473 phosphorylation sites in its myristoylated context. To do this, we mutated each site separately (T308A or S473A) and in combination (T308A/S473A) to alanine (**Figure 4C and Figure S4**). Indeed, western blot analysis of AKT1-NL, Myr-AKT1-NL, T308A-Myr-AKT1-NL, S473A-Myr-AKT1-NL, and T308A/S473A-Myr-AKT1-NL revealed that alanine mutation prevented phosphorylation of its corresponding residue (**Figure 4D and Figure S5**). We observed that the mutation of T308 (T308A-Myr-AKT1-NL) alone or in combination with S473 (T308A/S473A-Myr-AKT1-NL) was sufficient to reverse the increase in binding potency of ATP-competitive inhibitors and resembled binding to AKT1-NL. The binding affinities of ATP-competitive inhibitors to AKT1 harboring the S473A mutation matched more closely to the Myr-AKT1-NL construct. The allosteric connection between the ATP binding site and T308 has been previously shown where the presence of ATP or A-443654 promotes the resistance of pT308 to dephosphorylation through long-range interactions mediated by N-lobe to C-lobe closure. This closure places residues on the αC-helix in position to sequester pT308 away from solvent^28^. These results are the first representation of this allostery in the context of live-cell target engagement. Therefore, we conclude that residue T308 but not S473 allosterically enhances the binding of ATP-competitive inhibitors to AKT.

The same NanoBRET experiments were then performed using allosteric inhibitors, and worse binding was observed for all myristoylated constructs regardless of mutation relative to AKT1-NL (**Figure S4D**). In general, myristoylation reduced calculated K_i_ values for each allosteric inhibitor 10-100-fold. This result indicates that myristoylation makes it more difficult for allosteric inhibitors to stabilize the inactive form of AKT. The weaker binding of allosteric inhibitors, combined with stronger binding to ATP-competitive inhibitors, represents a characteristic inhibitor “profile” of Myr-AKT-NL constructs.

To ensure that changes in ATP-competitive inhibitor binding affinity were not from generally mutating an amino acid, we mutated residue C344S on Myr-AKT1-NL (**Figure S4**). This residue is located at the C-terminal end of the αF helix in the kinase C-lobe, away from residues that could influence kinase conformation, inhibitor binding sites, and catalytic residues. Western blot analysis of T308 and S473 phosphorylation in the C344S-Myr-AKT1-NL construct. This result matched closely with Myr-AKT1-NL and confirmed a minimal effect of the mutation (**Figure S5**). We also observed from our NanoBRET assay that the C344S-Myr-AKT1-NL mutant retained the ATP-competitive inhibitor binding affinity enhancement displayed by the Myr-AKT1-NL construct. ***The starting population of pT308 is important for ATP-competitive inhibitor binding***

In light of determining the role of T308 in enhancing the binding of ATP-competitive inhibitors, we wanted to see if we could shift the populations of phosphorylated T308, and thus ATP-competitive binding using a PDK1 inhibitor (BX-795). In other words, we were curious to see if BX-795 would influence ATP-competitive inhibitor binding to AKT1-NL (low pT308) and Myr-AKT1-NL (high pT308) (**Figure 5A**). To test this, we performed a steady-state NanoBRET competition assay for A-443654 in the presence of a constant concentration of BX-795 (1 μM). The concentration of BX-795 was determined empirically from dose-response curves of BX-795 on AKT1-NL and Myr-AKT1-NL (**Figure 5B and Figure S6A**). Notably, some binding of BX-795 to AKT1-NL and Myr-AKT1-NL was observed at concentrations higher than 1 μM. NanoBRET competition assays of A-443654 in the presence and absence of BX-795 for AKT1-NL and Myr-AKT1-NL revealed that A-443654 binding to AKT1-NL was unaffected by BX-795, and that A-443654 bound more weakly to Myr-AKT1-NL in the presence of BX-795 (**Figure 5C and Figure S6B-D**). This reversal of ATP-competitive inhibitor binding enhancement agrees well with the effect of the T308A-Myr-AKT1-NL mutant In **Figure 4C**.

**Figure 5.**
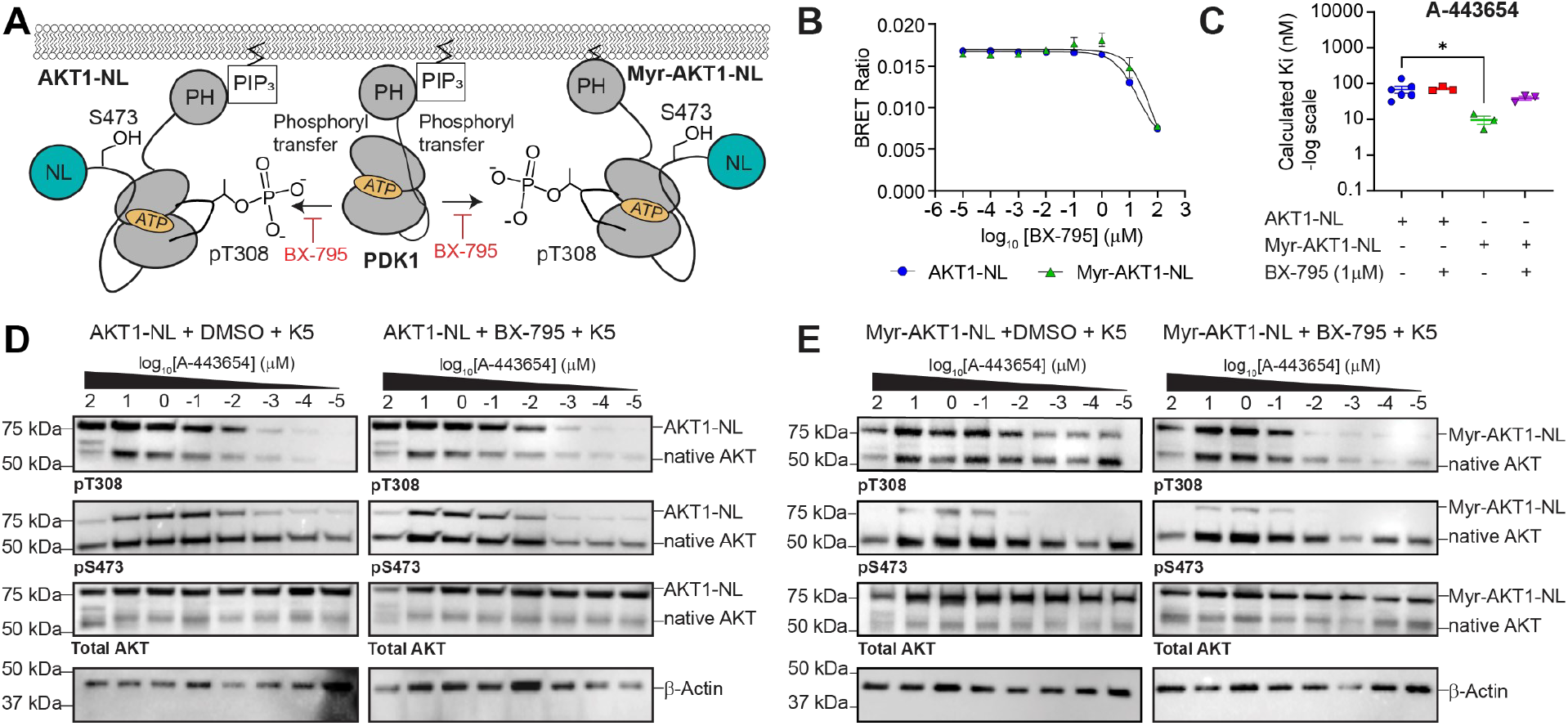
Pharmacological modulation of AKT phosphorylation and AKT inhibitor binding. **(A)** Schematic showing the role of BX-795 (PDK1 inhibitor) in blocking T308 phosphorylation on AKT1-NL and Myr-AKT1-NL. **(B)** Dose-response curves of BX-795 on AKT1-NL and Myr-AKT1-NL. **(C)** Calculated Ki values for A-443654 on AKT1-NL and Myr-AKT1-NL in the presence and absence of 1 μM BX-795. Data were analyzed by one-way ANOVA with Dunnett’s multiple comparison test (AKT1-NL vs. AKT1-NL + 1 μM BX-795 adjusted *P*=0.9982, AKT1-NL vs. Myr-AKT1-NL adjusted *P*=0.0234, AKT1-NL vs. Myr-AKT1-NL + 1 μM BX-795 adjusted *P*=0.3293). Note that the final DMSO concentration is higher in all conditions relative (0.2% DMSO) to that in Figure 4B (0.1% DMSO) and may contribute to differences in calculated Ki values. Importantly, the trends are the same. **(D)** Western Blot analysis of AKT1-NL and AKT1-NL + 1 μM BX-795 in the presence of an A-443654 dose-response curve. **(E)** Western Blot analysis of Myr-AKT1-NL and Myr-AKT1-NL + 1 μM BX-795 in the presence of an A-443654 dose-response curve. ^*^*P* < 0.05; ^**^*P* < 0.01; ^****^*P* < 0.0001. Each point for panel C represents a separate transfection (biological replicate). Each biological replicate has one technical replicate.

Given that the addition of A-443654 induces AKT hyperphosphorylation, we aimed to quantify the contribution of T308 and S473 hyperphosphorylation on the steady-state NanoBRET derived K_i_ values of A-443654 observed for AKT1-NL and Myr-AKT1-NL in the presence/absence of BX-795. We performed a Western blot analysis on a larger scale NanoBRET assay where 293T cells transfected with either AKT1-NL or Myr-AKT1-NL were separately incubated with each concentration of an A-443654 dose-response curve (100, 10, 1, 0.1, 0.01, 0.001, 0.0001, or 0.00001 μM), 1 μM K5 tracer, and either 1 μM of BX-795 or DMSO for 2 hrs at 37°C. For AKT1-NL, we observed that A-443654 created a dose-dependent increase in T308 and S473 phosphorylation, which was unaffected by BX-795 (**Figure 5D and Figure S7A-B**). Myr-AKT1-NL also showed a dose-dependent increase in T308 phosphorylation in the presence and absence of BX-795 (**Figure 5D and Figure S7C-D**). However, the addition of BX-795 strongly reduced the phosphorylation of T308 observed at lower concentrations of A-443654. This result correlates well with the steady-state NanoBRET experiments that show less A-443654 binding in the presence of BX-795.

Taken together, the K_i_ measurements from the NanoBRET steady-state experiments stem from an intrinsic inhibitor binding affinity modulated by AKT’s starting phosphorylation state population and the inhibitors’ induced phosphorylation changes. Nonetheless, the take-home message is that the binding affinity of ATP-competitive inhibitors is stronger for AKT when phosphorylated at T308.

### T308 phosphorylation plays a role in triple-negative breast cancer sensitivity to ATP-competitive AKT inhibitors

Triple-negative breast cancer subtypes are genetically heterogeneous and have different sensitivities to AKT inhibitors^29–31^. The MDA-MB-231 cell line is derived from a triple-negative breast cancer patient harboring mutations in the RAS-MAPK pathway in addition to additional alterations of P53. MDA-MB-468 is a cell line derived from a triple-negative breast cancer patient with deletion of PTEN that leads to the overstimulation of the PI3K-AKT pathway. It has been previously reported that the MDA-MB-231 cell line is resistant to AKT inhibitors and the MDA-MB-468 is sensitive to AKT inhibitors^31^. Based on data presented in **Figures 4 and 5**, we were curious to see if intrinsic differences in AKT phosphorylation between MDA-MB-231 and MDA-MB-468 cell lines would display differences in AKT inhibitor binding. We first tested the effects of Ipatasertib, Capivasertib, A-443654, MK-2206, Miransertib, and Inhibitor VIII on the cell viability of both cell lines. We found that the MDA-MB-468 cell line was more sensitive to ATP-competitive inhibitors and similarly as sensitive to allosteric inhibitors relative to the MDA-MB-231-cell line (**Figure 6A and Figure S8**). The largest difference in inhibitor sensitivity between the two cell lines was observed for A-443654 (IC_50_: MDA-MB-231 0.33 +/-0.01 μM; MDA-MB-468 0.020 +/-0.001 μM).

**Figure 6.**
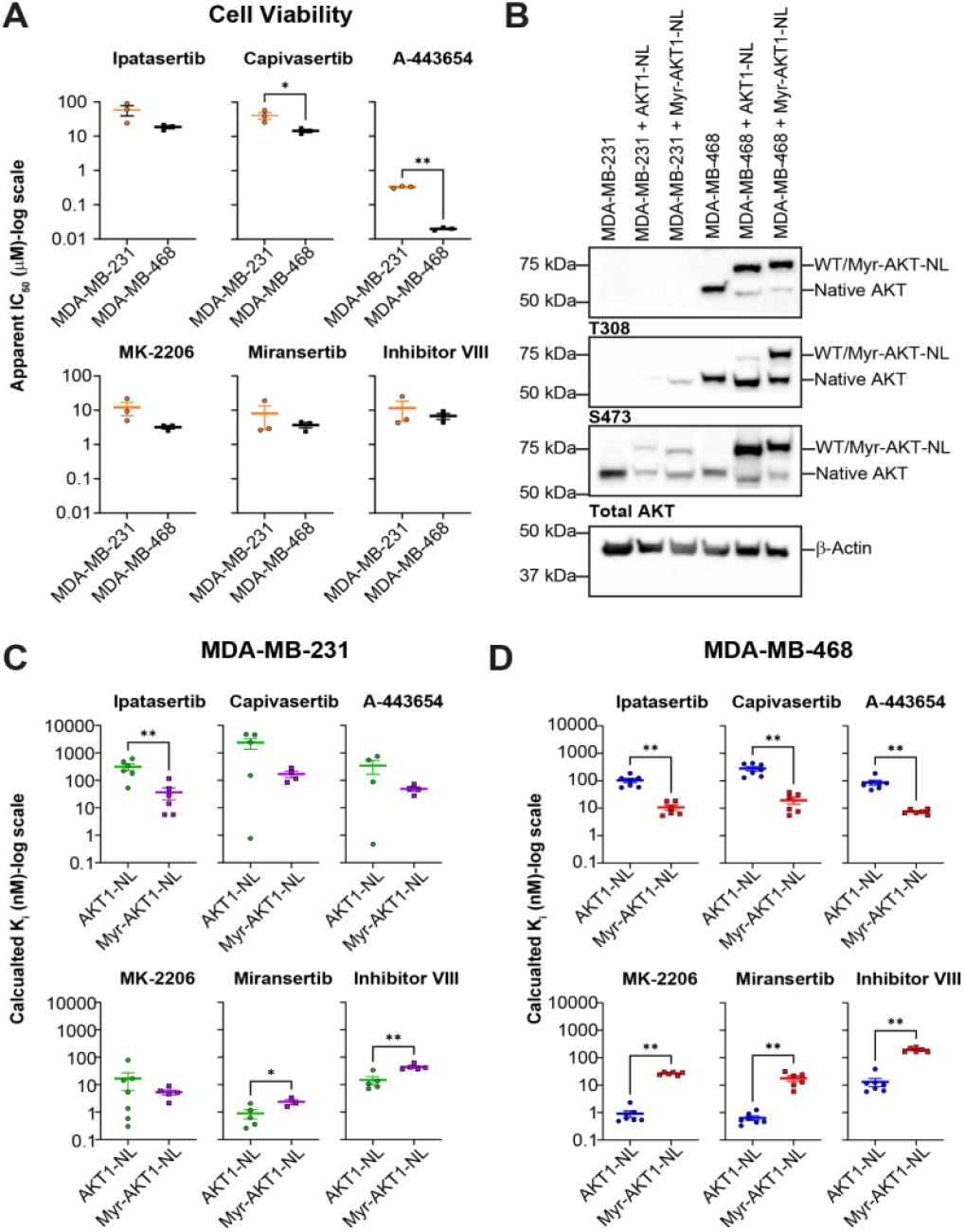
Correlation of cell viability and target engagement of AKT inhibitors sensitive and resistant triple-negative breast cancer cell lines. **(A)** Cell viability comparison of MDA-MB-231 and MDA-MB-468 in the presence of AKT inhibitors. **Capivasertib:** MDA-MB-231 vs MDA-MB-468, *P*=0.0483, *t*=2.811, *df*=4, two-tailed unpaired *t*-test. **A-443654:** MDA-MB-231 vs MDA-MB-468, *P*=0.0021, *t*= 21.14, *df*=2.023, two-tailed Welch’s *t*-test. **(B)** Western blot comparing the phosphorylation state of T308 and S473 in MDA-MB-231 and MDA-MB-468 cell lines alone or transfected with either AKT1-NL or Myr-AKT1-NL. **(C)** NanoBRET target engagement assays of AKT inhibitors on MDA-MB-231 cells transfected with either AKT1-NL or Myr-AKT1-NL. **Ipatasertib:** AKT1-NL vs. Myr-AKT1-NL, *P*=0.0043, *U*=1, two-tailed Mann-Whitney *U*-test. **Miransertib:** AKT1-NL vs. Myr-AKT1-NL, *P*=0.0714, *U*=1, two-tailed Mann-Whitney *U*-test. **Inhibitor VIII:** AKT1-NL vs. Myr-AKT1-NL, *P*=0.0025, *t*=4.347, *df*=8, two-tailed unpaired *t*-test. **(D)** NanoBRET target engagement assays of AKT inhibitors on MDA-MB-468 cells transfected with either AKT1-NL or Myr-AKT1-NL. **Ipatasertib:** AKT1-NL vs. Myr-AKT1-NL, *P*=0.0012, *U*=0, two-tailed Mann-Whitney *U*-test. **Capivasertib:** AKT1-NL vs. Myr-AKT1-NL, *P*=0.0012, *U*=0, two-tailed Mann-Whitney *U*-test. **A-443654:** AKT1-NL vs. Myr-AKT1-NL, *P*=0.0012, *U*=0, two-tailed Mann-Whitney *U*-test. **MK-2206:** AKT1-NL vs. Myr-AKT1-NL, *P*=0.0012, *U*=0, two-tailed Mann-Whitney *U*-test. **Miransertib:** AKT1-NL vs. Myr-AKT1-NL, *P*=0.0012, *U*=0, two-tailed Mann-Whitney *U*-test. **Inhibitor VIII:** AKT1-NL vs. Myr-AKT1-NL, *P*=0.0012, *U*=0, two-tailed Mann-Whitney *U*-test. ^*^*P* < 0.05; ^**^*P* < 0.01; ^****^*P* < 0.0001. Each point for panels A, C, and D represents a separate transfection (biological replicate). Each biological replicate has one technical replicate.

To understand the contribution of phosphorylation to these results, we performed a Western blot analysis to query the phosphorylation state of T308 and S473. In preparation for steady-state NanoBRET experiments, we separately transfected AKT1-NL and Myr-AK1-NL into each cell line. We found that our transfected constructs were not phosphorylated in MDA-MB-231 cells, but were in MDA-MB-468 cells (**Figure 6B and Figure S9**). In MDA-MB-468 cells, Myr-AKT1-NL showed 1.7x increased phosphorylation (see methods) of T308 compared to AKT1-NL. The lack of phosphorylation in the MDA-MB-231 cell line is consistent with previous literature^32^. To test if cell viability differences between cell lines correlated with differences in target engagement to different AKT conformations, we transfected AKT1-NL or Myr-AKT1-NL into both cell lines. We performed steady-state NanoBRET competition assays for the same AKT inhibitors. Interestingly for MDA-MB-231 cells, we observed similar binding between AKT1-NL and Myr-AKT1-NL for almost all inhibitors (**Figure 6C and Figure S10**). In agreement with experiments performed using HEK293T cells, we observed an increase in ATP-competitive inhibitor binding affinity and a decrease in allosteric inhibitor binding for MDA-MB-468 cells (**Figure 6D and Figure S10**). It is important to note that although there were larger differences in the binding of allosteric inhibitors to AKT1-NL and Myr-AKT1-NL in MDA-MB-468 vs. MDA-MB-231 cell lines, these differences did not translate to effects on cell viability. These results highlight the utility of using T308 phosphorylation to predict the effects of ATP-competitive inhibitors on cell viability. These results also indicate that the steady-state NanoBRET assay can be used to report on differences in genetic backgrounds between cell lines.

## CONCLUSIONS

In conclusion, we have demonstrated a NanoBRET target engagement assay to query the binding of ATP-competitive and allosteric AKT inhibitors to all AKT isoforms. We have shown that ATP-competitive inhibitors bind similarly to all isoforms, and that allosteric inhibitors are generally selective for AKT1 and AKT2 over AKT3. Using a combination of pathogenic and non-phosphorylatable mutants, and an upstream PDK1 inhibitor, we found that phosphorylation of AKT1 at T308 enhances the binding of ATP-competitive inhibitors. These results are the first to measure the direct binding of AKT inhibitors to AKT in a cellular context and represent a platform for screening new inhibitors. We also highlight that AKT gain-of-function mutations bind better to ATP-competitive inhibitors, and that T308 phosphorylation is an important consideration in developing inhibitors that target specific states of AKT and as a marker to indicate the potential effectiveness of ATP-competitive inhibitors.

## METHODS

### Materials

Myristoylated constructs of human AKT1 (UniProt: P31749), AKT2 (UniProt: P31751), and AKT3 (UniProt: Q9Y243) were purchased from Addgene (cat# 9008, 9016, and 9017, respectively). pNLF1-N (cat# N1351) and pNLF1-C (cat# N1361) vectors, K5 10,000x NanoBRET kits (cat# N2530), K16 1000x NanoBRET Kit (cat# CS1810C495), PDK1-NL vector (cat# CS1810C771), and GloMax Discover (cat# GM3000) were purchased from Promega. QuikChange mutagenesis kits were purchased from Agilent (cat# 200522). Ipatasertib (cat# HY-15186), Capivasertib (cat# HY-15431), MK-2206 (cat# HY-10358), Miransertib (cat# HY-19719), and Inhibitor VIII (cat# HY-10355) were purchased from MedChem Express. A-443654 was purchased from Ontario Chemicals (cat# I1939). Resazurin was purchased from Fisher Scientific (cat# AC418900050). White 96-well plates for NanoBRET experiments (cat# 07200628), and clear plastic 96-well plates (cat# 229195) for drug dilutions and cell viability experiments were purchased from Fisher Scientific. For cell culture, Opti-MEM (cat# 11-058-021), DMEM (cat# 11-965-092), FBS (cat# 26140079), Trypsin (cat# 25200056), Penicillin/Streptomycin (cat# 15140122) and PBS (cat# 20-012-027) were purchased from ThermoFisher Scientific. For transfections, PEI Star (cat# 78-541-00) was purchased from Fisher Scientific, and Fugene HD was purchased from Promega (cat# E2311). For western blots, BSA (cat# A9647), Thermo Scientific™ Halt™ Protease and Phosphatase Inhibitor Cocktail (100X) (cat# PI78440), RIPA buffer (cat# 89900), and SuperSignal West Femto Substrate Kit (cat# 34096) were purchased from Fisher Scientific. Pierce BCA Protein Assay Kits (cat# A55864) and Trans-Blot Turbo RTA Transfer Kits (cat# 1704273) were purchased from Bio-Rad. Antibodies Phospho-AKT(Thr308) (D25E6) XP Rabbit mAb (cat# 13038), Phospho-AKT (Ser473) (D9E) XP Rabbit mAb (cat# 4060), AKT (pan) (C67E7) Rabbit mAb (cat# 4691), β-Actin (D6A8) Rabbit mAb (cat# 8457), and Anti-rabbit IgG-HRP-linked Antibody (cat# 7074) were purchased from Cell Signaling. The HEK293T cell line was acquired from ATCC (cat# CRL-3216), and the MDA-MB-231 and MDA-MB-468 cell lines were generously provided by Dr. Natalie R. Gassman.

### Constructs and Mutagenesis

To generate N- or C-terminal NanoLuciferase fusion constructs, full-length versions of AKT1, AKT2, and AKT3, were separately subcloned into pNLF1-N or pNLF1-C vectors between PvuI and XbaI, and PvuI and XhoI, respectively. Myristoylated versions of AKT1, AKT2, and AKT3 were only subcloned into the pNLF1-C vector. PDK1-NL was purchased from Promega. All pathological and myristoylated mutants were made using the QuikChange mutagenesis kit.

### Steady-state NanoBRET competition Assays

HEK293T, MDA-MB-231, or MDA-MB-468 cells were transfected at a cell density of 200,000 cells/mL in Opti-MEM (containing 10% FBS and 1000 U/mL Pen/Strep) using either PEI Star or Fugene HD, and a ratio of 9.5:0.5 μg/mL carrier DNA:AKT-NL construct in Opti-MEM. This solution was aliquoted into white adherent 96-well plates at a volume of 100 μL/well and incubated for 24 hrs at 37°C and 5% CO_2_. For steady-state apparent IC_50_ measurements, 10 μL of a 10x AKT inhibitor/BX-795 dose-response curve (100, 10, 1, 0.1, 0.01, 0.001, 0.0001, 0.00001 μM final concentration at 0.1% DMSO) made in Opti-MEM, followed by 5 μL of a 20x K5 tracer solution (made according to Promega’s adherent target engagement protocol). For steady-state K5 EC_50_ measurements, 5 μL of 20x K5 tracer dose-response curve (4, 2, 1, 0.5, 0.1, 0.05, 0.01, 0.005 μM final concentration) was added to transfected cells. For steady-state IC_50_ measurements of AKT inhibitors in the presence of BX-795, 10 μL of a 10x dose-response for each AKT inhibitor was added to each well followed by 10 μL of BX-795 (1 μM final) and 5 μL of a 20x K5 tracer solution (0.2% DMSO final). Following all inhibitor and tracer additions, cells were incubated at 37°C and 5% CO_2_ for 2 hrs. 50 μL of a 3x solution of Opti-MEM containing a NanoGlo substrate and NanoLuciferase inhibitor provided in the NanoBRET kit were added to each well. BRET data were collected on a GloMax Discover using the preset NanoBRET 618 protocol, exported in Microsoft Excel format, and analyzed in GraphPad Prism 10. Statistics were performed in Origin 2023. Calculated K_i_ values were calculated using the following formula:

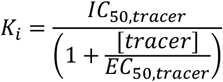

### Western Blot analysis of AKT phosphorylation

1.1 million cells in 5.28 mL of Opti-MEM were seeded into 35 mm tissue culture-treated dishes transfected using either polyethylenimine or Fugene HD, and a ratio of 9:1 ug/mL carrier DNA:AKT-NL construct. For drug-incubation experiments 100uL of A-443654 dose-response curve (100, 10, 1, 0.1, 0.01, 0.001, 0.0001, 0.00001 μM final concentration at 0.1% DMSO) and 528 uL of 1 uM BX-795 or 528 uL of 1 μM DMSO made in Opti-MEM, followed by 256 μL of a 20x K5 tracer solution (made according to Promega’s adherent target engagement protocol) were added to plates and incubated at 37°C for 24 hrs. Cells were harvested using a solution of phosphatase and protease inhibitors (1:100 dilution) in RIPA buffer. Protein quantification was performed utilizing the Pierce BCA Protein Assay Kit protocol. Data were quantified using the preset BCA 560 nm absorbance protocol on the GloMax Discover plate reader, exported in Microsoft Excel format, and analyzed in GraphPad Prism 10. SDS-Page gel electrophoresis was performed with 10 μg of protein per well. Proteins were transferred to PVDF membranes utilizing a Trans-Blot Turbo RTA Transfer Kit and Trans-Blot Turbo Transfer System. Membranes were blocked in 5% milk in TBST at room temperature for 1 hour before overnight incubation in a primary antibody solution at 4°C on a rocking shaker. Dilutions for T308, β-Actin, and Total AKT primary antibodies were 1:1,000 in 5% milk in TBST, and S473 was 1:2,000 dilution in 5% milk in TBST. The secondary antibody was made at a 1:20,000 dilution in 2.5% BSA in TBST, and was added after washing membranes 3x in TBST. Gels were imaged using SuperSignal West Femto Substrate Kit and ChemiDoc MP Imager.

#### Calculation of fold phosphorylation change

Densitometry analysis for the MDA-MB-468 cell line in Figure 6B was performed in Bio-Rad Image Lab 6. Raw intensities were used in the following formula:

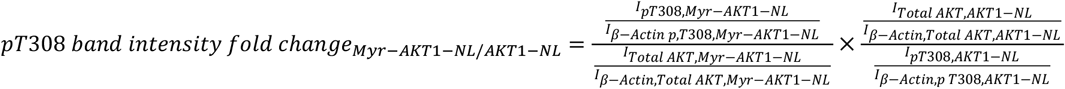

Band intensities for Myr-AKT1-NL and AKT1-NL (pT308 and Total AKT) were normalized to their corresponding β-Actin band intensities and then divided.

### MDA-MB-231 and MDA-MB-468 Cancer Cell Lines Viability Assay

Cultured MDA-MB-468 or MDA-MB-231 cells were trypsinized and transferred to clear adherent 96-well plates (5,000 cells/well, 90 µL/well) and incubated overnight in DMEM. AKT inhibitors Ipatasertib, Capivasertib, MK-2206, A443654, Miransertib, and Inhibitor VII were then added separately at final concentrations of 100, 10, 1, 0.1, 0.01, 0.001, and 0.0001 µM (0.1% DMSO final using 10 µL/well), and incubated for five days at 37°C with 5% CO_2_. Cell viability was assessed using a resazurin assay where 10 µL of resazurin (0.01mg/mL final, 0.1 mg/mL stock made in 1x PBS, pH 7.2) was added to each well and incubated at 37°C with 5% CO_2_ for 2 hrs. 96-well plates were then read on a GloxMax Discover plate reader using the CellTiter-Blue® Cell Viability Assay Protocol. For each condition, a control with DMSO (no inhibitor) was included, and viability was determined by dividing the intensity of each condition by the control. Data was exported and analyzed in Microsoft Excel and GraphPad Prism 10. Statistics were performed in Origin 2023.

## ACKNOWLEDGMENTS

The authors would like to acknowledge Dr. Natalie R. Gassman for providing MDA-MB-231 and MDA-MB-468 cell lines. The authors would also like to thank Drs. Nathaniel J. Traaseth, Arvin Dar, James Vasta, and Matthew Robers for feedback on the manuscript.

## AUTHOR CONTRIBUTIONS

Study Design: WMM. Data Collection: JWH, FAOS, ENR. Data Analysis: WMM, JWH, FAOS. Data interpretation: WMM, JWH, FAOS. Drafted Paper: WMM. Revised critically for intellectual merit: WMM, JWH, FAOS, ENR.

## REFERENCES

(1) Calleja, V.; Alcor, D.; Laguerre, M.; Park, J.; Vojnovic, B.; Hemmings, B. A.; Downward, J.; Parker, P. J.; Larijani, B. Intramolecular and Intermolecular Interactions of Protein Kinase B Define Its Activation in Vivo. PLoS Biol. 2007, 5 (4), e95.

(2) Altomare, D. A.; Tanno, S.; De Rienzo, A.; Klein-Szanto, A. J.; Tanno, S.; Skele, K. L.; Hoffman, J. P.; Testa, J. R. Frequent Activation of AKT2 Kinase in Human Pancreatic Carcinomas. J. Cell. Biochem. 2002, 87 (4), 470–476.

(3) Bellacosa, A.; de Feo, D.; Godwin, A. K.; Bell, D. W.; Cheng, J. Q.; Altomare, D. A.; Wan, M.; Dubeau, L.; Scambia, G.; Masciullo, V.; Ferrandina, G.; Benedetti Panici, P.; Mancuso, S.; Neri, G.; Testa, J. R. Molecular Alterations of the AKT2 Oncogene in Ovarian and Breast Carcinomas. Int. J. Cancer 1995, 64 (4), 280–285.

(4) Staal, S. P. Molecular Cloning of the Akt Oncogene and Its Human Homologues AKT1 and AKT2: Amplification of AKT1 in a Primary Human Gastric Adenocarcinoma. Proc. Natl. Acad. Sci. U. S. A. 1987, 84 (14), 5034–5037.

(5) She, Q.-B.; Chandarlapaty, S.; Ye, Q.; Lobo, J.; Haskell, K. M.; Leander, K. R.; DeFeo-Jones, D.; Huber, H. E.; Rosen, N. Breast Tumor Cells with PI3K Mutation or HER2 Amplification Are Selectively Addicted to Akt Signaling. PLoS One 2008, 3 (8), e3065.

(6) Yi, K. H.; Lauring, J. Recurrent AKT Mutations in Human Cancers: Functional Consequences and Effects on Drug Sensitivity. Oncotarget 2016, 7 (4), 4241–4251.

(7) Wu, W.; Chen, Y.; Huang, L.; Li, W.; Tao, C.; Shen, H. Effects of AKT1 E17K Mutation Hotspots on the Biological Behavior of Breast Cancer Cells. Int. J. Clin. Exp. Pathol. 2020, 13 (3), 332–346.

(8) Lin, K.; Lin, J.; Wu, W.-I.; Ballard, J.; Lee, B. B.; Gloor, S. L.; Vigers, G. P. A.; Morales, T. H.; Friedman, L. S.; Skelton, N.; Brandhuber, B. J. An ATP-Site on-off Switch That Restricts Phosphatase Accessibility of Akt. Sci. Signal. 2012, 5 (223), ra37.

(9) Okuzumi, T.; Fiedler, D.; Zhang, C.; Gray, D. C.; Aizenstein, B.; Hoffman, R.; Shokat, K. M. Inhibitor Hijacking of Akt Activation. Nat. Chem. Biol. 2009, 5 (7), 484–493.

(10) Ma, B. B. Y.; Goh, B. C.; Lim, W. T.; Hui, E. P.; Tan, E. H.; Lopes, G. de L.; Lo, K. W.; Li, L.; Loong, H.; Foster, N. R.; Erlichman, C.; King, A. D.; Kam, M. K. M.; Leung, S. F.; Chan, K. C.; Chan, A. T. C. Multicenter Phase II Study of the AKT Inhibitor MK-2206 in Recurrent or Metastatic Nasopharyngeal Carcinoma from Patients in the Mayo Phase II Consortium and the Cancer Therapeutics Research Group (MC1079). Invest. New Drugs 2015, 33 (4), 985–991.

(11) Shimizu, T.; Tolcher, A. W.; Papadopoulos, K. P.; Beeram, M.; Rasco, D. W.; Smith, L. S.; Gunn, S.; Smetzer, L.; Mays, T. A.; Kaiser, B.; Wick, M. J.; Alvarez, C.; Cavazos, A.; Mangold, G. L.; Patnaik, A. The Clinical Effect of the Dual-Targeting Strategy Involving PI3K/AKT/mTOR and RAS/MEK/ERK Pathways in Patients with Advanced Cancer. Clin. Cancer Res. 2012, 18 (8), 2316–2325.

(12) Robers, M. B.; Dart, M. L.; Woodroofe, C. C.; Zimprich, C. A.; Kirkland, T. A.; Machleidt, T.; Kupcho, K. R.; Levin, S.; Hartnett, J. R.; Zimmerman, K.; Niles, A. L.; Ohana, R. F.; Daniels, D. L.; Slater, M.; Wood, M. G.; Cong, M.; Cheng, Y.-Q.; Wood, K. V. Target Engagement and Drug Residence Time Can Be Observed in Living Cells with BRET. Nat. Commun. 2015, 6 (1), 10091.

(13) Wells, C. I.; Vasta, J. D.; Corona, C. R.; Wilkinson, J.; Zimprich, C. A.; Ingold, M. R.; Pickett, J. E.; Drewry, D. H.; Pugh, K. M.; Schwinn, M. K.; Hwang, B. B.; Zegzouti, H.; Huber, K. V. M.; Cong, M.; Meisenheimer, P. L.; Willson, T. M.; Robers, M. B. Quantifying CDK Inhibitor Selectivity in Live Cells. Nat. Commun. 2020, 11 (1), 2743.

(14) Stottrup, C.; Tsang, T.; Chin, Y. R. Upregulation of AKT3 Confers Resistance to the AKT Inhibitor MK2206 in Breast Cancer. Mol Cancer Ther 2016, 15 (8), 1964–1974.

(15) Yan, L. Abstract #DDT01-1: MK-2206: A Potent Oral Allosteric AKT Inhibitor. Cancer Research 2009, 69 (9_Supplement), DDT01–1.

(16) Quambusch, L.; Landel, I.; Depta, L.; Weisner, J.; Uhlenbrock, N.; Müller, M. P.; Glanemann, F.; Althoff, K.; Siveke, J. T.; Rauh, D. Covalent-Allosteric Inhibitors to Achieve Akt Isoform-Selectivity. Angew. Chem. Int. Ed Engl. 2019, 58 (52), 18823–18829.

(17) Quambusch, L.; Depta, L.; Landel, I.; Lubeck, M.; Kirschner, T.; Nabert, J.; Uhlenbrock, N.; Weisner, J.; Kostka, M.; Levy, L. M.; Schultz-Fademrecht, C.; Glanemann, F.; Althoff, K.; Müller, M. P.; Siveke, J. T.; Rauh, D. Cellular Model System to Dissect the Isoform-Selectivity of Akt Inhibitors. Nat. Commun. 2021, 12 (1), 5297.

(18) Yu, Y.; Savage, R. E.; Eathiraj, S.; Meade, J.; Wick, M. J.; Hall, T.; Abbadessa, G.; Schwartz, B. Targeting AKT1-E17K and the PI3K/AKT Pathway with an Allosteric AKT Inhibitor, ARQ 092. PLoS One 2015, 10 (10), e0140479.

(19) Parikh, C.; Janakiraman, V.; Wu, W.-I.; Foo, C. K.; Kljavin, N. M.; Chaudhuri, S.; Stawiski, E.; Lee, B.; Lin, J.; Li, H.; Lorenzo, M. N.; Yuan, W.; Guillory, J.; Jackson, M.; Rondon, J.; Franke, Y.; Bowman, K. K.; Sagolla, M.; Stinson, J.; Wu, T. D.; Wu, J.; Stokoe, D.; Stern, H. M.; Brandhuber, B. J.; Lin, K.; Skelton, N. J.; Seshagiri, S. Disruption of PH-Kinase Domain Interactions Leads to Oncogenic Activation of AKT in Human Cancers. Proc. Natl. Acad. Sci. U. S. A. 2012, 109 (47), 19368–19373.

(20) Carpten, J. D.; Faber, A. L.; Horn, C.; Donoho, G. P.; Briggs, S. L.; Robbins, C. M.; Hostetter, G.; Boguslawski, S.; Moses, T. Y.; Savage, S.; Uhlik, M.; Lin, A.; Du, J.; Qian, Y.-W.; Zeckner, D. J.; Tucker-Kellogg, G.; Touchman, J.; Patel, K.; Mousses, S.; Bittner, M.; Schevitz, R.; Lai, M.-H. T.; Blanchard, K. L.; Thomas, J. E. A Transforming Mutation in the Pleckstrin Homology Domain of AKT1 in Cancer. Nature 2007, 448 (7152), 439–444.

(21) Herberts, C.; Murtha, A. J.; Fu, S.; Wang, G.; Schönlau, E.; Xue, H.; Lin, D.; Gleave, A.; Yip, S.; Angeles, A.; Hotte, S.; Tran, B.; North, S.; Taavitsainen, S.; Beja, K.; Vandekerkhove, G.; Ritch, E.; Warner, E.; Saad, F.; Iqbal, N.; Nykter, M.; Gleave, M. E.; Wang, Y.; Annala, M.; Chi, K. N.; Wyatt, A. W. Activating AKT1 and PIK3CA Mutations in Metastatic Castration-Resistant Prostate Cancer. Eur. Urol. 2020, 78 (6), 834–844.

(22) Davies, M. A.; Stemke-Hale, K.; Tellez, C.; Calderone, T. L.; Deng, W.; Prieto, V. G.; Lazar, A. J. F.; Gershenwald, J. E.; Mills, G. B. A Novel AKT3 Mutation in Melanoma Tumours and Cell Lines. Br. J. Cancer 2008, 99 (8), 1265–1268.

(23) Wang, J.; Zhao, W.; Guo, H.; Fang, Y.; Stockman, S. E.; Bai, S.; Ng, P. K.-S.; Li, Y.; Yu, Q.; Lu, Y.; Jeong, K. J.; Chen, X.; Gao, M.; Liang, J.; Li, W.; Tian, X.; Jonasch, E.; Mills, G. B.; Ding, Z. AKT Isoform-Specific Expression and Activation across Cancer Lineages. BMC Cancer 2018, 18 (1), 742.

(24) Shrestha Bhattarai, T.; Shamu, T.; Gorelick, A. N.; Chang, M. T.; Chakravarty, D.; Gavrila, E. I.; Donoghue, M. T. A.; Gao, J.; Patel, S.; Gao, S. P.; Reynolds, M. H.; Phillips, S. M.; Soumerai, T.; Abida, W.; Hyman, D. M.; Schram, A. M.; Solit, D. B.; Smyth, L. M.; Taylor, B. S. AKT Mutant Allele-Specific Activation Dictates Pharmacologic Sensitivities. Nat. Commun. 2022, 13 (1), 2111.

(25) Ebner, M.; Lučić, I.; Leonard, T. A.; Yudushkin, I. PI(3,4,5)P3 Engagement Restricts Akt Activity to Cellular Membranes. Mol. Cell 2017, 65 (3), 416–431.e6.

(26) Kohn, A. D.; Takeuchi, F.; Roth, R. A. Akt, a Pleckstrin Homology Domain Containing Kinase, Is Activated Primarily by Phosphorylation. J. Biol. Chem. 1996, 271 (36), 21920–21926.

(27) Andjelković, M.; Alessi, D. R.; Meier, R.; Fernandez, A.; Lamb, N. J.; Frech, M.; Cron, P.; Cohen, P.; Lucocq, J. M.; Hemmings, B. A. Role of Translocation in the Activation and Function of Protein Kinase B. J. Biol. Chem. 1997, 272 (50), 31515–31524.

(28) Chan, T. O.; Zhang, J.; Rodeck, U.; Pascal, J. M.; Armen, R. S.; Spring, M.; Dumitru, C. D.; Myers, V.; Li, X.; Cheung, J. Y.; Feldman, A. M. Resistance of Akt Kinases to Dephosphorylation through ATP-Dependent Conformational Plasticity. Proc. Natl. Acad. Sci. U. S. A. 2011, 108 (46), E1120–E1127.

(29) Lehmann, B. D.; Bauer, J. A.; Chen, X.; Sanders, M. E.; Chakravarthy, A. B.; Shyr, Y.; Pietenpol, J. A. Identification of Human Triple-Negative Breast Cancer Subtypes and Preclinical Models for Selection of Targeted Therapies. J. Clin. Invest. 2011, 121 (7), 2750–2767.

(30) Yam, C.; Mani, S. A.; Moulder, S. L. Targeting the Molecular Subtypes of Triple Negative Breast Cancer: Understanding the Diversity to Progress the Field. Oncologist 2017, 22 (9), 1086–1093.

(31) van der Noord, V. E.; McLaughlin, R. P.; Smid, M.; Foekens, J. A.; Martens, J. W. M.; Zhang, Y.; van de Water, B. An Increased Cell Cycle Gene Network Determines MEK and Akt Inhibitor Double Resistance in Triple-Negative Breast Cancer. Sci. Rep. 2019, 9 (1), 13308.

(32) Yang, W.; Ju, J.-H.; Lee, K.-M.; Shin, I. Akt Isoform-Specific Inhibition of MDA-MB-231 Cell Proliferation. Cell. Signal. 2011, 23 (1), 19–26.

