## Supplementary Information for "Live-cell NanoBRET assay to measure AKT inhibitor binding to conformational states of AKT"

### Table of Contents


| Figure | Page |
|---|---|
| Figure S1 | 2-4 |
| Figure S2 | 5-7 |
| Figure S3 | 8-10 |
| Figure S4 | 11-20 |
| Figure S5 | 21 |
| Figure S6 | 22 |
| Figure S7 | 23-26 |
| Figure S8 | 27-28 |
| Figure S9 | 39 |
| Figure S10 | 30-35 |

A

### AKT-NL

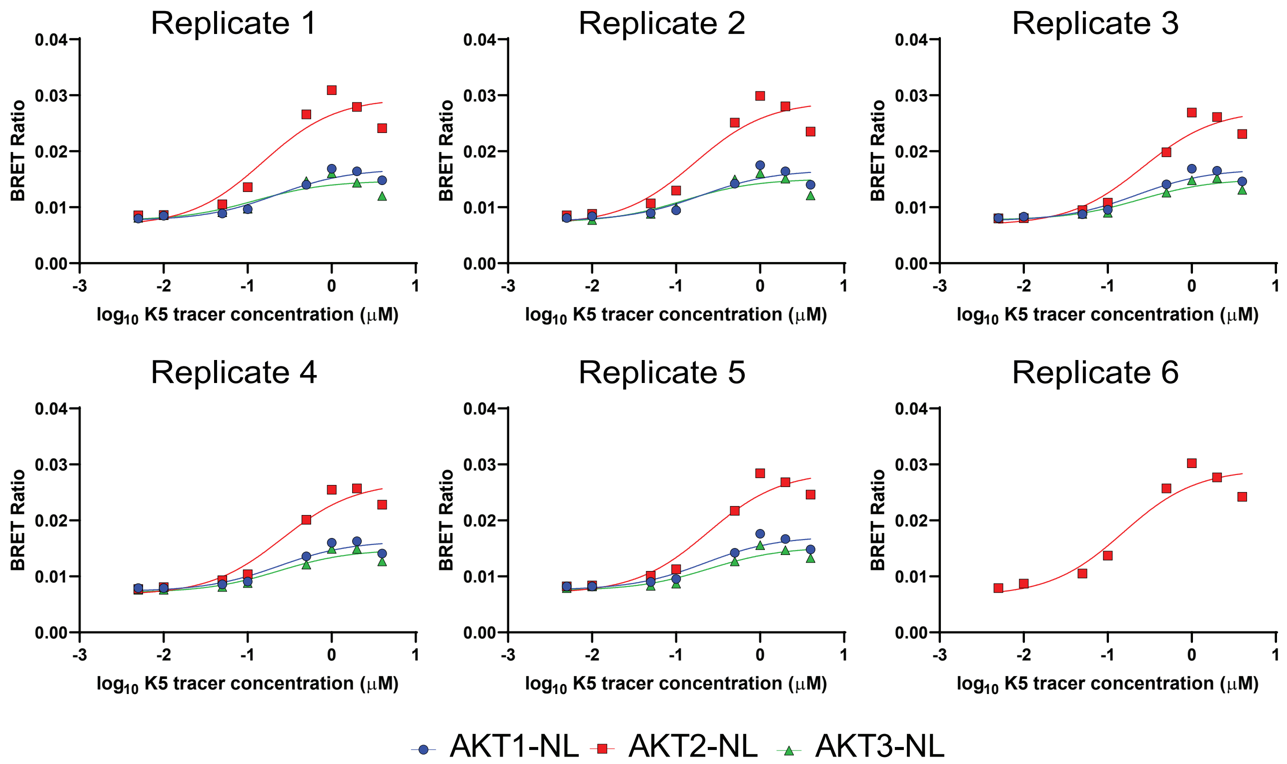


| K5 $EC_{50}$ | Replicate 1 (μM) | Replicate 2 (μM) | Replicate 3 (μM) | Replicate 4 (μM) | Replicate 5 (μM) | Replicate 6 (μM) | Average (μM) | Standard Error |
|---|---|---|---|---|---|---|---|---|
| AKT1-NL | 0.2232 | 0.1977 | 0.2204 | 0.2331 | 0.2179 | | 0.2185 | 0.0058 |
| AKT2-NL | 0.1525 | 0.1666 | 0.2598 | 0.2591 | 0.2395 | 0.1527 | 0.2050 | 0.0217 |
| AKT3-NL | 0.1219 | 0.1215 | 0.2444 | 0.2320 | 0.2460 | | 0.1932 | 0.0293 |

### NL-AKT

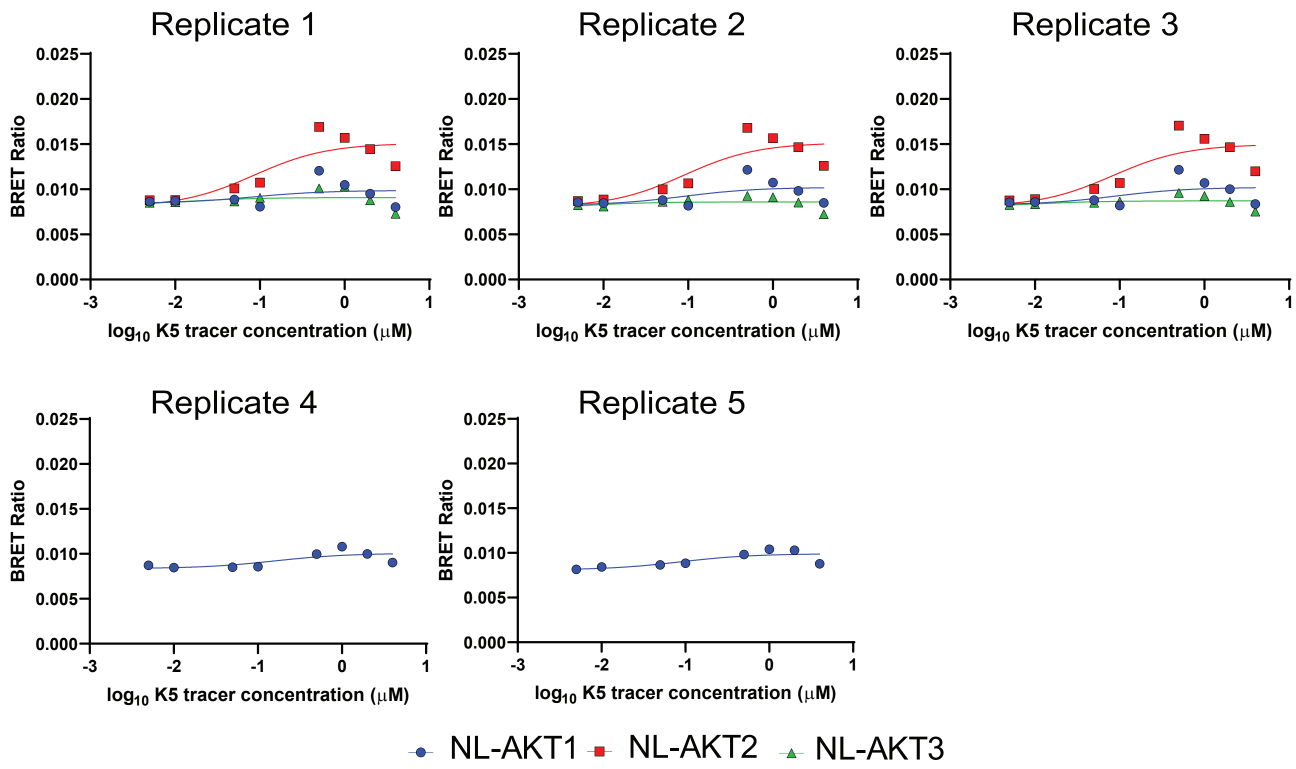


| K5 $EC_{50}$ | Replicate 1 (μM) | Replicate 2 (μM) | Replicate 3 (μM) | Replicate 4 (μM) | Replicate 5 (μM) | Average (μM) | Standard Error |
|---|---|---|---|---|---|---|---|
| NL-AKT1 | 0.0819 | 0.0883 | 0.0890 | 0.1580 | 0.0867 | 0.1008 | 0.0144 |
| NL-AKT2 | 0.0853 | 0.0903 | 0.0830 | | | 0.0862 | 0.0022 |
| NL-AKT3 | 0.0136 | 0.0042 | 0.0079 | | | 0.0085 | 0.0027 |

B

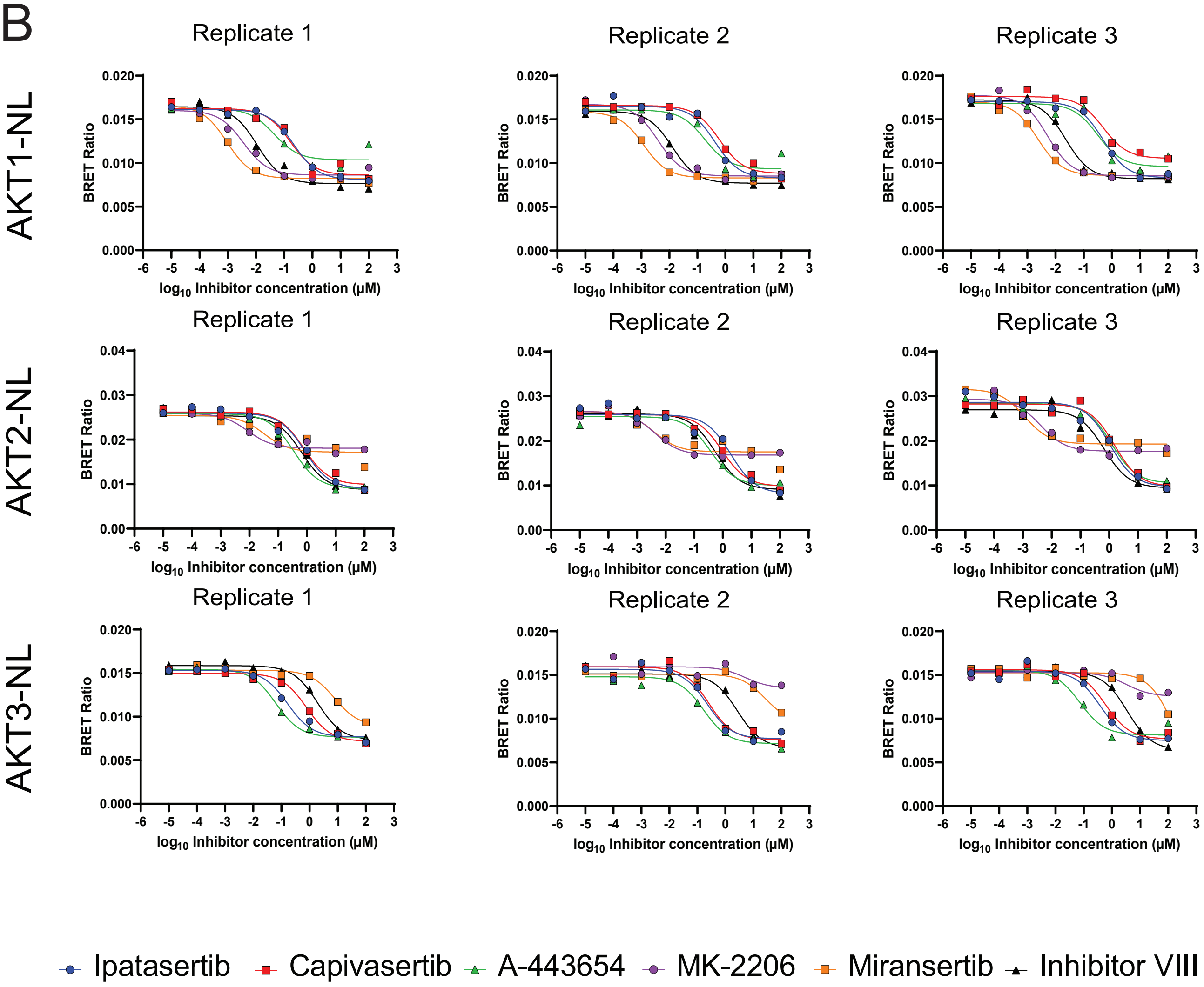


C

Apparent $IC_{50}$ values

| AKT1-NL $IC_{50}$ | Replicate 1 (μM) | Replicate 2 (μM) | Replicate 3 (μM) | Average (μM) | Standard Error |
|---|---|---|---|---|---|
| Ipatasertib | 0.2220 | 0.4338 | 0.4931 | 0.3830 | 0.0823 |
| Capivasertib | 0.1661 | 0.6136 | 0.4913 | 0.4237 | 0.1335 |
| A-443654 | 0.0445 | 0.1863 | 0.3170 | 0.1826 | 0.0787 |
| MK-2206 | 0.0036 | 0.0039 | 0.0052 | 0.0042 | 0.0005 |
| Miransertib | 0.0009 | 0.0012 | 0.0023 | 0.0015 | 0.0004 |
| Inhibitor VIII | 0.0112 | 0.0129 | 0.0232 | 0.0158 | 0.0038 |

| AKT2-NL $IC_{50}$ | Replicate 1 (μM) | Replicate 2 (μM) | Replicate 3 (μM) | Average (μM) | Standard Error |
|---|---|---|---|---|---|
| Ipatasertib | 0.8388 | 1.9130 | 1.0600 | 1.2706 | 0.3275 |
| Capivasertib | 0.7908 | 0.9428 | 1.5450 | 1.0929 | 0.2303 |
| A-443654 | 0.3491 | 0.3740 | 1.1840 | 0.6357 | 0.2742 |
| MK-2206 | 0.0087 | 0.0043 | 0.0041 | 0.0057 | 0.0015 |
| Miransertib | 0.0325 | 0.0039 | 0.0011 | 0.0125 | 0.0100 |
| Inhibitor VIII | 0.6635 | 0.5476 | 0.6634 | 0.6248 | 0.0386 |

| AKT3-NL $IC_{50}$ | Replicate 1 (μM) | Replicate 2 (μM) | Replicate 3 (μM) | Average (μM) | Standard Error |
|---|---|---|---|---|---|
| Ipatasertib | 0.1660 | 0.2252 | 0.3850 | 0.2587 | 0.0654 |
| Capivasertib | 0.7786 | 0.2567 | 0.5909 | 0.5421 | 0.1526 |
| A-443654 | 0.0634 | 0.1698 | 0.0746 | 0.1026 | 0.0338 |
| MK-2206 | N/A | N/A | N/A | N/A | N/A |
| Miransertib | 7.81 | 26.78 | 154.90 | 63.16 | 46.19 |
| Inhibitor VIII | 1.94 | 2.71 | 3.48 | 2.71 | 0.45 |

D

Calculated $K_i$ values

| AKT1-NL $K_i$ | Replicate 1 (nM) | Replicate 2 (nM) | Replicate 3 (nM) | Average (nM) | Standard Error |
|---|---|---|---|---|---|
| Ipatasertib | 39.80 | 77.77 | 88.41 | 68.66 | 14.75 |
| Capivasertib | 29.78 | 110.01 | 88.09 | 75.96 | 23.94 |
| A-443654 | 7.98 | 33.40 | 56.84 | 32.74 | 14.11 |
| MK-2206 | 0.6433 | 0.6991 | 0.9264 | 0.7563 | 0.0866 |
| Miransertib | 0.1685 | 0.2152 | 0.4195 | 0.2677 | 0.0771 |
| Inhibitor VIII | 2.00 | 2.32 | 4.16 | 2.83 | 0.67 |

| AKT2-NL $K_i$ | Replicate 1 (nM) | Replicate 2 (nM) | Replicate 3 (nM) | Average (nM) | Standard Error |
|---|---|---|---|---|---|
| Ipatasertib | 142.72 | 325.49 | 180.36 | 216.19 | 55.72 |
| Capivasertib | 134.55 | 160.42 | 262.88 | 185.95 | 39.18 |
| A-443654 | 59.40 | 63.64 | 201.45 | 108.16 | 46.66 |
| MK-2206 | 1.47 | 0.73 | 0.69 | 0.97 | 0.25 |
| Miransertib | 5.53 | 0.66 | 0.20 | 2.13 | 1.71 |
| Inhibitor VIII | 112.89 | 93.17 | 112.88 | 106.31 | 6.57 |

| AKT3-NL $K_i$ | Replicate 1 (nM) | Replicate 2 (nM) | Replicate 3 (nM) | Average (nM) | Standard Error |
|---|---|---|---|---|---|
| Ipatasertib | 26.87 | 36.46 | 62.33 | 41.89 | 10.59 |
| Capivasertib | 126.05 | 41.56 | 95.66 | 87.75 | 24.71 |
| A-443654 | 10.26 | 27.49 | 12.08 | 16.61 | 5.46 |
| MK-2206 | N/A | N/A | N/A | N/A | N/A |
| Miransertib | 1264.36 | 4335.40 | 25076.67 | 10225.48 | 7478.33 |
| Inhibitor VIII | 313.42 | 438.72 | 564.02 | 438.72 | 72.34 |

**Figure S1**-Assay optimization for NanoLuciferase placement, and AKT inhibitor binding curve replicate data. **(A)** Signal buildup curves and fitted $EC_{50}$ values using the K5 tracer on 293T cells transfected with AKT-NL or NL-AKT. **(B)** Binding curves for AKT inhibitors on AKT1-NL, AKT2-NL, and AKT3-NL. **(C)**Fitted apparent $IC_{50}$ values for curves in (B). **(D)** Calculated $K_i$ values derived from $EC_{50}$ and $IC_{50}$ values in (A) and (C).

A

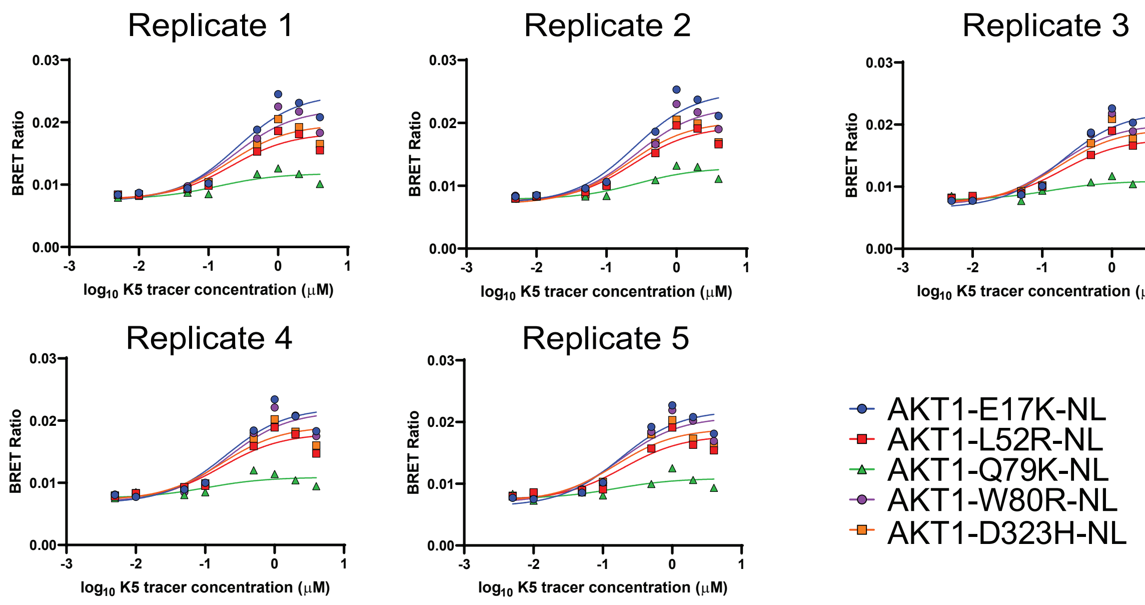


| K5 $EC_{50}$ | Replicate 1 (μM) | Replicate 2 (μM) | Replicate 3 (μM) | Replicate 4 (μM) | Replicate 5 (μM) | Average (μM) | Standard Error |
|---|---|---|---|---|---|---|---|
| AKT1-E17K-NL | 0.2542 | 0.2503 | 0.1959 | 0.1958 | 0.1777 | 0.2148 | 0.0157 |
| AKT1-L52R-NL | 0.2103 | 0.2384 | 0.1890 | 0.1741 | 0.1983 | 0.2020 | 0.0109 |
| AKT1-Q79K-NL | 0.1340 | 0.2766 | 0.1313 | 0.1043 | 0.1534 | 0.1599 | 0.0302 |
| AKT1-W80R-NL | 0.2318 | 0.2607 | 0.1640 | 0.2160 | 0.1763 | 0.2098 | 0.0178 |
| AKT1-D323H-NL | 0.1965 | 0.2087 | 0.1723 | 0.1724 | 0.1532 | 0.1806 | 0.0098 |

B

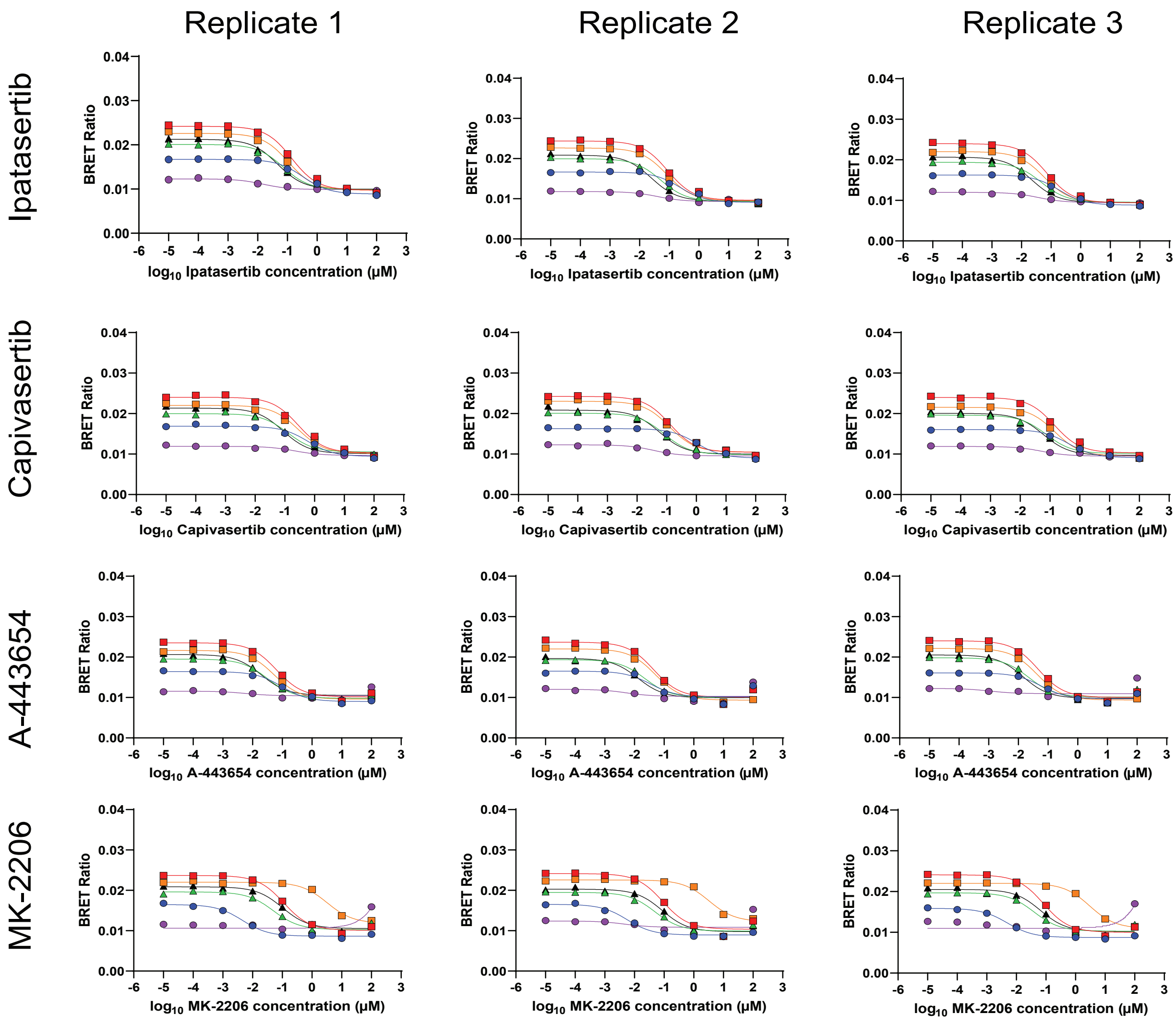

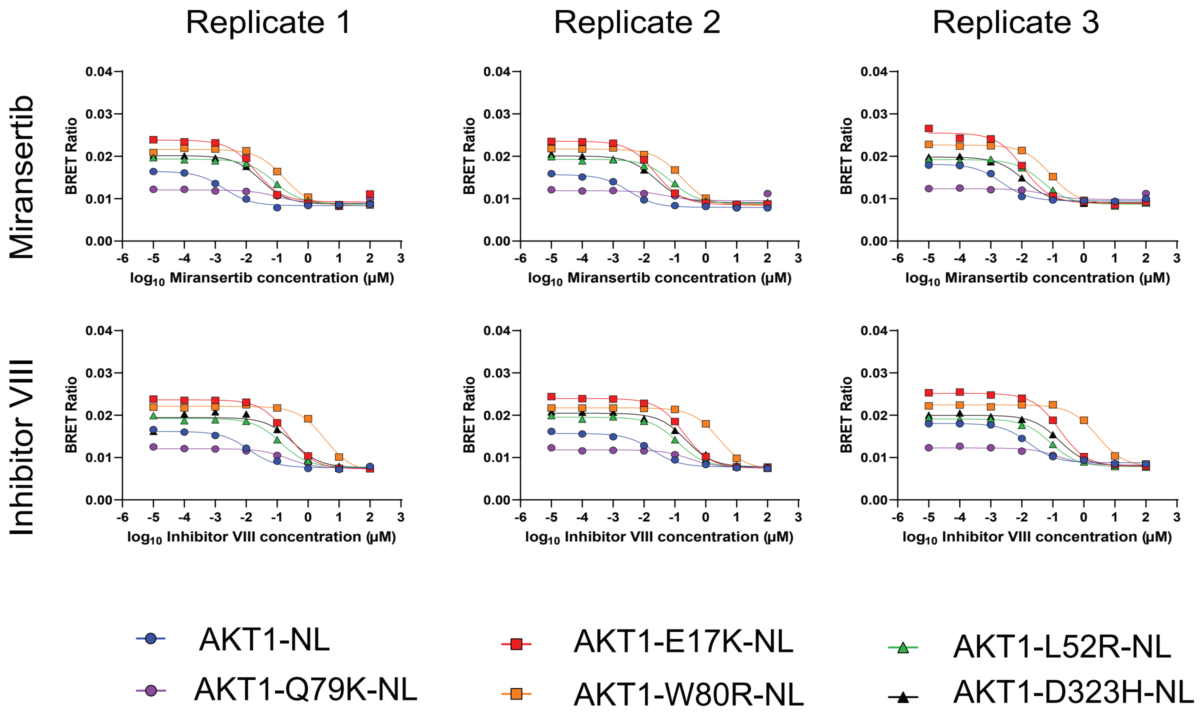


# C

#### Apparent $IC_{50}$ values

| | Replicate 1 (µM) | Replicate 2 (µM) | Replicate 3 (µM) | Average (µM) | Standard Error |
|---|---|---|---|---|---|
| WT-AKT1-NL | 0.3331 | 0.2278 | 0.1880 | 0.2496 | 0.0433 |
| AKT1-E17K-NL | 0.1384 | 0.0883 | 0.0724 | 0.0997 | 0.0199 |
| AKT1-L52R-NL | 0.0719 | 0.0515 | 0.0466 | 0.0567 | 0.0077 |
| AKT1-Q79K-NL | 0.0155 | 0.0402 | 0.0356 | 0.0304 | 0.0076 |
| AKT1-W80R-NL | 0.1060 | 0.0829 | 0.0586 | 0.0825 | 0.0137 |
| AKT1-D323H-NL | 0.0474 | 0.0252 | 0.0222 | 0.0316 | 0.0079 |

| | Replicate 1 (µM) | Replicate 2 (µM) | Replicate 3 (µM) | Average (µM) | Standard Error |
|---|---|---|---|---|---|
| WT-AKT1-NL | 0.5989 | 1.0380 | 0.4196 | 0.6855 | 0.1837 |
| AKT1-E17K-NL | 0.2811 | 0.1305 | 0.1409 | 0.1842 | 0.0486 |
| AKT1-L52R-NL | 0.1251 | 0.0820 | 0.0740 | 0.0937 | 0.0159 |
| AKT1-Q79K-NL | 0.2128 | 0.0359 | 0.0529 | 0.1005 | 0.0563 |
| AKT1-W80R-NL | 0.2812 | 0.1243 | 0.1444 | 0.1833 | 0.0493 |
| AKT1-D323H-NL | 0.0867 | 0.0569 | 0.0589 | 0.0675 | 0.0096 |

| | Replicate 1 (µM) | Replicate 2 (µM) | Replicate 3 (µM) | Average (µM) | Standard Error |
|---|---|---|---|---|---|
| WT-AKT1-NL | 0.0965 | 0.0349 | 0.0580 | 0.0631 | 0.0180 |
| AKT1-E17K-NL | 0.0637 | 0.0447 | 0.0462 | 0.0515 | 0.0061 |
| AKT1-L52R-NL | 0.0321 | 0.0226 | 0.0224 | 0.0257 | 0.0032 |
| AKT1-Q79K-NL | 0.0058 | 0.0052 | 0.0011 | 0.0041 | 0.0015 |
| AKT1-W80R-NL | 0.0531 | 0.0502 | 0.0424 | 0.0486 | 0.0032 |
| AKT1-D323H-NL | 0.0199 | 0.0130 | 0.0143 | 0.0157 | 0.0021 |

| | Replicate 1 (µM) | Replicate 2 (µM) | Replicate 3 (µM) | Average (µM) | Standard Error |
|---|---|---|---|---|---|
| WT-AKT1-NL | 0.0046 | 0.0057 | 0.0049 | 0.0050 | 0.0003 |
| AKT1-E17K-NL | 0.1125 | 0.0896 | 0.0821 | 0.0947 | 0.0091 |
| AKT1-L52R-NL | 0.0502 | 0.0489 | 0.0397 | 0.0463 | 0.0033 |
| AKT1-Q79K-NL | N/A | 0.0048 | N/A | 0.0048 | N/A |
| AKT1-W80R-NL | 3.2430 | 3.2500 | 2.9920 | 3.1617 | 0.0849 |
| AKT1-D323H-NL | 0.1013 | 0.0863 | 0.0656 | 0.0844 | 0.0103 |

| | Replicate 1 (µM) | Replicate 2 (µM) | Replicate 3 (µM) | Average (µM) | Standard Error |
|---|---|---|---|---|---|
| WT-AKT1-NL | 0.0019 | 0.0032 | 0.0023 | 0.0025 | 0.0004 |
| AKT1-E17K-NL | 0.0199 | 0.0236 | 0.0113 | 0.0183 | 0.0036 |
| AKT1-L52R-NL | 0.0798 | 0.0776 | 0.0461 | 0.0678 | 0.0109 |
| AKT1-Q79K-NL | 0.1000 | 0.0534 | 0.0580 | 0.0705 | 0.0148 |
| AKT1-W80R-NL | 0.1638 | 0.1601 | 0.0852 | 0.1364 | 0.0256 |
| AKT1-D323H-NL | 0.0292 | 0.0244 | 0.0117 | 0.0218 | 0.0052 |

| | Replicate 1 (µM) | Replicate 2 (µM) | Replicate 3 (µM) | Average (µM) | Standard Error |
|---|---|---|---|---|---|
| WT-AKT1-NL | 0.0122 | 0.0189 | 0.0169 | 0.0160 | 0.0020 |
| AKT1-E17K-NL | 0.2102 | 0.1978 | 0.1647 | 0.1909 | 0.0136 |
| AKT1-L52R-NL | 0.1332 | 0.1232 | 0.0849 | 0.1138 | 0.0147 |
| AKT1-Q79K-NL | 0.2213 | 0.2867 | 0.1492 | 0.2191 | 0.0397 |
| AKT1-W80R-NL | 3.4990 | 2.6190 | 2.7090 | 2.9423 | 0.2795 |
| AKT1-D323H-NL | 0.3305 | 0.2285 | 0.1628 | 0.2406 | 0.0488 |

D

Calculated $K_i$ values

| | Replicate 1 (nM) | Replicate 2 (nM) | Replicate 3 (nM) | Average (nM) | Standard Error |
|---|---|---|---|---|---|
| WT-AKT1-NL | 59.7221 | 40.8427 | 33.7069 | 44.7572 | 7.7608 |
| AKT1-E17K-NL | 24.4699 | 15.6137 | 12.8078 | 17.6305 | 3.5143 |
| AKT1-L52R-NL | 12.0807 | 8.6555 | 7.8386 | 9.5249 | 1.2994 |
| AKT1-Q79K-NL | 2.1370 | 5.5438 | 4.9110 | 4.1973 | 1.0462 |
| AKT1-W80R-NL | 18.3793 | 14.3723 | 10.1641 | 14.3052 | 2.3718 |
| AKT1-D323H-NL | 7.2455 | 3.8599 | 3.3917 | 4.8324 | 1.2141 |

| | Replicate 1 (nM) | Replicate 2 (nM) | Replicate 3 (nM) | Average (nM) | Standard Error |
|---|---|---|---|---|---|
| WT-AKT1-NL | 107.3779 | 186.1050 | 75.2309 | 122.9046 | 32.9347 |
| AKT1-E17K-NL | 49.7001 | 23.0731 | 24.9119 | 32.5617 | 8.5856 |
| AKT1-L52R-NL | 21.0252 | 13.7882 | 12.4370 | 15.7501 | 2.6662 |
| AKT1-Q79K-NL | 29.3391 | 4.9482 | 7.2962 | 13.8611 | 7.7686 |
| AKT1-W80R-NL | 48.7572 | 21.5524 | 25.0375 | 31.7823 | 8.5468 |
| AKT1-D323H-NL | 13.2610 | 8.7080 | 9.0171 | 10.3287 | 1.4688 |

| | Replicate 1 (nM) | Replicate 2 (nM) | Replicate 3 (nM) | Average (nM) | Standard Error |
|---|---|---|---|---|---|
| WT-AKT1-NL | 17.3017 | 6.2573 | 10.4043 | 11.3211 | 3.2210 |
| AKT1-E17K-NL | 11.2608 | 7.9085 | 8.1720 | 9.1137 | 1.0762 |
| AKT1-L52R-NL | 5.4017 | 3.8034 | 3.7580 | 4.3210 | 0.5405 |
| AKT1-Q79K-NL | 0.8057 | 0.7224 | 0.1517 | 0.5599 | 0.2056 |
| AKT1-W80R-NL | 9.2035 | 8.7059 | 7.3535 | 8.4210 | 0.5527 |
| AKT1-D323H-NL | 3.0368 | 1.9873 | 2.1847 | 2.4029 | 0.3220 |

| | Replicate 1 (nM) | Replicate 2 (nM) | Replicate 3 (nM) | Average (nM) | Standard Error |
|---|---|---|---|---|---|
| WT-AKT1-NL | 0.8188 | 1.0146 | 0.8819 | 0.9051 | 0.0577 |
| AKT1-E17K-NL | 19.8906 | 15.8383 | 14.5193 | 16.7494 | 1.6161 |
| AKT1-L52R-NL | 8.4386 | 8.2134 | 6.6723 | 7.7748 | 0.5551 |
| AKT1-Q79K-NL | N/A | 0.6627 | N/A | 0.6627 | N/A |
| AKT1-W80R-NL | 562.3030 | 563.5167 | 518.7822 | 548.2006 | 14.7134 |
| AKT1-D323H-NL | 15.4976 | 13.1982 | 10.0390 | 12.9116 | 1.5823 |

| | Replicate 1 (nM) | Replicate 2 (nM) | Replicate 3 (nM) | Average (nM) | Standard Error |
|---|---|---|---|---|---|
| WT-AKT1-NL | 0.3485 | 0.5786 | 0.4052 | 0.4441 | 0.0692 |
| AKT1-E17K-NL | 3.5202 | 4.1762 | 2.0032 | 3.2332 | 0.6435 |
| AKT1-L52R-NL | 13.4185 | 13.0370 | 7.7462 | 11.4006 | 1.8305 |
| AKT1-Q79K-NL | 13.7816 | 7.3665 | 7.9966 | 9.7149 | 2.0415 |
| AKT1-W80R-NL | 28.4012 | 27.7597 | 14.7693 | 23.6434 | 4.4409 |
| AKT1-D323H-NL | 4.4734 | 3.7390 | 1.7930 | 3.3351 | 0.7997 |

| | Replicate 1 (nM) | Replicate 2 (nM) | Replicate 3 (nM) | Average (nM) | Standard Error |
|---|---|---|---|---|---|
| WT-AKT1-NL | 2.1802 | 3.3886 | 3.0211 | 2.8633 | 0.3577 |
| AKT1-E17K-NL | 37.1646 | 34.9722 | 29.1199 | 33.7522 | 2.4011 |
| AKT1-L52R-NL | 22.3865 | 20.7059 | 14.2723 | 19.1216 | 2.4727 |
| AKT1-Q79K-NL | 30.5110 | 39.5278 | 20.5704 | 30.2031 | 5.4747 |
| AKT1-W80R-NL | 606.6908 | 454.1078 | 469.7129 | 510.1705 | 48.4699 |
| AKT1-D323H-NL | 50.5623 | 34.9576 | 24.9064 | 36.8088 | 7.4639 |

**Figure S2**- Replicate data for Figure 3. **(A)** Signal buildup curves and fitted $EC_{50}$ values using the K5 tracer on 293T cells transfected with AKT1-NL mutants. **(B)** Replicate binding curves for AKT inhibitors on AKT1-NL mutants. **(C)** Fitted apparent $IC_{50}$ values for curves in (B). **(D)** Calculated $K_i$ values derived from $EC_{50}$ and $IC_{50}$ values in (A and FIgure S1A) and (C).

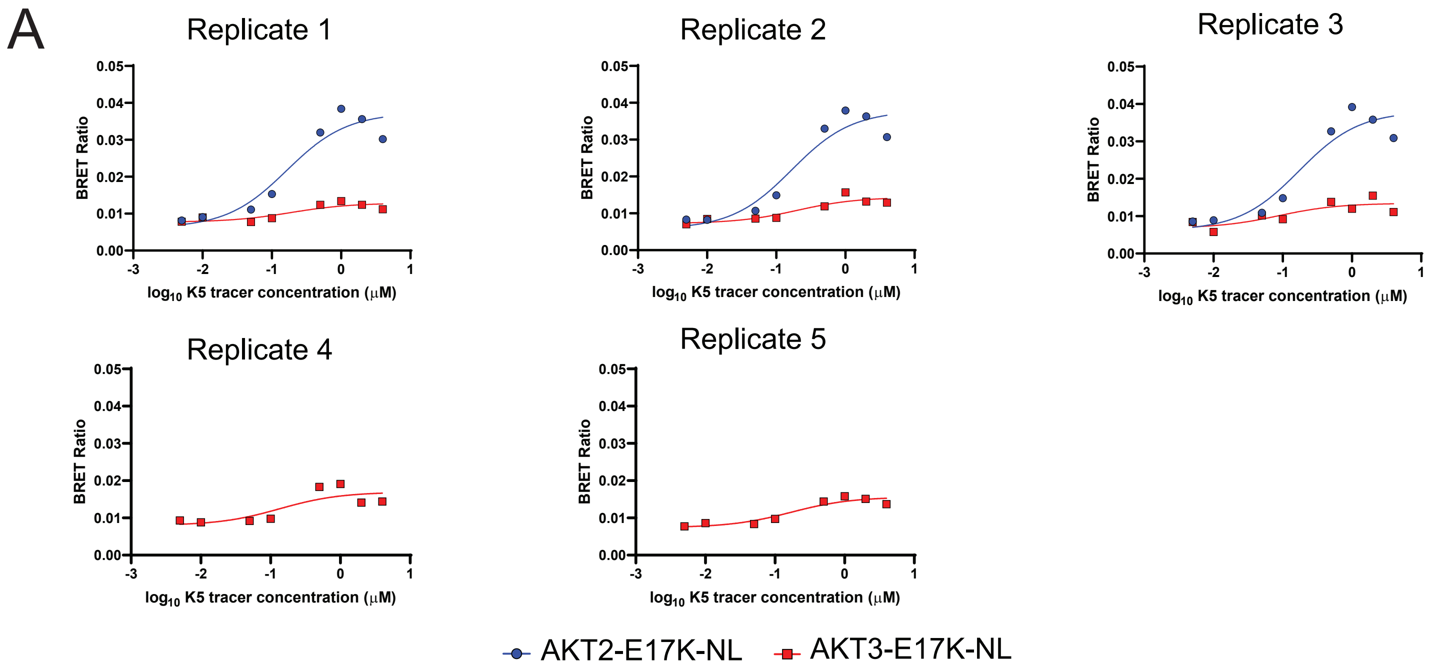


| EC50 | Replicate 1 (μM) | Replicate 2 (μM) | Replicate 3 (μM) | Replicate 4 (μM) | Replicate 5 (μM) | Average (μM) | Standard Error |
|---|---|---|---|---|---|---|---|
| AKT2-E17K-NL | 0.1643 | 0.1671 | 0.1721 | | | 0.1678 | 0.0023 |
| AKT3-E17K-NL | 0.1348 | 0.1698 | 0.1865 | 0.1997 | 0.0873 | 0.1556 | 0.0203 |

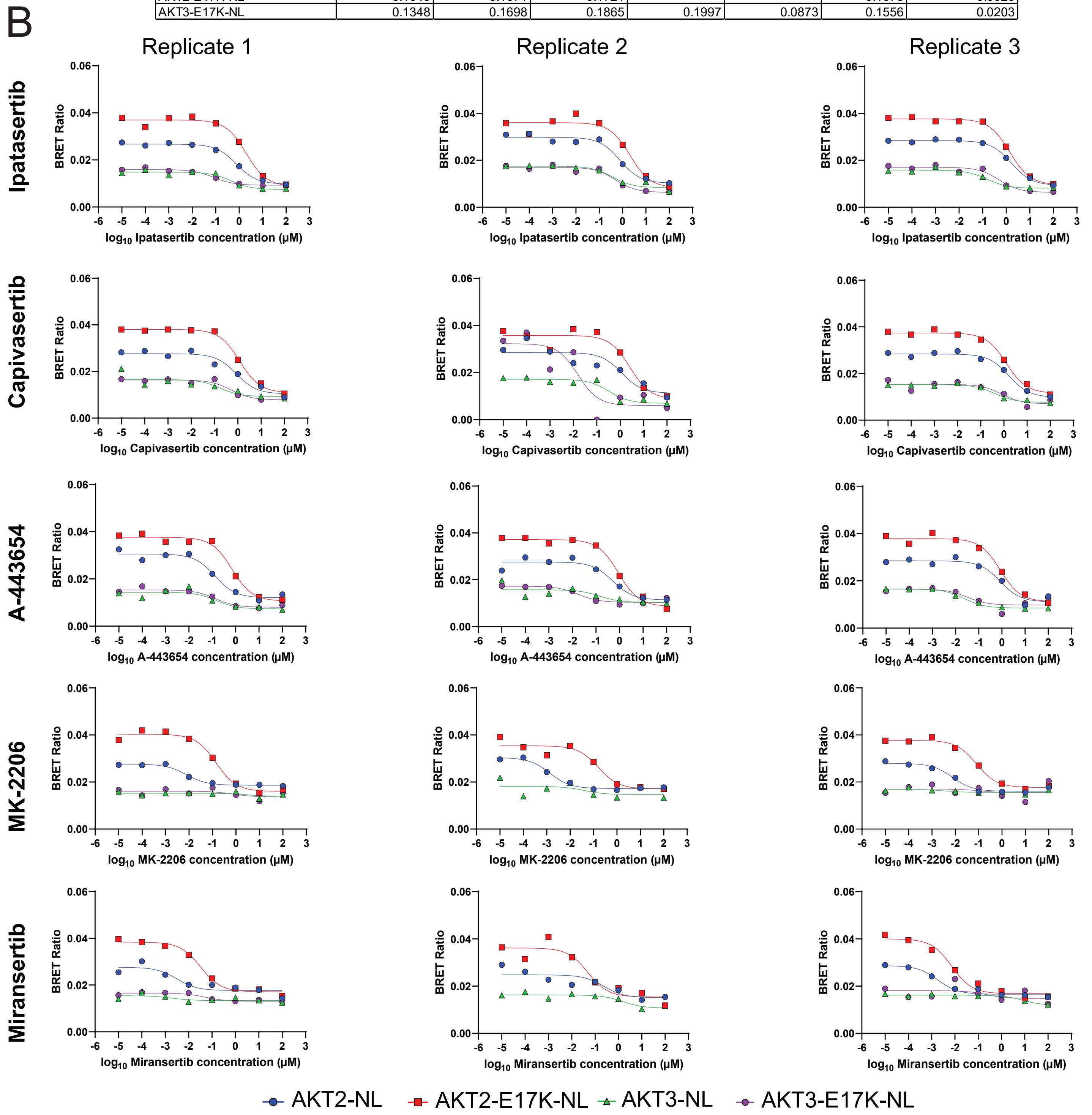

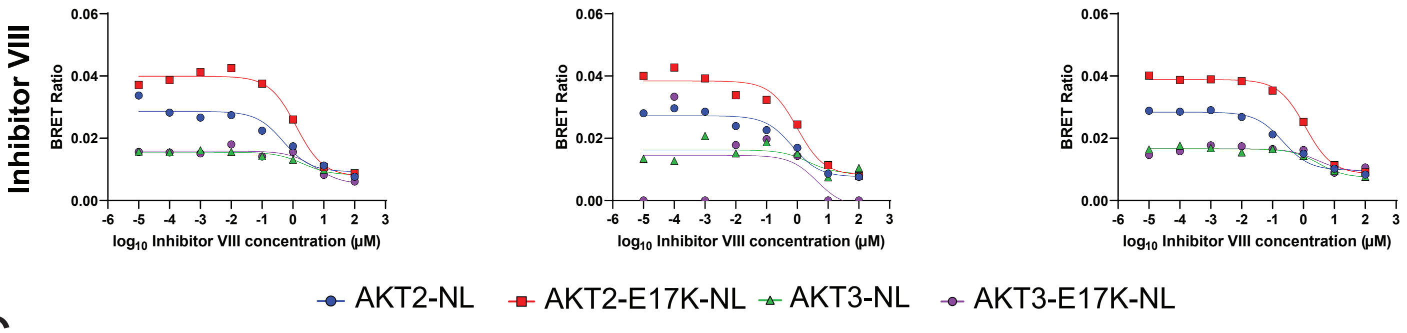


C

| Ipatasertib | Replicate 1 (μM) | Replicate 2 (μM) | Replicate 3 (μM) | Average (μM) | Standard Error |
|---|---|---|---|---|---|
| AKT2-NL | 0.7773 | 0.7825 | 1.6560 | 1.0719 | 0.2920 |
| AKT2-E17K-NL | 1.9840 | 2.0770 | 1.4120 | 1.8243 | 0.2079 |
| AKT3-NL | 0.4538 | 0.3678 | 0.1546 | 0.3254 | 0.0889 |
| AKT3-E17K-NL | 0.0827 | 0.1434 | 0.4971 | 0.2411 | 0.1292 |

| Capivasertib | Replicate 1 (μM) | Replicate 2 (μM) | Replicate 3 (μM) | Average (μM) | Standard Error |
|---|---|---|---|---|---|
| AKT2-NL | 0.8630 | 1.1680 | 1.6310 | 1.2207 | 0.2233 |
| AKT2-E17K-NL | 1.2020 | 2.6300 | 1.2740 | 1.7020 | 0.4645 |
| AKT3-NL | 0.1709 | 0.3384 | 0.4087 | 0.3060 | 0.0705 |
| AKT3-E17K-NL | 0.5087 | 0.0156 | 0.7948 | 0.4397 | 0.2276 |

| A-443654 | Replicate 1 (μM) | Replicate 2 (μM) | Replicate 3 (μM) | Average (μM) | Standard Error |
|---|---|---|---|---|---|
| AKT2-NL | 0.1291 | 0.5223 | 0.8603 | 0.5039 | 0.2113 |
| AKT2-E17K-NL | 0.7003 | 0.9087 | 0.8643 | 0.8244 | 0.0634 |
| AKT3-NL | 0.1496 | 0.1418 | 0.0241 | 0.1052 | 0.0406 |
| AKT3-E17K-NL | 0.1357 | 0.0175 | 0.0315 | 0.0616 | 0.0373 |

| MK-2206 | Replicate 1 (μM) | Replicate 2 (μM) | Replicate 3 (μM) | Average (μM) | Standard Error |
|---|---|---|---|---|---|
| AKT2-NL | 0.0078 | 0.0014 | 0.0076 | 0.0056 | 0.0021 |
| AKT2-E17K-NL | 0.1427 | 0.1601 | 0.0815 | 0.1281 | 0.0238 |
| AKT3-NL | N/A | N/A | N/A | N/A | N/A |
| AKT3-E17K-NL | N/A | N/A | N/A | N/A | N/A |

| Miransertib | Replicate 1 (μM) | Replicate 2 (μM) | Replicate 3 (μM) | Average (μM) | Standard Error |
|---|---|---|---|---|---|
| AKT2-NL | 0.0032 | 0.2368 | 0.0018 | 0.0806 | 0.0781 |
| AKT2-E17K-NL | 0.0324 | 0.0495 | 0.0073 | 0.0297 | 0.0122 |
| AKT3-NL | N/A | 1.52 | 8.70 | 5.11 | 3.59 |
| AKT3-E17K-NL | N/A | N/A | N/A | N/A | N/A |

| Inhibitor VIII | Replicate 1 (μM) | Replicate 2 (μM) | Replicate 3 (μM) | Average (μM) | Standard Error |
|---|---|---|---|---|---|
| AKT2-NL | 0.4895 | 0.6986 | 0.2136 | 0.4672 | 0.1404 |
| AKT2-E17K-NL | 1.2750 | 0.9939 | 1.1500 | 1.1396 | 0.0813 |
| AKT3-NL | 2.1010 | 2.2910 | 3.0170 | 2.4697 | 0.2791 |
| AKT3-E17K-NL | 5.4410 | N/A | 2.8120 | 4.1265 | 1.3145 |

D

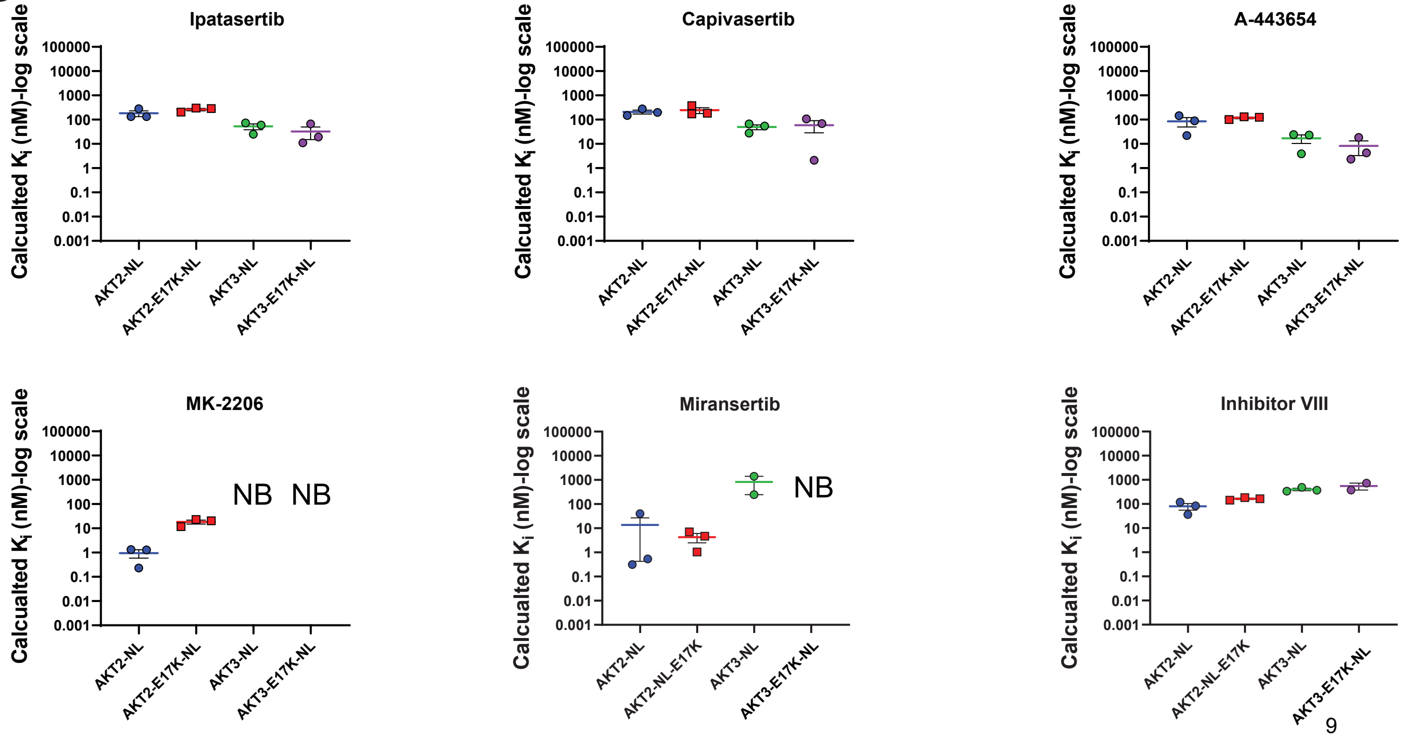

E

| Ipatasertib | Replicate 1 (nM) | Replicate 2 (nM) | Replicate 3 (nM) | Average (nM) | Standard Error |
|---|---|---|---|---|---|
| AKT2-NL | 132.26 | 133.14 | 281.76 | 182.39 | 49.69 |
| AKT2-E17K-NL | 285.13 | 298.49 | 202.92 | 262.18 | 29.88 |
| AKT3-NL | 73.47 | 59.54 | 25.03 | 52.68 | 14.40 |
| AKT3-E17K-NL | 11.13 | 19.31 | 66.94 | 32.46 | 17.40 |

| Capivasertib | Replicate 1 (nM) | Replicate 2 (nM) | Replicate 3 (nM) | Average (nM) | Standard Error |
|---|---|---|---|---|---|
| AKT2-NL | 146.84 | 198.73 | 277.51 | 207.69 | 37.99 |
| AKT2-E17K-NL | 172.74 | 377.97 | 183.09 | 244.60 | 66.75 |
| AKT3-NL | 27.67 | 54.78 | 66.16 | 49.54 | 11.42 |
| AKT3-E17K-NL | 68.50 | 2.10 | 107.03 | 59.21 | 30.64 |

| A-443654 | Replicate 1 (nM) | Replicate 2 (nM) | Replicate 3 (nM) | Average (nM) | Standard Error |
|---|---|---|---|---|---|
| AKT2-NL | 21.97 | 88.87 | 146.38 | 85.74 | 35.95 |
| AKT2-E17K-NL | 100.64 | 130.59 | 124.21 | 118.48 | 9.11 |
| AKT3-NL | 24.22 | 22.96 | 3.90 | 17.03 | 6.57 |
| AKT3-E17K-NL | 18.27 | 2.36 | 4.24 | 8.29 | 5.02 |

| MK-2206 | Replicate 1 (nM) | Replicate 2 (nM) | Replicate 3 (nM) | Average (nM) | Standard Error |
|---|---|---|---|---|---|
| AKT2-NL | 1.32 | 0.23 | 1.28 | 0.94 | 0.36 |
| AKT2-E17K-NL | 20.51 | 23.01 | 11.72 | 18.41 | 3.42 |
| AKT3-NL | N/A | N/A | N/A | N/A | N/A |
| AKT3-E17K-NL | N/A | N/A | N/A | N/A | N/A |

| Miransertib | Replicate 1 (nM) | Replicate 2 (nM) | Replicate 3 (nM) | Average (nM) | Standard Error |
|---|---|---|---|---|---|
| AKT2-NL | 0.54 | 40.29 | 0.31 | 13.71 | 13.29 |
| AKT2-E17K-NL | 4.65 | 7.11 | 1.04 | 4.27 | 1.76 |
| AKT3-NL | N/A | 245.91 | 1409.09 | 827.50 | 581.59 |
| AKT3-E17K-NL | N/A | N/A | N/A | N/A | N/A |

| Inhibitor VIII | Replicate 1 (nM) | Replicate 2 (nM) | Replicate 3 (nM) | Average (nM) | Standard Error |
|---|---|---|---|---|---|
| AKT2-NL | 83.29 | 118.86 | 36.34 | 79.50 | 23.90 |
| AKT2-E17K-NL | 183.23 | 142.84 | 165.27 | 163.78 | 11.69 |
| AKT3-NL | 340.13 | 370.89 | 488.42 | 399.81 | 45.18 |
| AKT3-E17K-NL | 732.67 | N/A | 378.66 | 555.60 | 177.01 |

**Figure S3**- Additional data associated with Figure 3. **(A)** Signal build-up curves and fitted $EC_{50}$ values using the K5 tracer on 293T cells transfected with AKT2-NL and AKT3-NL mutants. **(B)** Replicate binding curves for AKT inhibitors on AKT2-NL and AKT3-NL mutants. **(C)** Fitted apparent $IC_{50}$ values for curves in (B). **(D)** Plotted Calculated $K_i$ values derived from $EC_{50}$ and $IC_{50}$ values in (A and Figure S1A) and (C). **(E)** $K_i$ values plotted in (D). Note that “NB” denotes no binding.

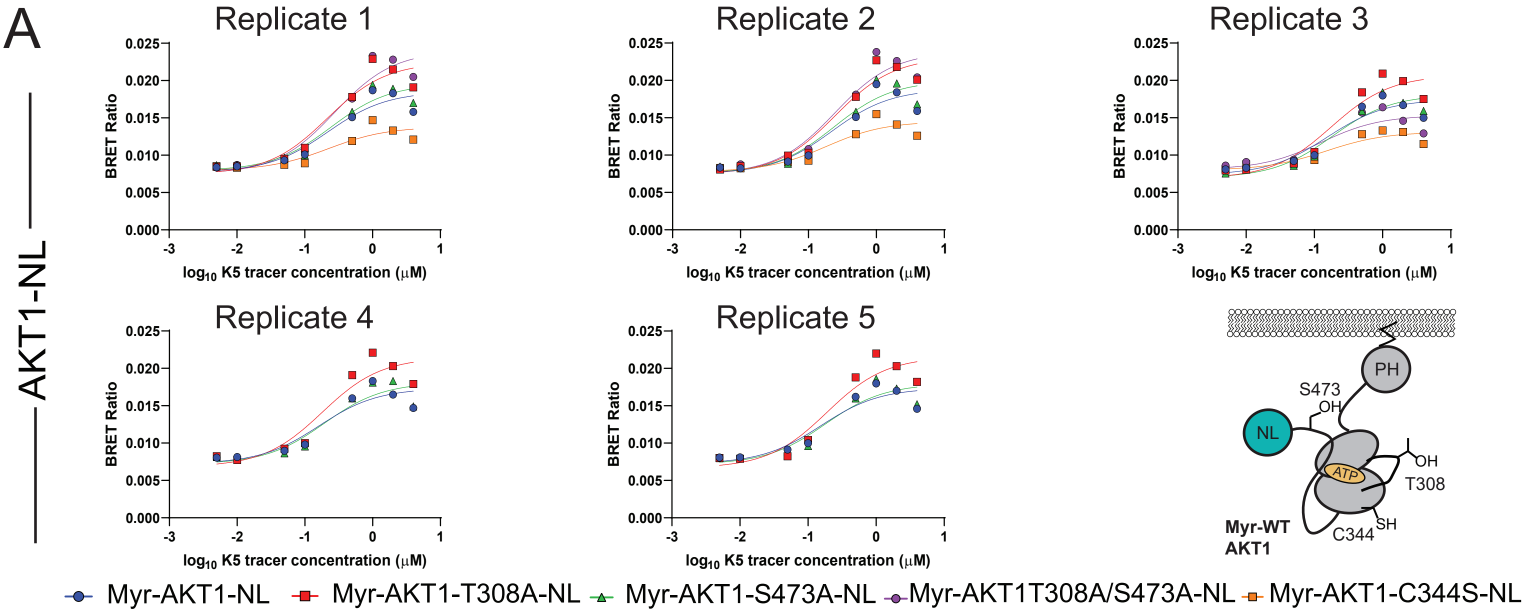


EC$_{50}$ Values

| AKT1 | Replicate 1(μM) | Replicate 2 (μM) | Replicate 3 (μM) | Replicate 4 (μM) | Replicate 5 (μM) | Average (μM) | Standard Error |
|---|---|---|---|---|---|---|---|
| Myr-AKT1-NL | 0.2196 | 0.22 | 0.1546 | 0.1609 | 0.1533 | 0.1817 | 0.0156 |
| Myr-AKT1-T308A-NL | 0.2154 | 0.2469 | 0.1778 | 0.1806 | 0.1889 | 0.2019 | 0.0131 |
| Myr-AKT1-S473A-NL | 0.2433 | 0.2376 | 0.1829 | 0.1942 | 0.1801 | 0.2076 | 0.0136 |
| Myr-AKT1-T308A/S473A-NL | 0.2766 | 0.2442 | 0.1259 | | | 0.2156 | 0.0458 |
| Myr-AKT1-C344S-NL | 0.2224 | 0.1814 | 0.1486 | | | 0.1841 | 0.0213 |

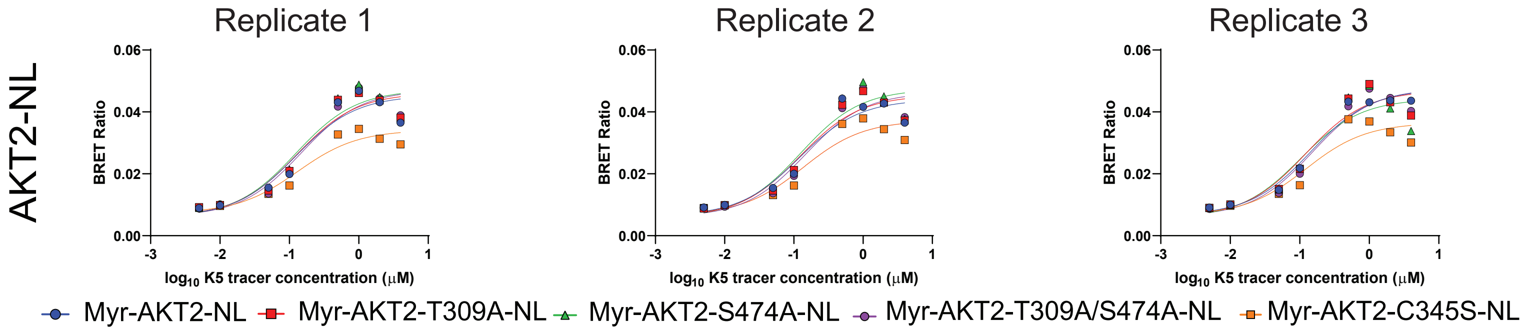


EC$_{50}$ Values

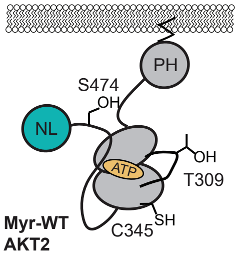


| AKT2 | Replicate 1(μM) | Replicate 2 (μM) | Replicate 3 (μM) | Average (μM) | Standard Error |
|---|---|---|---|---|---|
| Myr-AKT2-NL | 0.1237 | 0.1195 | 0.1420 | 0.1284 | 0.0069 |
| Myr-AKT2-T309A-NL | 0.1269 | 0.1230 | 0.1215 | 0.1238 | 0.0016 |
| Myr-AKT2-S474A-NL | 0.1235 | 0.1247 | 0.1034 | 0.1172 | 0.0069 |
| Myr-AKT2-T309A/S474A-NL | 0.1463 | 0.1458 | 0.1502 | 0.1474 | 0.0014 |
| Myr-AKT2-C345S-NL | 0.1304 | 0.1360 | 0.1219 | 0.1294 | 0.0041 |

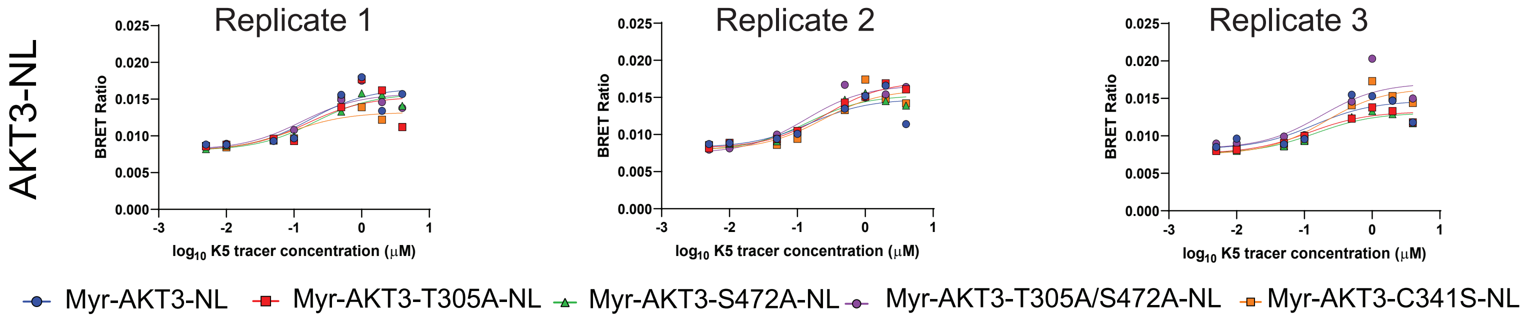


EC$_{50}$ Values

| AKT3 | Replicate 1(μM) | Replicate 2 (μM) | Replicate 3 (μM) | Average (μM) | Standard Error |
|---|---|---|---|---|---|
| Myr-AKT3-NL | 0.1766 | 0.1534 | 0.14 | 0.1567 | 0.0107 |
| Myr-AKT3-T305A-NL | 0.1529 | 0.261 | 0.116 | 0.1766 | 0.0435 |
| Myr-AKT3-S472A-NL | 0.2026 | 0.1392 | 0.147 | 0.1629 | 0.0200 |
| Myr-AKT3-T305A/S472A-NL | 0.1293 | 0.1332 | 0.1752 | 0.1459 | 0.0147 |
| Myr-AKT3-C341S-NL | 0.0873 | 0.2345 | 0.204 | 0.1753 | 0.0449 |

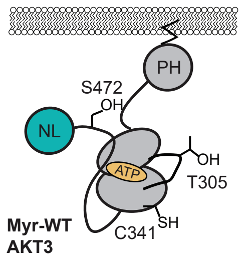

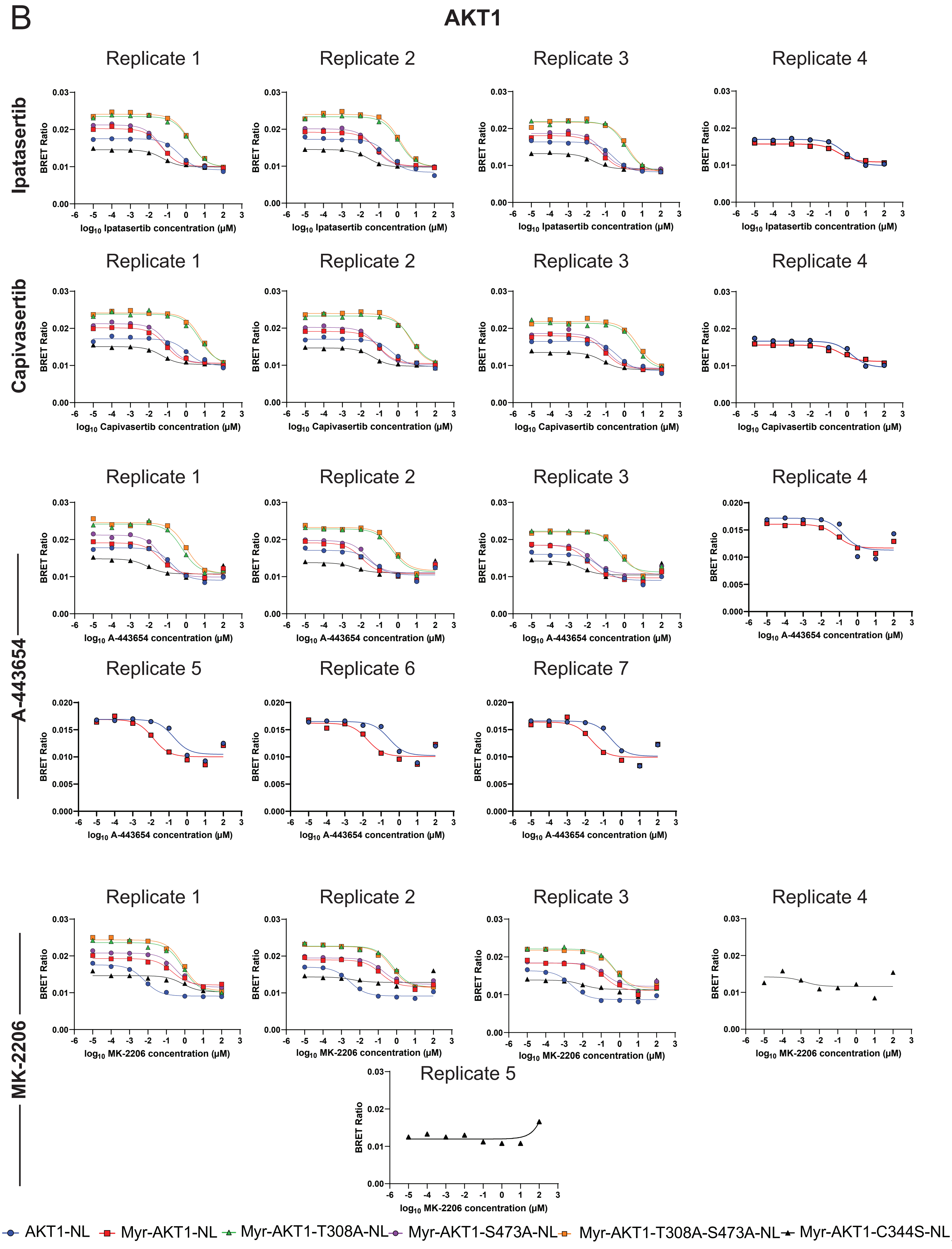
B
AKT1
Ipatasertib
Capivasertib
A-443654
MK-2206
Replicate 1
Replicate 2
Replicate 3
Replicate 4
Replicate 5
Replicate 6
Replicate 7
BRET Ratio
log10 Ipatasertib concentration (µM)
log10 Capivasertib concentration (µM)
log10 A-443654 concentration (µM)
log10 MK-2206 concentration (µM)
AKT1-NL
Myr-AKT1-NL
Myr-AKT1-T308A-NL
Myr-AKT1-S473A-NL
Myr-AKT1-T308A-S473A-NL
Myr-AKT1-C344S-NL

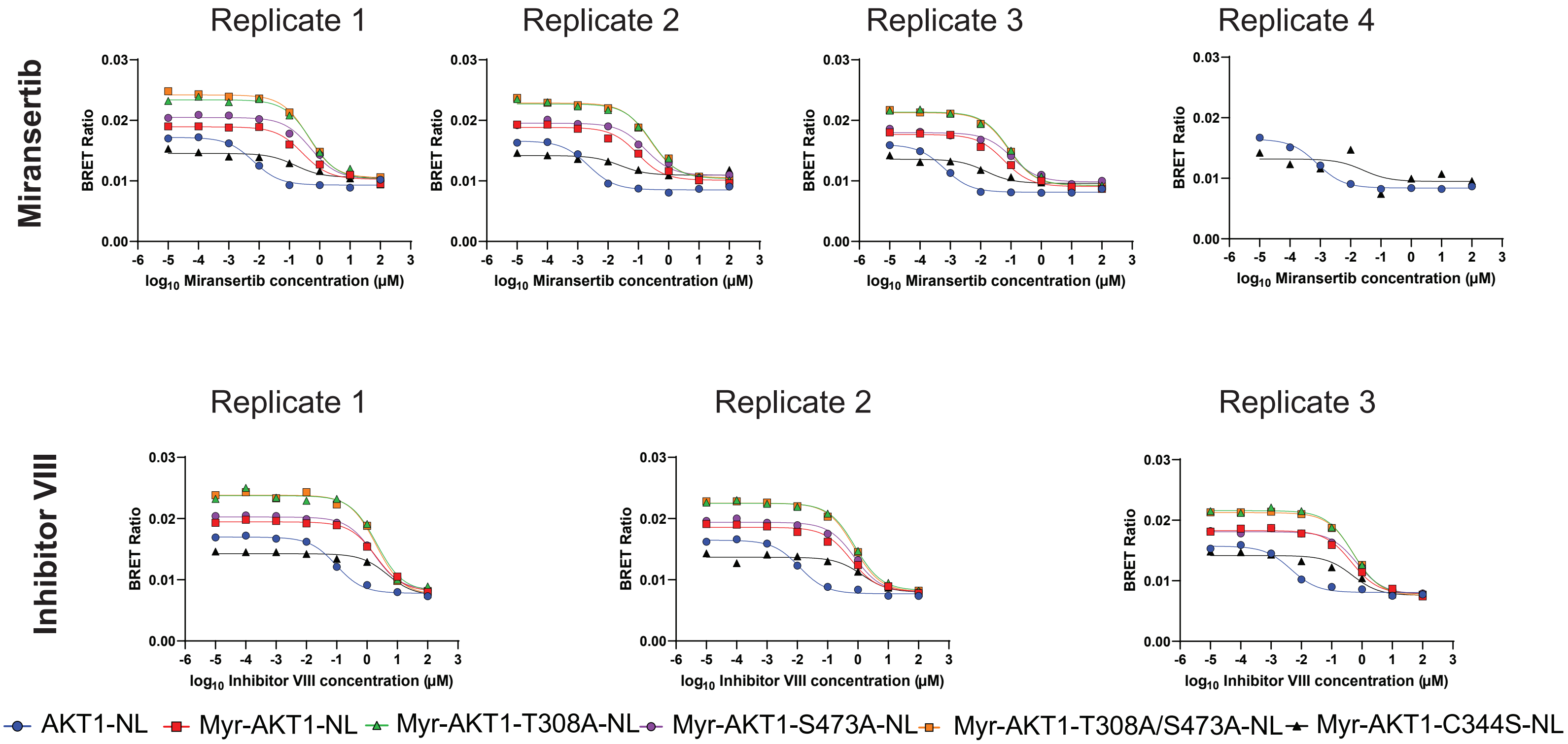


# C

#### Apparent $IC_{50}$ Values

| Ipatasertib | Replicate 1 (µM) | Replicate 2 (µM) | Replicate 3 (µM) | Replicate 4 (µM) | Average (µM) | Standard Error |
|---|---|---|---|---|---|---|
| AKT1-NL | 0.4446 | 0.3463 | 0.2727 | 0.7315 | 0.4488 | 0.1006 |
| Myr-AKT1-NL | 0.0502 | 0.0744 | 0.0641 | 0.3554 | 0.1360 | 0.0733 |
| Myr-AKT1-T308A-NL | 1.5760 | 1.2390 | 1.2290 | | 1.3480 | 0.1140 |
| Myr-AKT1-S473A-NL | 0.0454 | 0.0581 | 0.0792 | | 0.0609 | 0.0099 |
| Myr-AKT1-T308A-S473A-NL | 1.5320 | 1.3750 | 1.5860 | | 1.4977 | 0.0633 |
| Myr-AKT1-C344S-NL | 0.0555 | 0.0264 | 0.0247 | | 0.0356 | 0.0100 |

| Capivasertib | Replicate 1 (µM) | Replicate 2 (µM) | Replicate 3 (µM) | Replicate 4 (µM) | Average (µM) | Standard Error |
|---|---|---|---|---|---|---|
| AKT1-NL | 1.1590 | 0.5347 | 0.4594 | 1.6560 | 0.9523 | 0.2821 |
| Myr-AKT1-NL | 0.0808 | 0.0979 | 0.0919 | 0.4651 | 0.1839 | 0.0938 |
| Myr-AKT1-T308A-NL | 5.2900 | 5.4340 | 4.1430 | | 4.9557 | 0.4085 |
| Myr-AKT1-S473A-NL | 0.1016 | 0.0792 | 0.1129 | | 0.0979 | 0.0099 |
| Myr-AKT1-T308A-S473A-NL | 6.5200 | 4.4800 | 5.1900 | | 5.3967 | 0.5979 |
| Myr-AKT1-C344S-NL | 0.0488 | 0.0448 | 0.0692 | | 0.0543 | 0.0076 |

| A-443654 | Replicate 1 (µM) | Replicate 2 (µM) | Replicate 3 (µM) | Replicate 4 (µM) | Replicate 5 (µM) | Replicate 6 (µM) | Replicate 7 (µM) | Average (µM) | Standard Error |
|---|---|---|---|---|---|---|---|---|---|
| AKT1-NL | 0.1704 | 0.0405 | 0.0538 | 0.1650 | 0.1758 | 0.2730 | 0.2575 | 0.1623 | 0.0339 |
| Myr-AKT1-NL | 0.0403 | 0.0112 | 0.0103 | 0.0740 | 0.0133 | 0.0178 | 0.0206 | 0.0268 | 0.0088 |
| Myr-AKT1-T308A-NL | 0.6119 | 0.4913 | 0.4175 | | | | | 0.5069 | 0.0567 |
| Myr-AKT1-S473A-NL | 0.0486 | 0.0218 | 0.0186 | | | | | 0.0297 | 0.0095 |
| Myr-AKT1-T308A-S473A-NL | 1.0160 | 0.5635 | 0.6601 | | | | | 0.7465 | 0.1376 |
| Myr-AKT1-C344S-NL | 0.0082 | 0.0046 | 0.0046 | | | | | 0.0058 | 0.0012 |

| MK-2206 | Replicate 1 (µM) | Replicate 2 (µM) | Replicate 3 (µM) | Replicate 4 (µM) | Replicate 5 (µM) | Average (µM) | Standard Error |
|---|---|---|---|---|---|---|---|
| AKT1-NL | 0.0065 | 0.0028 | 0.0025 | | | 0.0039 | 0.0013 |
| Myr-AKT1-NL | 0.2072 | 0.1358 | 0.1484 | | | 0.1638 | 0.0220 |
| Myr-AKT1-T308A-NL | 0.7748 | 0.6234 | 0.5784 | | | 0.6589 | 0.0594 |
| Myr-AKT1-S473A-NL | 0.3440 | 0.1792 | 0.1719 | | | 0.2317 | 0.0562 |
| Myr-AKT1-T308A-S473A-NL | 0.8123 | 0.5354 | 0.5055 | | | 0.6177 | 0.0977 |
| Myr-AKT1-C344S-NL | 0.7654 | 0.0040 | 0.0097 | 0.0015 | 0.0555 | 0.1672 | 0.1499 |

| Miransertib | Replicate 1 (µM) | Replicate 2 (µM) | Replicate 3 (µM) | Replicate 4 (µM) | Average (µM) | Standard Error |
|---|---|---|---|---|---|---|
| AKT1-NL | 0.0065 | 0.0022 | 0.0006 | 0.0008 | 0.0025 | 0.0014 |
| Myr-AKT1-NL | 0.2706 | 0.0953 | 0.0591 | | 0.1417 | 0.0653 |
| Myr-AKT1-T308A-NL | 0.4973 | 0.2705 | 0.0855 | | 0.2844 | 0.1191 |
| Myr-AKT1-S473A-NL | 0.4985 | 0.1694 | 0.1069 | | 0.2583 | 0.1215 |
| Myr-AKT1-T308A-S473A-NL | 0.4170 | 0.2454 | 0.0827 | | 0.2484 | 0.0965 |
| Myr-AKT1-C344S-NL | 0.1571 | 0.0248 | 0.0149 | 0.0261 | 0.0557 | 0.0339 |

| Inhibitor VIII | Replicate 1 (µM) | Replicate 2 (µM) | Replicate 3 (µM) | Average (µM) | Standard Error |
|---|---|---|---|---|---|
| AKT1-NL | 0.0962 | 0.0117 | 0.0045 | 0.0375 | 0.0294 |
| Myr-AKT1-NL | 2.0050 | 0.5831 | 0.4534 | 1.0138 | 0.4970 |
| Myr-AKT1-T308A-NL | 2.0810 | 0.8179 | 0.4901 | 1.1297 | 0.4850 |
| Myr-AKT1-S473A-NL | 1.6180 | 0.7689 | 0.6730 | 1.0200 | 0.3003 |
| Myr-AKT1-T308A-S473A-NL | 1.9970 | 0.7452 | 0.5350 | 1.0924 | 0.4564 |
| Myr-AKT1-C344S-NL | 4.7720 | 1.3600 | 0.4882 | 2.2067 | 1.3071 |

D

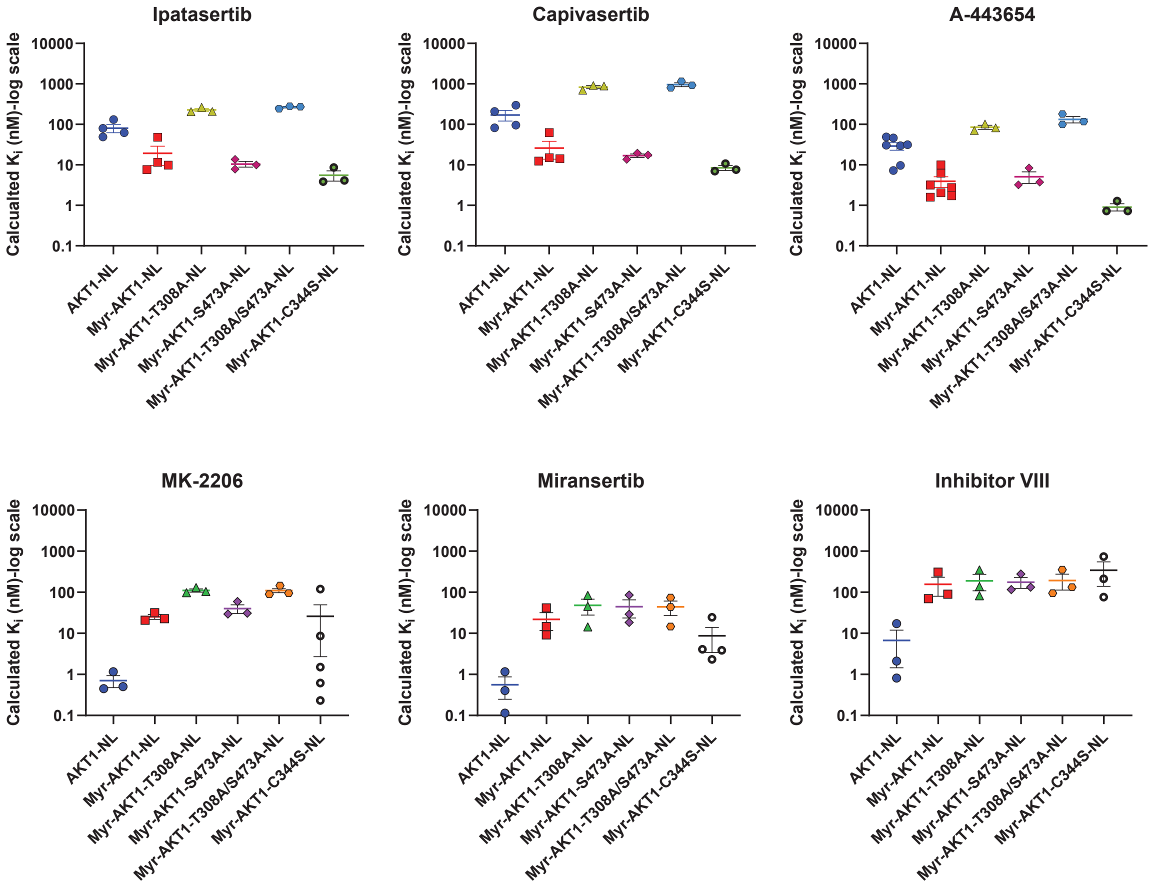


E

#### Calculated $K_i$ Values

| Ipatasertib | Replicate 1 (nM) | Replicate 2 (nM) | Replicate 3 (nM) | Replicate 4 (nM) | Average (nM) | Standard Error |
|---|---|---|---|---|---|---|
| AKT1-NL | 79.71 | 62.09 | 48.89 | 131.15 | 80.46 | 18.04 |
| Myr-AKT1-NL | 7.713 | 11.44 | 9.86 | 47.86 | 19.22 | 9.58 |
| Myr-AKT1-T308A-NL | 264.76 | 208.15 | 206.47 | | 226.46 | 19.16 |
| Myr-AKT1-S473A-NL | 7.81 | 10.00 | 13.62 | | 10.48 | 1.69 |
| Myr-AKT1-T308A-S473A-NL | 271.68 | 243.84 | 281.26 | | 265.59 | 11.22 |
| Myr-AKT1-C344S-NL | 8.64 | 4.10 | 3.84 | | 5.53 | 1.56 |

| Capivasertib | Replicate 1 (nM) | Replicate 2 (nM) | Replicate 3 (nM) | Replicate 4 (nM) | Average (nM) | Standard Error |
|---|---|---|---|---|---|---|
| AKT1-NL | 207.80 | 95.87 | 82.37 | 296.91 | 170.74 | 50.59 |
| Myr-AKT1-NL | 12.42 | 15.05 | 14.13 | 62.63 | 26.06 | 12.20 |
| Myr-AKT1-T308A-NL | 888.71 | 912.90 | 696.02 | | 832.54 | 68.62 |
| Myr-AKT1-S473A-NL | 17.47 | 13.62 | 19.41 | | 16.83 | 1.70 |
| Myr-AKT1-T308A-S473A-NL | 1156.25 | 794.48 | 920.39 | | 957.04 | 106.03 |
| Myr-AKT1-C344S-NL | 7.59 | 6.96 | 10.76 | | 8.44 | 1.18 |

| A-443654 | Replicate 1 (nM) | Replicate 2 (nM) | Replicate 3 (nM) | Replicate 4 (nM) | Replicate 5 (nM) | Replicate 6 (nM) | Replicate 7 (nM) | Average (nM) | Standard Error |
|---|---|---|---|---|---|---|---|---|---|
| AKT1-NL | 30.55 | 7.26 | 9.65 | 29.58 | 31.52 | 48.95 | 46.17 | 29.10 | 6.07 |
| Myr-AKT1-NL | 6.19 | 1.72 | 1.59 | 9.96 | 2.05 | 2.74 | 3.17 | 3.92 | 1.17 |
| Myr-AKT1-T308A-NL | 102.80 | 82.54 | 70.14 | | | | | 85.16 | 9.52 |
| Myr-AKT1-S473A-NL | 8.35 | 3.75 | 3.19 | | | | | 5.10 | 1.63 |
| Myr-AKT1-T308A-S473A-NL | 180.18 | 99.93 | 117.06 | | | | | 132.39 | 24.40 |
| Myr-AKT1-C344S-NL | 1.28 | 0.72 | 0.72 | | | | | 0.91 | 0.19 |

| MK-2206 | Replicate 1 (nM) | Replicate 2 (nM) | Replicate 3 (nM) | Replicate 4 (nM) | Replicate 5 (nM) | Average (nM) | Standard Error |
|---|---|---|---|---|---|---|---|
| AKT1-NL | 1.16 | 0.51 | 0.45 | | | 0.70 | 0.23 |
| Myr-AKT1-NL | 31.86 | 20.88 | 22.82 | | | 25.18 | 3.38 |
| Myr-AKT1-T308A-NL | 130.16 | 104.73 | 97.17 | | | 110.69 | 9.98 |
| Myr-AKT1-S473A-NL | 59.14 | 30.81 | 29.55 | | | 39.84 | 9.66 |
| Myr-AKT1-T308A-S473A-NL | 144.05 | 94.95 | 89.64 | | | 109.55 | 17.32 |
| Myr-AKT1-C344S-NL | 119.02 | 0.61 | 1.51 | 0.23 | 8.63 | 26.00 | 23.31 |

| Miransertib | Replicate 1 (nM) | Replicate 2 (nM) | Replicate 3 (nM) | Replicate 4 (nM) | Average (nM) | Standard Error |
|---|---|---|---|---|---|---|
| AKT1-NL | 1.17 | 0.40 | 0.11 | 0.14 | 0.56 | 0.31 |
| Myr-AKT1-NL | 41.60 | 14.65 | 9.09 | | 21.78 | 10.04 |
| Myr-AKT1-T308A-NL | 83.55 | 45.44 | 14.37 | | 47.79 | 20.00 |
| Myr-AKT1-S473A-NL | 85.70 | 29.12 | 18.38 | | 44.40 | 20.88 |
| Myr-AKT1-T308A-S473A-NL | 73.95 | 43.52 | 14.67 | | 44.05 | 17.11 |
| Myr-AKT1-C344S-NL | 24.43 | 3.85 | 2.32 | 4.06 | 8.66 | 5.27 |

| Inhibitor VIII | Replicate 1 (nM) | Replicate 2 (nM) | Replicate 3 (nM) | Average (nM) | Standard Error |
|---|---|---|---|---|---|
| AKT1-NL | 17.24 | 2.10 | 0.81 | 6.72 | 5.28 |
| Myr-AKT1-NL | 308.26 | 89.65 | 69.71 | 155.87 | 76.41 |
| Myr-AKT1-T308A-NL | 349.60 | 137.41 | 82.34 | 189.78 | 81.48 |
| Myr-AKT1-S473A-NL | 278.17 | 132.19 | 115.71 | 175.36 | 51.63 |
| Myr-AKT1-T308A-S473A-NL | 354.14 | 132.15 | 94.88 | 193.72 | 80.93 |
| Myr-AKT1-C344S-NL | 742.05 | 211.48 | 75.92 | 343.15 | 203.25 |

F

AKT2

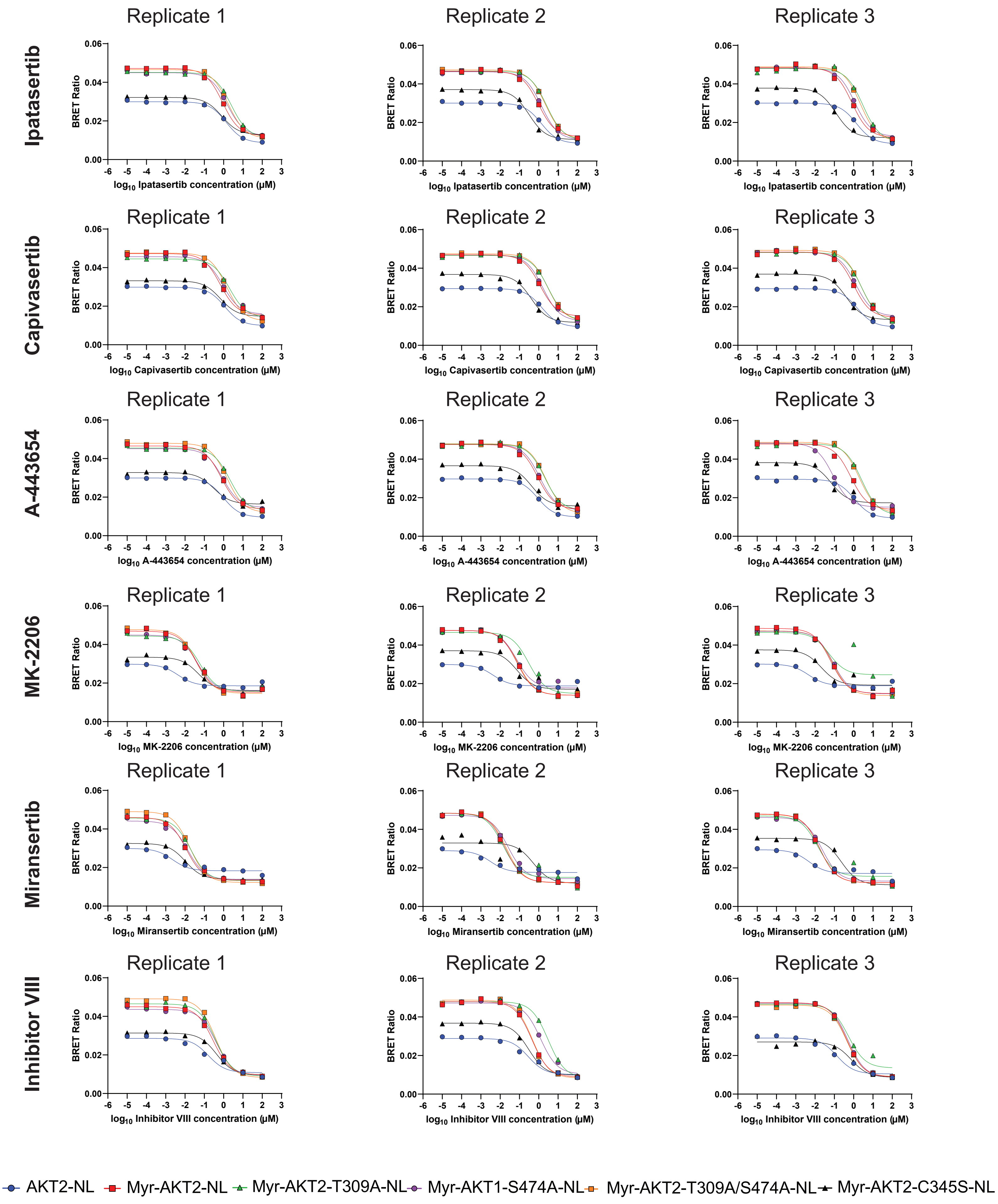
Ipatasertib
Capivasertib
A-443654
MK-2206
Miransertib
Inhibitor VIII
Replicate 1
Replicate 2
Replicate 3
BRET Ratio
0.06
0.04
0.02
0.00
-6 -5 -4 -3 -2 -1 0 1 2 3
log10 Ipatasertib concentration (μM)
log10 Capivasertib concentration (μM)
log10 A-443654 concentration (μM)
log10 MK-2206 concentration (μM)
log10 Miransertib concentration (μM)
log10 Inhibitor VIII concentration (μM)
AKT2-NL
Myr-AKT2-NL
Myr-AKT2-T309A-NL
Myr-AKT1-S474A-NL
Myr-AKT2-T309A/S474A-NL
Myr-AKT2-C345S-NL

### G Apparent $IC_{50}$ Values

| Ipatasertib | Replicate 1 (µM) | Replicate 2 (µM) | Replicate 3 (µM) | Average (µM) | Standard Error |
|---|---|---|---|---|---|
| AKT2-NL | 1.3790 | 1.3440 | 1.4020 | 1.3750 | 0.0169 |
| Myr-AKT2-NL | 0.8675 | 0.9573 | 0.8078 | 0.8775 | 0.0434 |
| Myr-AKT2-T309A-NL | 2.4750 | 2.5710 | 2.9760 | 2.6740 | 0.1535 |
| Myr-AKT2-S474A-NL | 1.5160 | 1.3790 | 1.1420 | 1.3457 | 0.1092 |
| Myr-AKT2-T309A-S474A-NL | 1.7490 | 2.2480 | 2.2410 | 2.0793 | 0.1652 |
| Myr-AKT2-C345S-NL | 0.8083 | 0.3157 | 0.1203 | 0.4148 | 0.2047 |

| Capivasertib | Replicate 1 (µM) | Replicate 2 (µM) | Replicate 3 (µM) | Average (µM) | Standard Error |
|---|---|---|---|---|---|
| AKT2-NL | 1.1280 | 1.4710 | 1.4990 | 1.3660 | 0.1193 |
| Myr-AKT2-NL | 0.6666 | 1.1720 | 0.9790 | 0.9392 | 0.1472 |
| Myr-AKT2-T309A-NL | 1.9980 | 3.0850 | 2.5290 | 2.5373 | 0.3138 |
| Myr-AKT2-S474A-NL | 0.9233 | 1.6560 | 1.2920 | 1.2904 | 0.2115 |
| Myr-AKT2-T309A-S474A-NL | 1.4550 | 2.9200 | 2.3510 | 2.2420 | 0.4264 |
| Myr-AKT2-C345S-NL | 0.6808 | 0.4054 | 0.4076 | 0.4979 | 0.0914 |

| A-443654 | Replicate 1 (µM) | Replicate 2 (µM) | Replicate 3 (µM) | Average (µM) | Standard Error |
|---|---|---|---|---|---|
| AKT2-NL | 0.9754 | 0.8890 | 1.0780 | 0.9808 | 0.0546 |
| Myr-AKT2-NL | 0.7544 | 0.8746 | 0.7114 | 0.7801 | 0.0488 |
| Myr-AKT2-T309A-NL | 2.0840 | 2.0060 | 2.5390 | 2.2097 | 0.1662 |
| Myr-AKT2-S474A-NL | 0.8956 | 1.1160 | 0.0840 | 0.6985 | 0.3138 |
| Myr-AKT2-T309A-S474A-NL | 1.4820 | 2.2940 | 1.9520 | 1.9093 | 0.2354 |
| Myr-AKT2-C345S-NL | 0.3088 | 0.4220 | 0.0680 | 0.2663 | 0.1044 |

| MK-2206 | Replicate 1 (µM) | Replicate 2 (µM) | Replicate 3 (µM) | Average (µM) | Standard Error |
|---|---|---|---|---|---|
| AKT2-NL | 0.0044 | 0.0044 | 0.0046 | 0.0044 | 0.0001 |
| Myr-AKT2-NL | 0.0377 | 0.0759 | 0.0644 | 0.0593 | 0.0113 |
| Myr-AKT2-T309A-NL | 0.0587 | 0.3011 | 0.0524 | 0.1374 | 0.0819 |
| Myr-AKT2-S474A-NL | 0.0440 | 0.0705 | 0.0856 | 0.0667 | 0.0122 |
| Myr-AKT2-T309A-S474A-NL | 0.0396 | 0.0818 | 0.0830 | 0.0681 | 0.0143 |
| Myr-AKT2-C345S-NL | 0.0501 | 0.0734 | 0.0213 | 0.0483 | 0.0151 |

| Miransertib | Replicate 1 (µM) | Replicate 2 (µM) | Replicate 3 (µM) | Average (µM) | Standard Error |
|---|---|---|---|---|---|
| AKT2-NL | 0.0022 | 0.0031 | 0.0048 | 0.0034 | 0.0007 |
| Myr-AKT2-NL | 0.0113 | 0.0183 | 0.0174 | 0.0157 | 0.0022 |
| Myr-AKT2-T309A-NL | 0.0196 | 0.0139 | 0.0136 | 0.0157 | 0.0019 |
| Myr-AKT2-S474A-NL | 0.0141 | 0.0257 | 0.0231 | 0.0210 | 0.0035 |
| Myr-AKT2-T309A-S474A-NL | 0.0172 | 0.0209 | 0.0173 | 0.0185 | 0.0012 |
| Myr-AKT2-C345S-NL | 0.0105 | 0.5261 | 0.2255 | 0.2540 | 0.1495 |

| Inhibitor VIII | Replicate 1 (µM) | Replicate 2 (µM) | Replicate 3 (µM) | Average (µM) | Standard Error |
|---|---|---|---|---|---|
| AKT2-NL | 0.1617 | 0.2822 | 0.1373 | 0.1937 | 0.0448 |
| Myr-AKT2-NL | 0.2909 | 0.4360 | 0.4450 | 0.3906 | 0.0499 |
| Myr-AKT2-T309A-NL | 0.3852 | 3.0730 | 0.5332 | 1.3305 | 0.8723 |
| Myr-AKT2-S474A-NL | 0.4180 | 1.2280 | 0.5092 | 0.7184 | 0.2562 |
| Myr-AKT2-T309A-S474A-NL | 0.3821 | 0.4589 | 0.4408 | 0.4273 | 0.0232 |
| Myr-AKT2-C345S-NL | 0.4216 | 0.2956 | 0.7540 | 0.4904 | 0.1367 |

### I Calculated $K_i$ Values

| Ipatasertib | Replicate 1 (nM) | Replicate 2 (nM) | Replicate 3 (nM) | Average (nM) | Standard Error |
|---|---|---|---|---|---|
| AKT2-NL | 234.6333 | 228.6782 | 238.5467 | 233.9527 | 2.8691 |
| Myr-AKT2-NL | 98.7123 | 108.9306 | 91.9191 | 99.8540 | 4.9439 |
| Myr-AKT2-T309A-NL | 272.6508 | 283.2264 | 327.8420 | 294.5731 | 16.9123 |
| Myr-AKT2-S474A-NL | 159.0362 | 144.6642 | 119.8016 | 141.1673 | 11.4602 |
| Myr-AKT2-T309A-S474A-NL | 224.7284 | 288.8448 | 287.9454 | 267.1729 | 21.2238 |
| Myr-AKT2-C345S-NL | 92.6314 | 36.1793 | 13.7864 | 47.5324 | 23.4578 |

| Capivasertib | Replicate 1 (nM) | Replicate 2 (nM) | Replicate 3 (nM) | Average (nM) | Standard Error |
|---|---|---|---|---|---|
| AKT2-NL | 191.9263 | 250.2869 | 255.0510 | 232.4214 | 20.2942 |
| Myr-AKT2-NL | 75.8520 | 133.3612 | 111.3999 | 106.8710 | 16.7552 |
| Myr-AKT2-T309A-NL | 220.1036 | 339.8496 | 278.5996 | 279.5176 | 34.5708 |
| Myr-AKT2-S474A-NL | 96.8589 | 173.7229 | 135.5374 | 135.3731 | 22.1889 |
| Myr-AKT2-T309A-S474A-NL | 186.9525 | 375.1898 | 302.0792 | 288.0738 | 54.7888 |
| Myr-AKT2-C345S-NL | 78.0198 | 46.4589 | 46.7111 | 57.0633 | 10.4785 |

| A-443654 | Replicate 1 (nM) | Replicate 2 (nM) | Replicate 3 (nM) | Average (nM) | Standard Error |
|---|---|---|---|---|---|
| AKT2-NL | 165.9618 | 151.2611 | 183.4189 | 166.8806 | 9.2945 |
| Myr-AKT2-NL | 85.8428 | 99.5202 | 80.9498 | 88.7709 | 5.5572 |
| Myr-AKT2-T309A-NL | 229.5775 | 220.9849 | 279.7012 | 243.4212 | 18.3088 |
| Myr-AKT2-S474A-NL | 93.9530 | 117.0741 | 8.8162 | 73.2811 | 32.9162 |
| Myr-AKT2-T309A-S474A-NL | 190.4217 | 294.7553 | 250.8118 | 245.3296 | 30.2430 |
| Myr-AKT2-C345S-NL | 35.3886 | 48.3613 | 7.7951 | 30.5150 | 11.9613 |

| MK-2206 | Replicate 1 (nM) | Replicate 2 (nM) | Replicate 3 (nM) | Average (nM) | Standard Error |
|---|---|---|---|---|---|
| AKT2-NL | 0.7408 | 0.7408 | 0.7888 | 0.7568 | 0.0160 |
| Myr-AKT2-NL | 4.2933 | 8.6378 | 7.3258 | 6.7523 | 1.2865 |
| Myr-AKT2-T309A-NL | 6.4654 | 33.1698 | 5.7769 | 15.1374 | 9.0184 |
| Myr-AKT2-S474A-NL | 4.6106 | 7.3969 | 8.9778 | 6.9951 | 1.2766 |
| Myr-AKT2-T309A-S474A-NL | 5.0895 | 10.5053 | 10.6595 | 8.7514 | 1.8315 |
| Myr-AKT2-C345S-NL | 5.7369 | 8.4162 | 2.4375 | 5.5302 | 1.7290 |

| Miransertib | Replicate 1 (nM) | Replicate 2 (nM) | Replicate 3 (nM) | Average (nM) | Standard Error |
|---|---|---|---|---|---|
| AKT2-NL | 0.3827 | 0.5358 | 0.8182 | 0.5789 | 0.1276 |
| Myr-AKT2-NL | 1.2892 | 2.0801 | 1.9799 | 1.7831 | 0.2486 |
| Myr-AKT2-T309A-NL | 2.1581 | 1.5357 | 1.4993 | 1.7310 | 0.2138 |
| Myr-AKT2-S474A-NL | 1.4760 | 2.6992 | 2.4223 | 2.1992 | 0.3703 |
| Myr-AKT2-T309A-S474A-NL | 2.2139 | 2.6880 | 2.2216 | 2.3745 | 0.1568 |
| Myr-AKT2-C345S-NL | 1.2067 | 60.2912 | 25.8424 | 29.1134 | 17.1344 |

| Inhibitor VIII | Replicate 1 (nM) | Replicate 2 (nM) | Replicate 3 (nM) | Average (nM) | Standard Error |
|---|---|---|---|---|---|
| AKT2-NL | 27.5128 | 48.0156 | 23.3612 | 32.9632 | 7.6210 |
| Myr-AKT2-NL | 33.1013 | 49.6122 | 50.6363 | 44.4499 | 5.6820 |
| Myr-AKT2-T309A-NL | 42.4344 | 338.5277 | 58.7384 | 146.5668 | 96.0958 |
| Myr-AKT2-S474A-NL | 43.8503 | 128.8235 | 53.4177 | 75.3638 | 26.8721 |
| Myr-AKT2-T309A-S474A-NL | 49.0959 | 58.9639 | 56.6382 | 54.8994 | 2.9784 |
| Myr-AKT2-C345S-NL | 48.3155 | 33.8758 | 86.4086 | 56.2000 | 15.6689 |

H

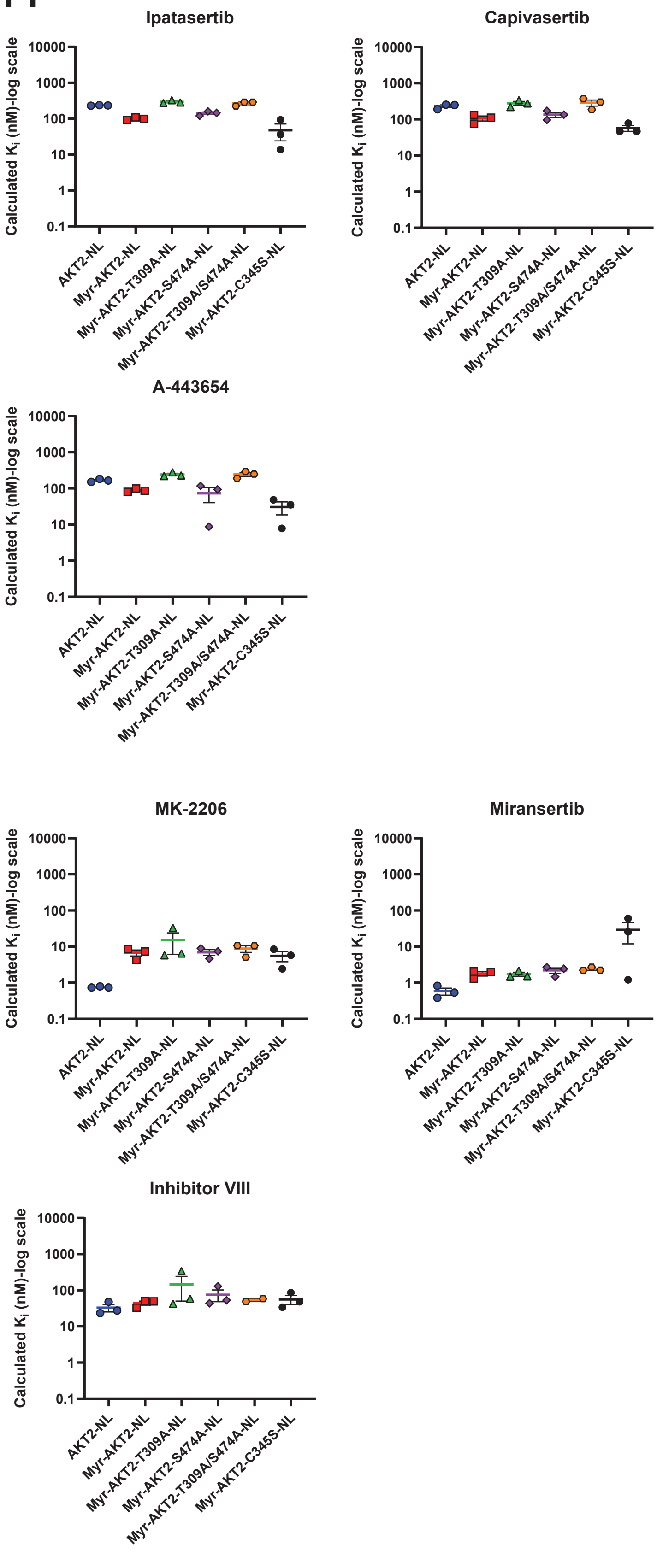

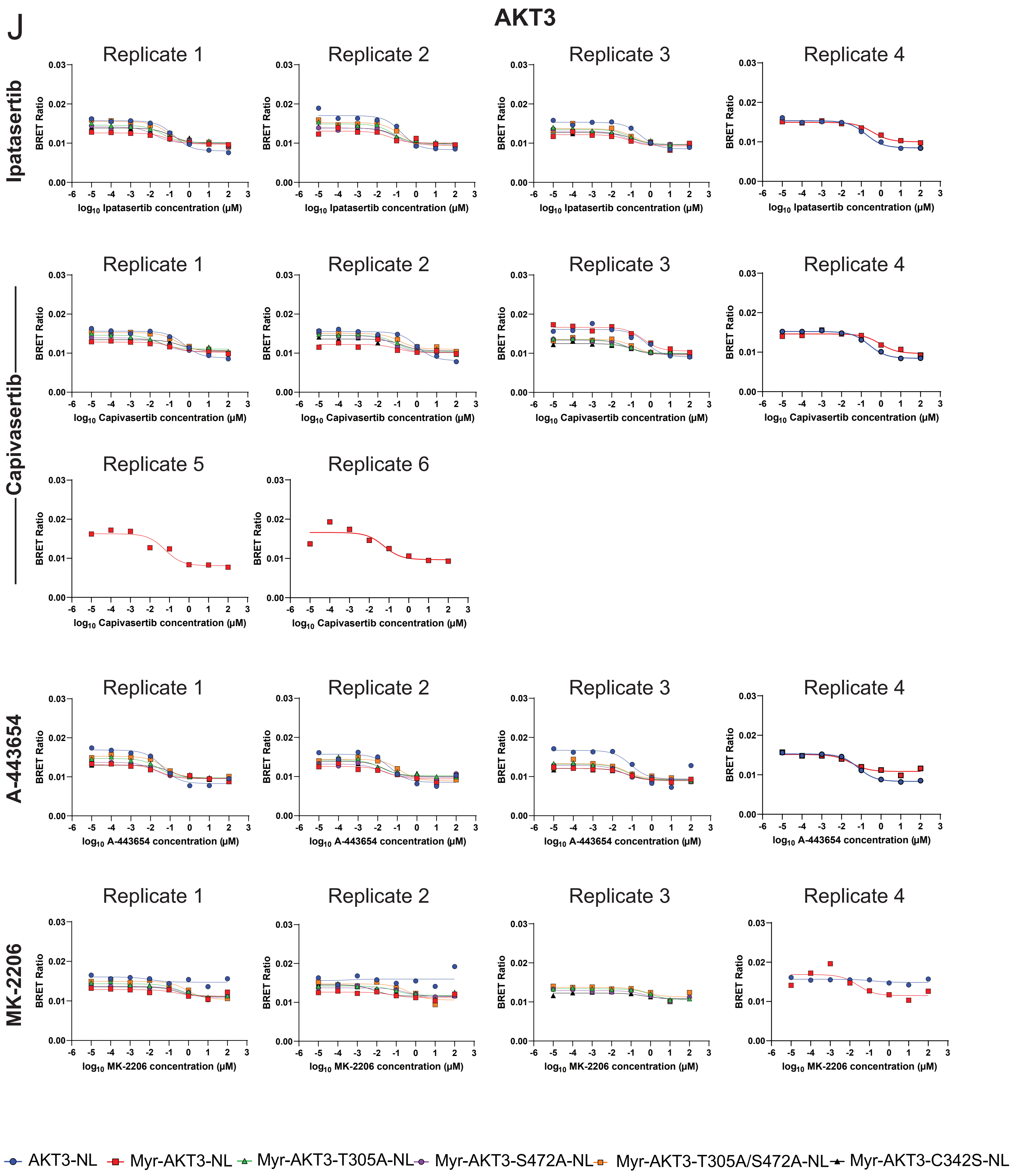
J
AKT3
Ipatasertib
Capivasertib
A-443654
MK-2206
Replicate 1
Replicate 2
Replicate 3
Replicate 4
Replicate 5
Replicate 6
BRET Ratio
log10 Ipatasertib concentration (µM)
log10 Capivasertib concentration (µM)
log10 A-443654 concentration (µM)
log10 MK-2206 concentration (µM)
AKT3-NL
Myr-AKT3-NL
Myr-AKT3-T305A-NL
Myr-AKT3-S472A-NL
Myr-AKT3-T305A/S472A-NL
Myr-AKT3-C342S-NL

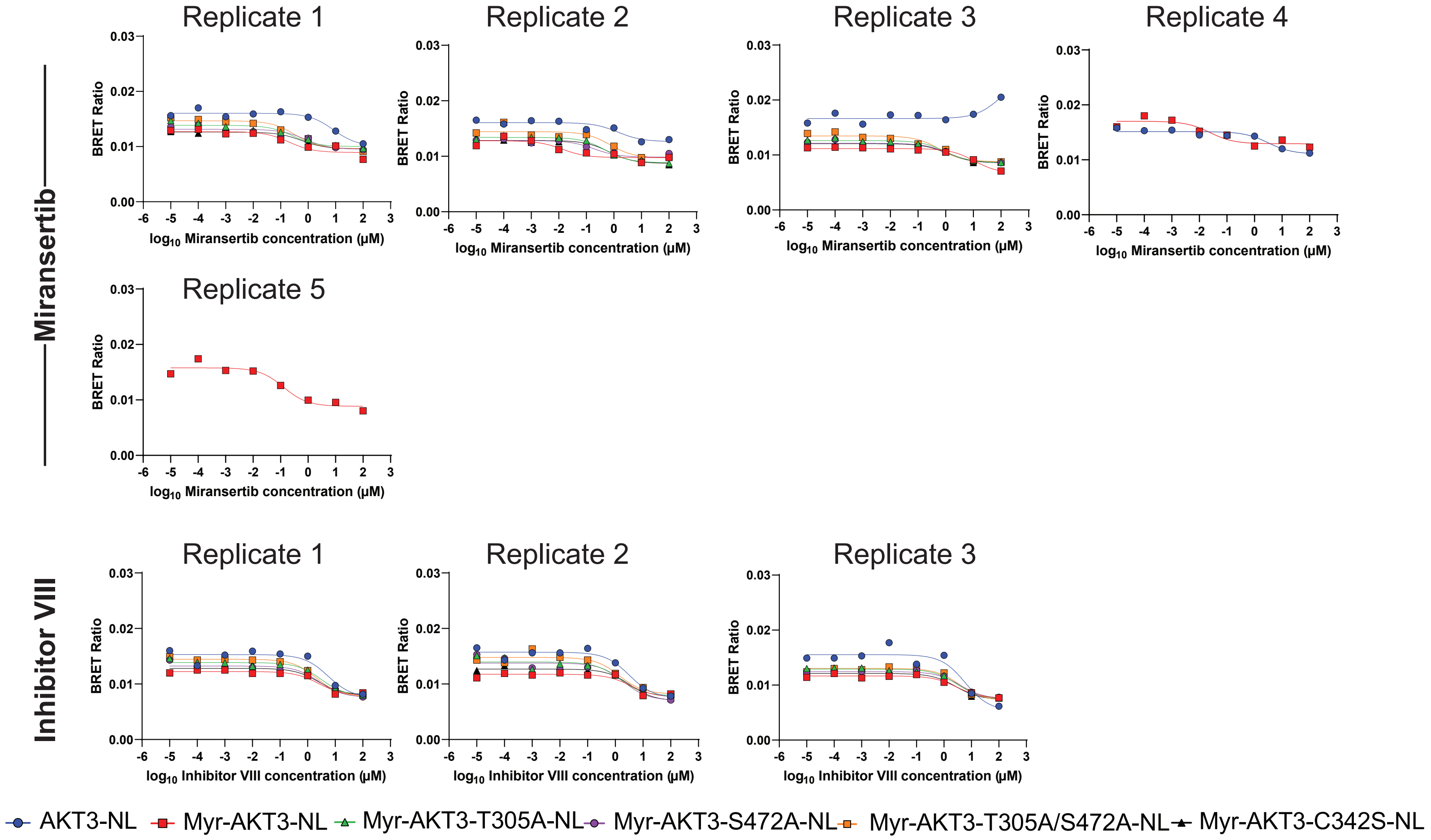


K

#### Apparent $IC_{50}$ Values

| Ipatasertib | Replicate 1 (µM) | Replicate 2 (µM) | Replicate 3 (µM) | Replicate 4 (µM) | Average (µM) | Standard Error |
|---|---|---|---|---|---|---|
| AKT3-NL | 0.169 | 0.017 | 0.319 | 0.173 | 0.170 | 0.062 |
| Myr-AKT3-NL | 0.080 | 0.013 | 0.078 | 0.379 | 0.138 | 0.082 |
| Myr-AKT3-T305A-NL | 0.041 | 0.015 | 0.093 | | 0.050 | 0.023 |
| Myr-AKT3-S473A-NL | 0.017 | 0.014 | 0.052 | | 0.028 | 0.012 |
| Myr-AKT3-T305A-S473A-NL | 0.079 | 0.015 | 0.127 | | 0.074 | 0.032 |
| Myr-AKT3-C342S-NL | 0.115 | 0.014 | 0.114 | | 0.080 | 0.034 |

| Capivasertib | Replicate 1 (µM) | Replicate 2 (µM) | Replicate 3 (µM) | Replicate 4 (µM) | Replicate 5 (µM) | Replicate 6 (µM) | Average (µM) | Standard Error |
|---|---|---|---|---|---|---|---|---|
| AKT3-NL | 0.562 | 0.998 | 0.598 | 0.250 | | | 0.602 | 0.154 |
| Myr-AKT3-NL | 0.164 | 0.097 | 15.770 | 0.059 | 0.056 | 0.892 | 2.840 | 2.589 |
| Myr-AKT3-T305A-NL | 0.027 | 0.094 | 0.049 | | | | 0.057 | 0.020 |
| Myr-AKT3-S473A-NL | 0.053 | 0.061 | 0.074 | | | | 0.063 | 0.006 |
| Myr-AKT3-T305A-S473A-NL | 0.251 | 0.139 | 0.159 | | | | 0.183 | 0.035 |
| Myr-AKT3-C342S-NL | 0.360 | 0.166 | 0.109 | | | | 0.212 | 0.076 |

| A-443654 | Replicate 1 (µM) | Replicate 2 (µM) | Replicate 3 (µM) | Replicate 4 (µM) | Average (µM) | Standard Error |
|---|---|---|---|---|---|---|
| AKT3-NL | 0.041 | 0.047 | 0.084 | 0.070 | 0.061 | 0.010 |
| Myr-AKT3-NL | 0.030 | 0.031 | 0.062 | 0.032 | 0.039 | 0.008 |
| Myr-AKT3-T305A-NL | 0.027 | 0.016 | 0.053 | | 0.032 | 0.011 |
| Myr-AKT3-S473A-NL | 0.018 | 0.015 | 0.032 | | 0.022 | 0.006 |
| Myr-AKT3-T305A-S473A-NL | 0.050 | 0.136 | 0.049 | | 0.078 | 0.029 |
| Myr-AKT3-C342S-NL | 0.114 | 0.023 | 0.052 | | 0.063 | 0.027 |

| MK-2206 | Replicate 1 (µM) | Replicate 2 (µM) | Replicate 3 (µM) | Average (µM) | Standard Error |
|---|---|---|---|---|---|
| AKT3-NL | 0.0108 | 0.0004 | 0.1361 | 0.0491 | 0.0436 |
| Myr-AKT3-NL | 0.231 | 0.065 | 0.025 | 0.107 | 0.063 |
| Myr-AKT3-T305A-NL | 0.306 | 0.091 | 1.038 | 0.478 | 0.287 |
| Myr-AKT3-S473A-NL | 0.571 | 0.314 | 0.583 | 0.489 | 0.088 |
| Myr-AKT3-T305A-S473A-NL | 0.847 | 0.453 | 0.535 | 0.612 | 0.120 |
| Myr-AKT3-C342S-NL | 0.285 | 0.008 | 0.425 | 0.239 | 0.123 |

| Miransertib | Replicate 1 (µM) | Replicate 2 (µM) | Replicate 3 (µM) | Replicate 4 (µM) | Replicate 5 (µM) | Average (µM) | Standard Error |
|---|---|---|---|---|---|---|---|
| AKT3-NL | 8.36 | 1.41 | 92.45 | 3.17 | 11.74 | 23.42 | 17.35 |
| Myr-AKT3-NL | 0.203 | 0.013 | 11.020 | 0.024 | | 2.815 | 2.735 |
| Myr-AKT3-T305A-NL | 0.279 | 0.489 | 0.989 | | | 0.586 | 0.21 |
| Myr-AKT3-S473A-NL | 0.790 | 0.195 | 1.198 | | | 0.727 | 0.291 |
| Myr-AKT3-T305A-S473A-NL | 0.337 | 0.728 | 0.585 | | | 0.550 | 0.114 |
| Myr-AKT3-C342S-NL | 0.578 | 0.688 | 1.134 | | | 0.800 | 0.170 |

| Inhibitor VIII | Replicate 1 (µM) | Replicate 2 (µM) | Replicate 3 (µM) | Average (µM) | Standard Error |
|---|---|---|---|---|---|
| AKT3-NL | 5.56 | 3.19 | 5.95 | 4.90 | 0.86 |
| Myr-AKT3-NL | 2.53 | 3.59 | 2.92 | 3.01 | 0.31 |
| Myr-AKT3-T305A-NL | 3.12 | 1.40 | 2.98 | 2.50 | 0.55 |
| Myr-AKT3-S473A-NL | 2.64 | 2.00 | 3.94 | 2.86 | 0.57 |
| Myr-AKT3-T305A-S473A-NL | 2.07 | 1.40 | 3.66 | 2.37 | 0.67 |
| Myr-AKT3-C342S-NL | 2.41 | 3.43 | 2.73 | 2.86 | 0.30 |

L

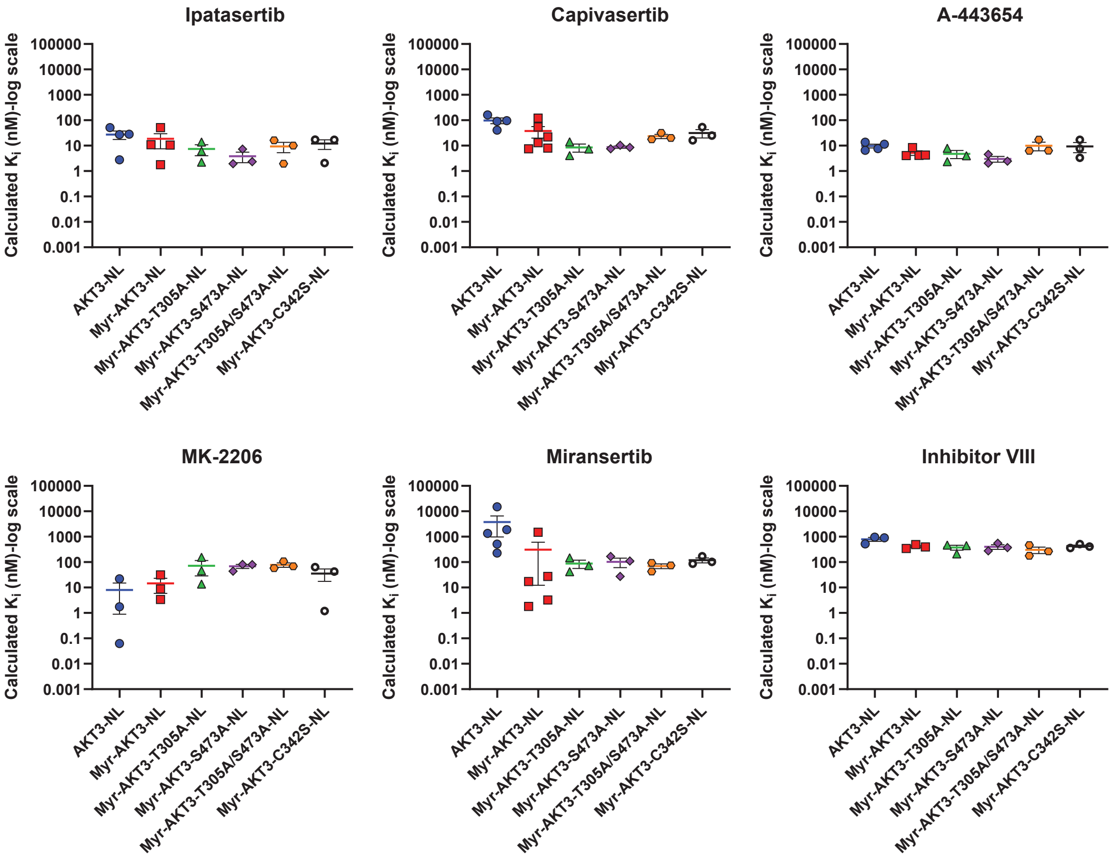


M

#### Calculated $K_i$ Values

| Ipatasertib | Replicate 1 (nM) | Replicate 2 (nM) | Replicate 3 (nM) | Replicate 4 (nM) | Average (nM) | Standard Error |
|---|---|---|---|---|---|---|
| AKT3-NL | 27.42 | 2.76 | 51.66 | 28.01 | 27.46 | 9.98 |
| Myr-AKT3-NL | 10.88 | 1.77 | 10.56 | 51.27 | 18.62 | 11.09 |
| Myr-AKT3-T305A-NL | 6.83 | 2.50 | 15.58 | | 8.31 | 3.85 |
| Myr-AKT3-S473A-NL | 2.69 | 2.21 | 8.26 | | 4.39 | 1.94 |
| Myr-AKT3-T305A-S473A-NL | 10.073 | 1.94 | 16.16 | | 9.39 | 4.12 |
| Myr-AKT3-C342S-NL | 17.12 | 2.07 | 16.95 | | 12.05 | 4.99 |

| Capivasertib | Replicate 1 (nM) | Replicate 2 (nM) | Replicate 3 (nM) | Replicate 4 (nM) | Replicate 5 (nM) | Replicate 6 (nM) | Average (nM) | Standard Error |
|---|---|---|---|---|---|---|---|---|
| AKT3-NL | 90.97 | 161.60 | 96.87 | 40.46 | | | 97.47 | 24.84 |
| Myr-AKT3-NL | 22.20 | 13.08 | 8.04 | 54.95 | 7.51 | 120.85 | 37.77 | 18.12 |
| Myr-AKT3-T305A-NL | 4.53 | 15.81 | 8.25 | | | | 9.53 | 3.32 |
| Myr-AKT3-S473A-NL | 8.52 | 9.68 | 11.86 | | | | 10.02 | 1.00 |
| Myr-AKT3-T305A-S473A-NL | 32.06 | 17.70 | 20.24 | | | | 23.33 | 4.42 |
| Myr-AKT3-C342S-NL | 53.73 | 24.81 | 16.24 | | | | 31.59 | 11.34 |

| A-443654 | Replicate 1 (nM) | Replicate 2 (nM) | Replicate 3 (nM) | Replicate 4 (nM) | Average (nM) | Standard Error |
|---|---|---|---|---|---|---|
| AKT3-NL | 6.58 | 7.66 | 13.67 | 11.32 | 9.80 | 1.63 |
| Myr-AKT3-NL | 4.07 | 4.14 | 8.41 | 4.27 | 5.22 | 1.06 |
| Myr-AKT3-T305A-NL | 4.48 | 2.62 | 8.87 | | 5.32 | 1.85 |
| Myr-AKT3-S473A-NL | 2.79 | 2.33 | 5.16 | | 3.43 | 0.88 |
| Myr-AKT3-T305A-S473A-NL | 6.34 | 17.25 | 6.28 | | 9.96 | 3.65 |
| Myr-AKT3-C342S-NL | 16.94 | 3.35 | 7.68 | | 9.32 | 4.01 |

| MK-2206 | Replicate 1 (nM) | Replicate 2 (nM) | Replicate 3 (nM) | Average (nM) | Standard Error |
|---|---|---|---|---|---|
| AKT3-NL | 1.75 | 0.063 | 22.03 | 7.95 | 7.06 |
| Myr-AKT3-NL | 31.30 | 8.76 | 3.39 | 14.48 | 8.55 |
| Myr-AKT3-T305A-NL | 51.33 | 15.22 | 174.06 | 80.20 | 48.07 |
| Myr-AKT3-S473A-NL | 91.09 | 50.06 | 93.12 | 78.09 | 14.03 |
| Myr-AKT3-T305A-S473A-NL | 107.88 | 57.63 | 68.1 | 77.89 | 15.30 |
| Myr-AKT3-C342S-NL | 42.50 | 1.20 | 63.40 | 35.70 | 18.27 |

| Miransertib | Replicate 1 (nM) | Replicate 2 (nM) | Replicate 3 (nM) | Replicate 4 (nM) | Replicate 5 (nM) | Average (nM) | Standard Error |
|---|---|---|---|---|---|---|---|
| AKT3-NL | 1353.07 | 227.94 | 14966.68 | 512.54 | 1900.58 | 3792.16 | 2809.42 |
| Myr-AKT3-NL | 27.46 | 1.82 | 1492.62 | 3.26 | 17.19 | 308.47 | 296.08 |
| Myr-AKT3-T305A-NL | 46.78 | 82.01 | 165.76 | | | 98.18 | 35.28 |
| Myr-AKT3-S473A-NL | 126.07 | 31.07 | 191.25 | | | 116.13 | 46.51 |
| Myr-AKT3-T305A-S473A-NL | 42.84 | 92.68 | 74.45 | | | 70.00 | 14.56 |
| Myr-AKT3-C342S-NL | 86.15 | 102.60 | 169.10 | | | 119.28 | 25.36 |

| Inhibitor VIII | Replicate 1 (nM) | Replicate 2 (nM) | Replicate 3 (nM) | Average (nM) | Standard Error |
|---|---|---|---|---|---|
| AKT3-NL | 900.75 | 516.91 | 963.73 | 793.80 | 139.63 |
| Myr-AKT3-NL | 343.09 | 486.39 | 395.23 | 408.24 | 41.88 |
| Myr-AKT3-T305A-NL | 523.34 | 235.26 | 500.03 | 419.55 | 92.39 |
| Myr-AKT3-S473A-NL | 420.65 | 318.64 | 629.46 | 456.25 | 91.47 |
| Myr-AKT3-T305A-S473A-NL | 263.05 | 178.13 | 465.62 | 302.27 | 85.28 |
| Myr-AKT3-C342S-NL | 359.52 | 511.77 | 407.67 | 426.33 | 44.93 |

**Figure S4**- Replicate data associated with Figure 4. **(A)** Signal build-up curves and fitted $EC_{50}$ values using the K5 tracer on 293T cells transfected with AKT1-NL, AKT2-NL, and AKT3-NL mutants. **(B)** Replicate binding curves for AKT inhibitors on AKT1-NL, Myr-AKT1-NL, Myr-AKT1-T308A-NL, Myr-AKT1-S473A-NL, Myr-AKT1-T308A/S473A-NL, and Myr-AKT1-C344S-NL mutants. **(C)** Fitted apparent $IC_{50}$ values for curves in (B). **(D)** Plotted calculated $K_i$ values derived from $EC_{50}$ and $IC_{50}$ values in (A and Figure S1A) and (C). **(E)** $K_i$ values plotted in (D). **(F)** Replicate binding curves for AKT inhibitors on AKT2-NL, Myr-AKT2-NL, Myr-AKT2-T309A-NL, Myr-AKT2-S474A-NL, Myr-AKT2-T309A/S474A-NL, and Myr-AKT2-C345S-NL mutants. **(G)** Fitted apparent $IC_{50}$ values for curves in (F). **(H)** Plotted Calculated $K_i$ values derived from $EC_{50}$ and $IC_{50}$ values in (A and Figure S1A) and (G). **(I)** $K_i$ values plotted in (H). **(J)** Replicate binding curves for AKT inhibitors on AKT3-NL, Myr-AKT3-NL, Myr-AKT3-T305A-NL, Myr-AKT3-S473A-NL, Myr-AKT3-T305A/S473A-NL, and Myr-AKT3-C342S-NL mutants. **(K)** Fitted apparent $IC_{50}$ values for curves in (I). **(L)** Plotted calculated $K_i$ values derived from $EC_{50}$ and $IC_{50}$ values in (A and Figure S1A) and (K). **(M)** $K_i$ values plotted in (L).

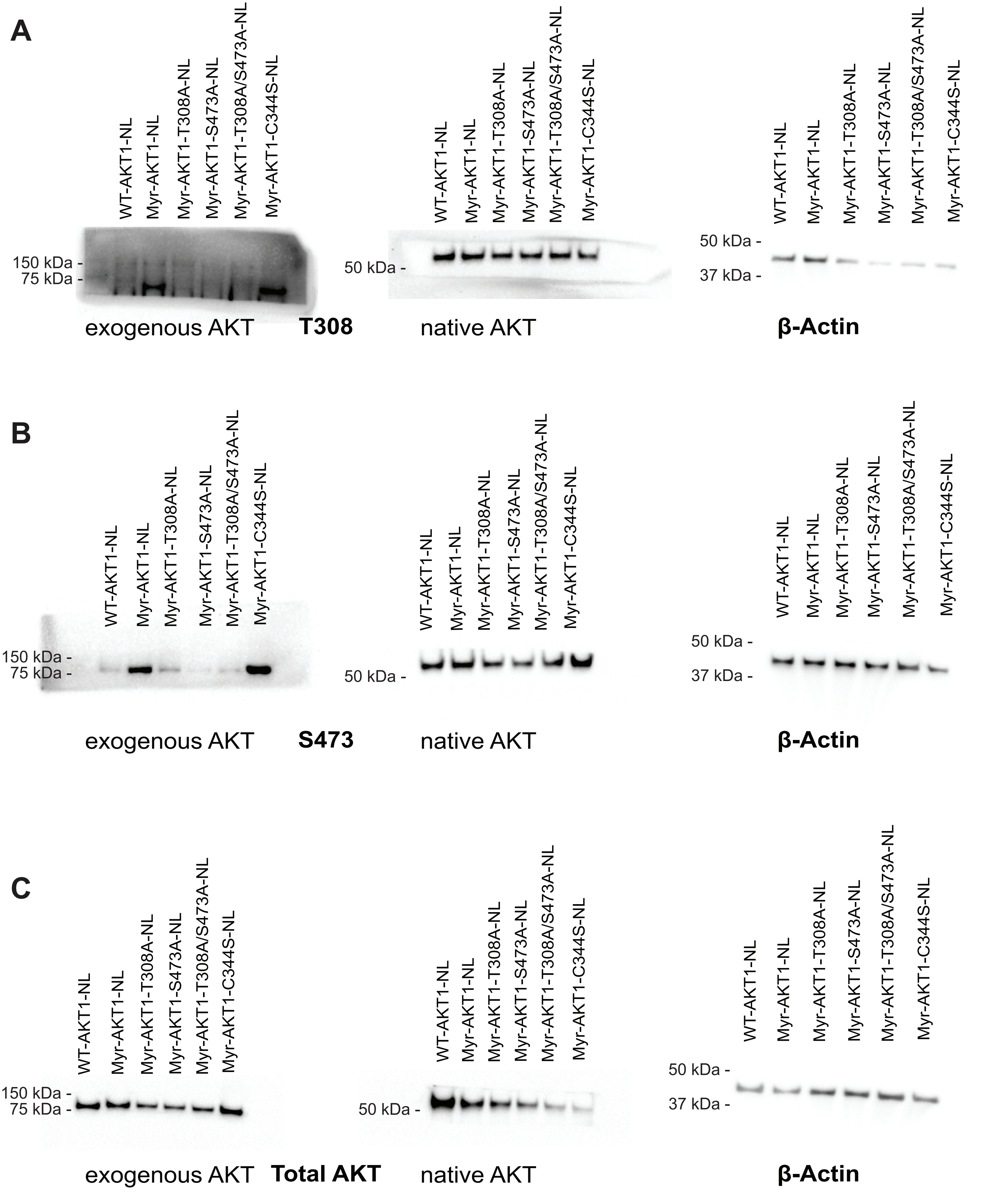


**Figure S5-** Uncropped Blots corresponding to Figure 4. Each set of blots are cut from the same SDS-PAGE gel. **(A)** T308 and β-Actin. **(B)** S473 and β-Actin. **(C)** Total AKT and β-Actin.

A

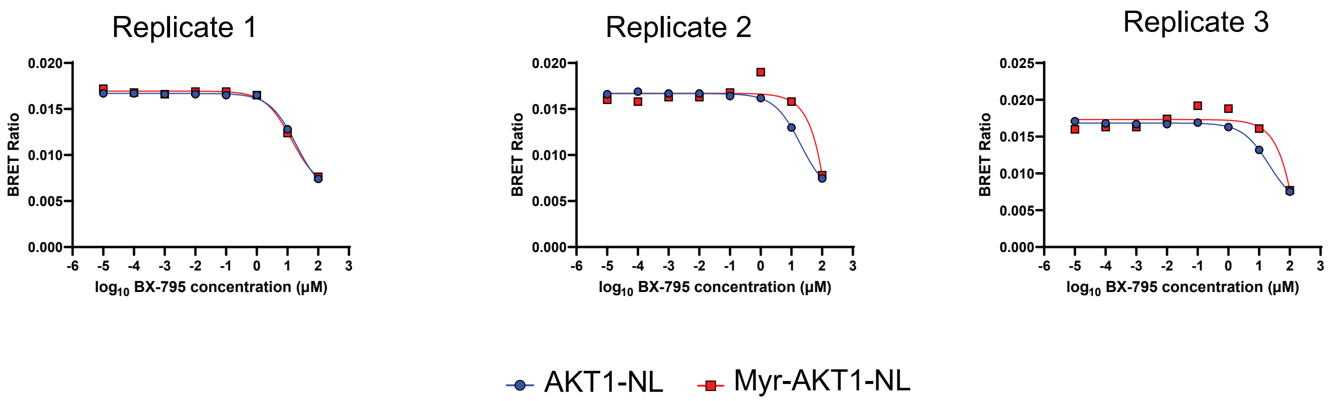


B

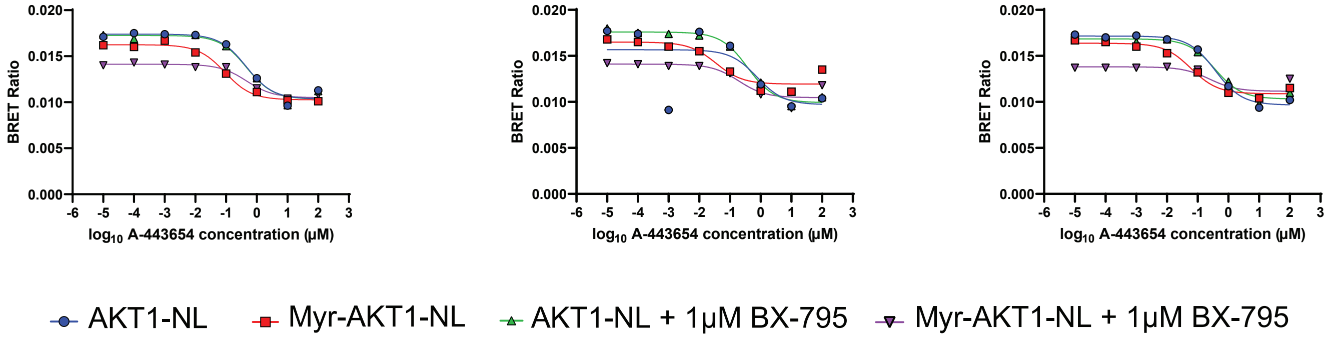


C

Apparent $IC_{50}$ Values

| AKT1-NL | Replicate 1 (μM) | Replicate 2 (μM) | Replicate 3 (μM) | Average (μM) | Standard Error |
|---|---|---|---|---|---|
| BX-795 | 19.0600 | 20.1800 | 20.8300 | 20.0233 | 0.5169 |
| A-443654 | 0.4532 | 0.7665 | 0.3743 | 0.5313 | 0.1198 |
| A-443654 + 1μM BX-795 | 0.459 | 0.372 | 0.364 | 0.398 | 0.030 |

| Myr-AKT1-NL | Replicate 1 (nM) | Replicate 2 (nM) | Replicate 3 (nM) | Average (nM) | Standard Error |
|---|---|---|---|---|---|
| BX-795 | 13.77 | N/A | 1385.00 | 699.39 | 685.62 |
| A-443654 | 0.092 | 0.035 | 0.061 | 0.063 | 0.017 |
| A-443654 + 1μM BX-795 | 0.278 | 0.192 | 0.298 | 0.256 | 0.033 |

D

Calculated $K_i$ Values

| AKT1-NL | Replicate 1 (nM) | Replicate 2 (nM) | Replicate 3 (nM) | Average (nM) | Standard Error |
|---|---|---|---|---|---|
| BX-795 | 3417.30 | 3618.11 | 3734.65 | 3590.02 | 92.68 |
| A-443654 | 81.26 | 137.43 | 67.11 | 95.26 | 21.47 |
| A-443654 + 1μM BX-795 | 82.277 | 66.750 | 65.244 | 71.424 | 5.444 |

| Myr-AKT1-NL | Replicate 1 (nM) | Replicate 2 (nM) | Replicate 3 (nM) | Average (nM) | Standard Error |
|---|---|---|---|---|---|
| BX-795 | 2117.10 | N/A | 212939.88 | 107528.49 | 105411.39 |
| A-443654 | 14.145 | 5.332 | 9.411 | 9.629 | 2.546 |
| A-443654 + 1μM BX-795 | 42.757 | 29.458 | 45.817 | 39.344 | 5.021 |

**Figure S6**- Data associated with Figure 5B and C. **(A)** Dose-response curves of BX-795 on AKT1-NL and Myr-AKT1-NL. **(B)** Dose-response curves of A-443654 in the presence and absence of 1 μM BX-795. **(C)** Fitted apparent $IC_{50}$ values corresponding to panels (A) and (B). **(D)** Calculated Ki values using $IC_{50}$ values from (C) and $EC_{50}$ values from Figure S1A and Figure S4A.

A

AKT1-NL

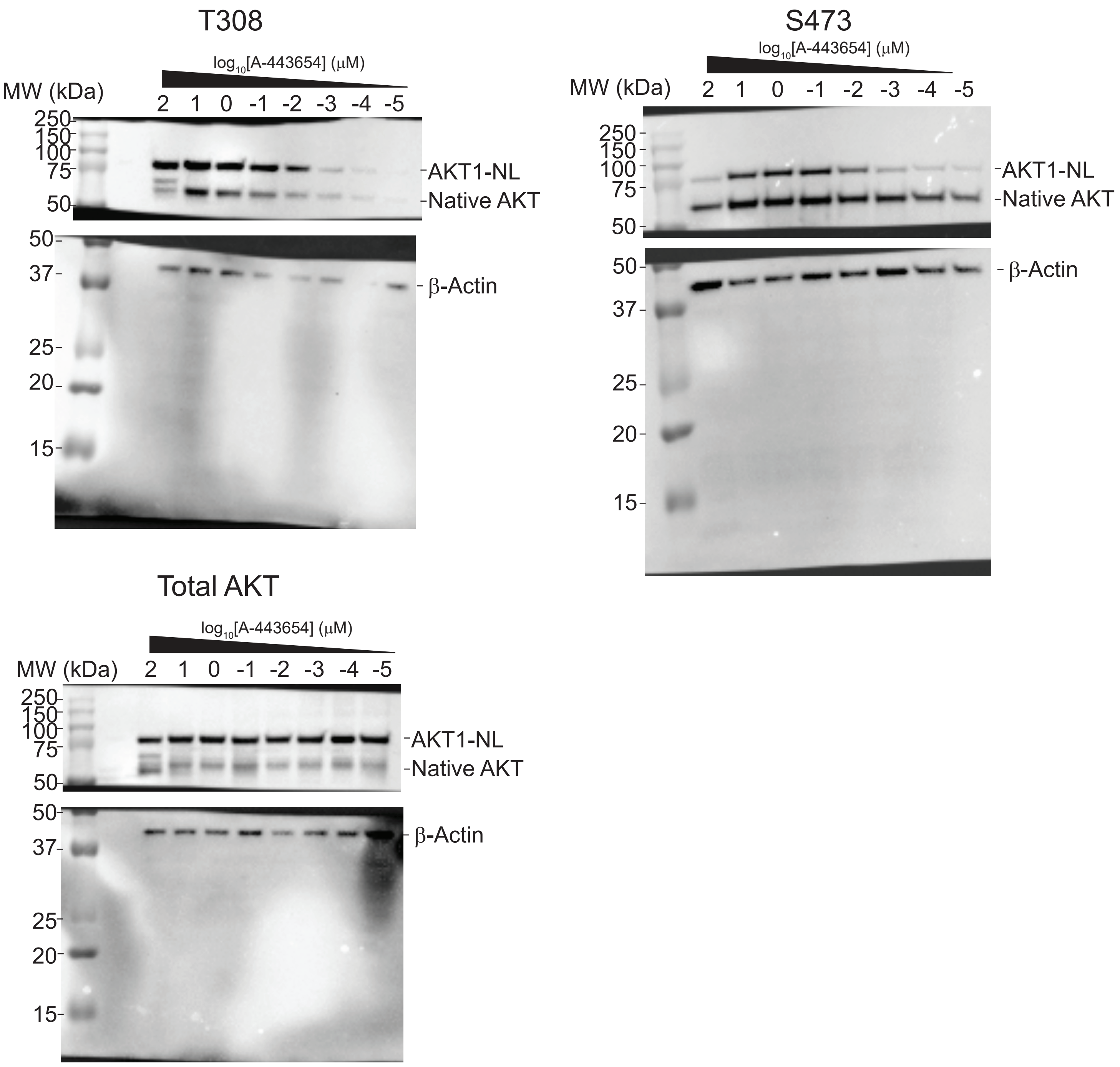
T308
log10[A-443654] (μM)
MW (kDa)
2 1 0 -1 -2 -3 -4 -5
250
150
100
75
50
AKT1-NL
Native AKT
50
37
β-Actin
25
20
15
S473
log10[A-443654] (μM)
MW (kDa)
2 1 0 -1 -2 -3 -4 -5
250
150
100
75
50
AKT1-NL
Native AKT
50
β-Actin
37
25
20
15
Total AKT
log10[A-443654] (μM)
MW (kDa)
2 1 0 -1 -2 -3 -4 -5
250
150
100
75
50
AKT1-NL
Native AKT
50
37
β-Actin
25
20
15

B

### AKT1-NL + 1 μM BX-795

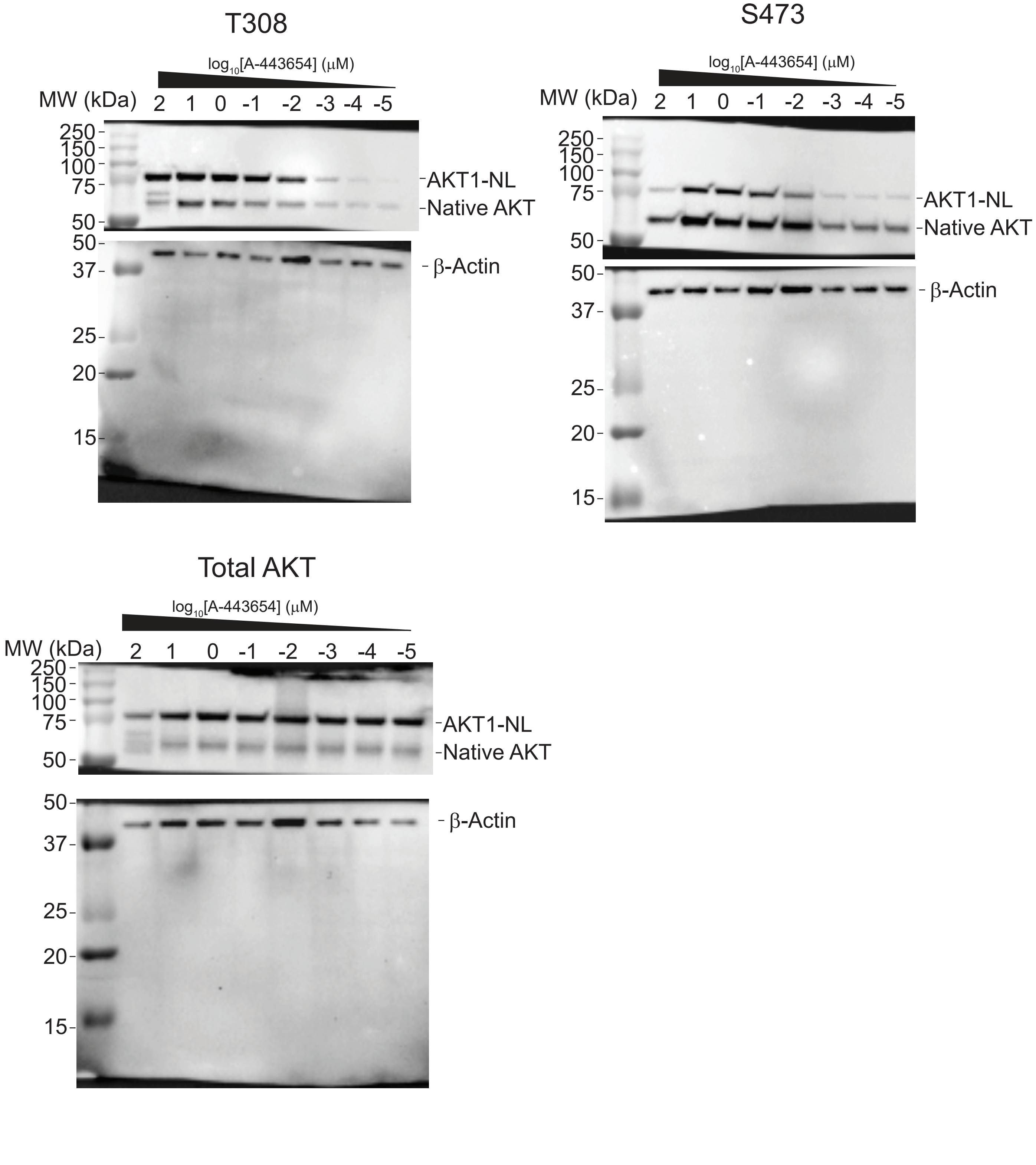

C

### Myr-AKT1-NL

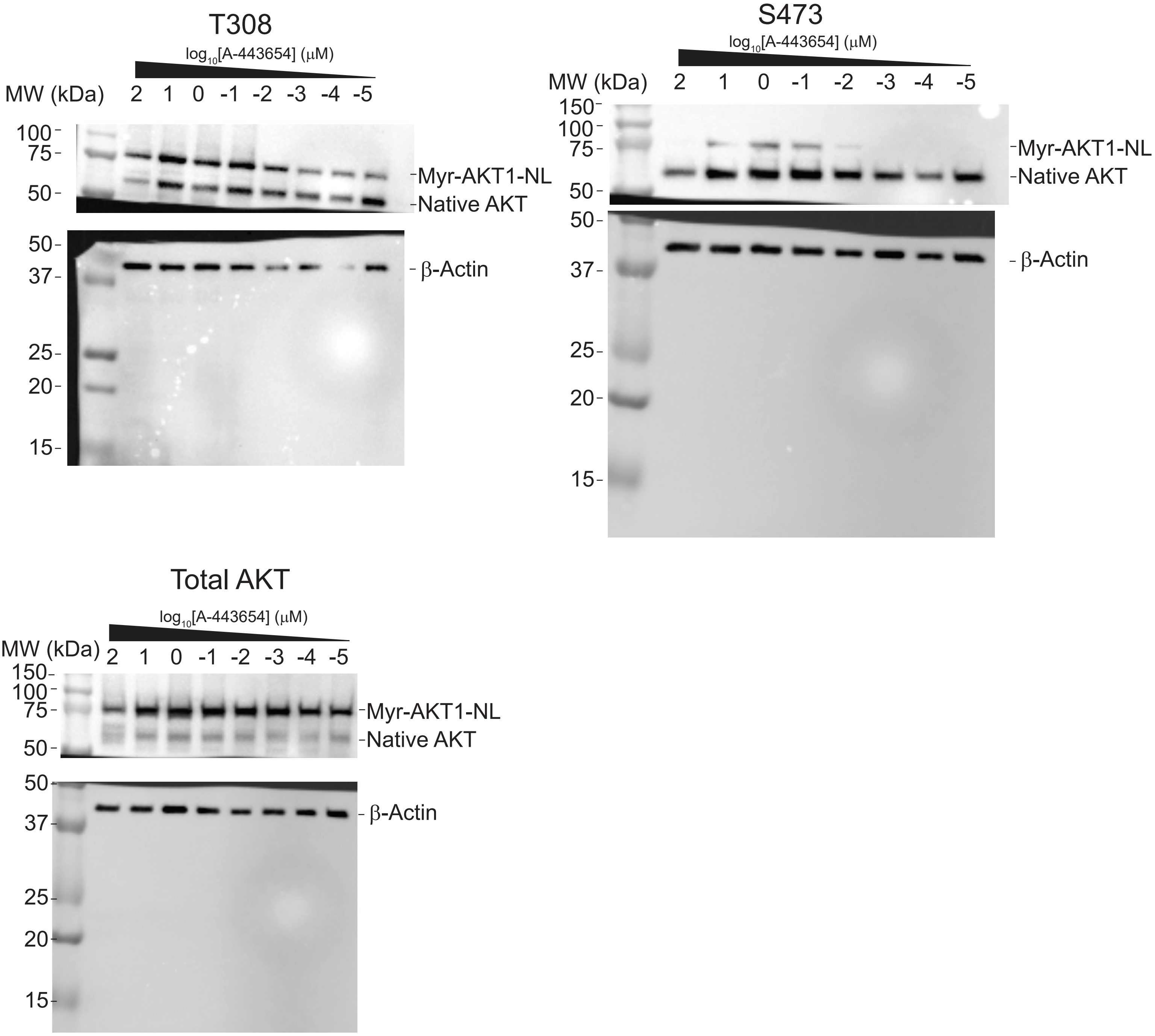
T308
log10[A-443654] (μM)
MW (kDa)
2 1 0 -1 -2 -3 -4 -5
100
75
50
Myr-AKT1-NL
Native AKT
50
37
25
20
15
β-Actin
S473
log10[A-443654] (μM)
MW (kDa)
2 1 0 -1 -2 -3 -4 -5
150
100
75
50
Myr-AKT1-NL
Native AKT
50
37
25
20
15
β-Actin
Total AKT
log10[A-443654] (μM)
MW (kDa)
2 1 0 -1 -2 -3 -4 -5
150
100
75
50
Myr-AKT1-NL
Native AKT
50
37
25
20
15
β-Actin

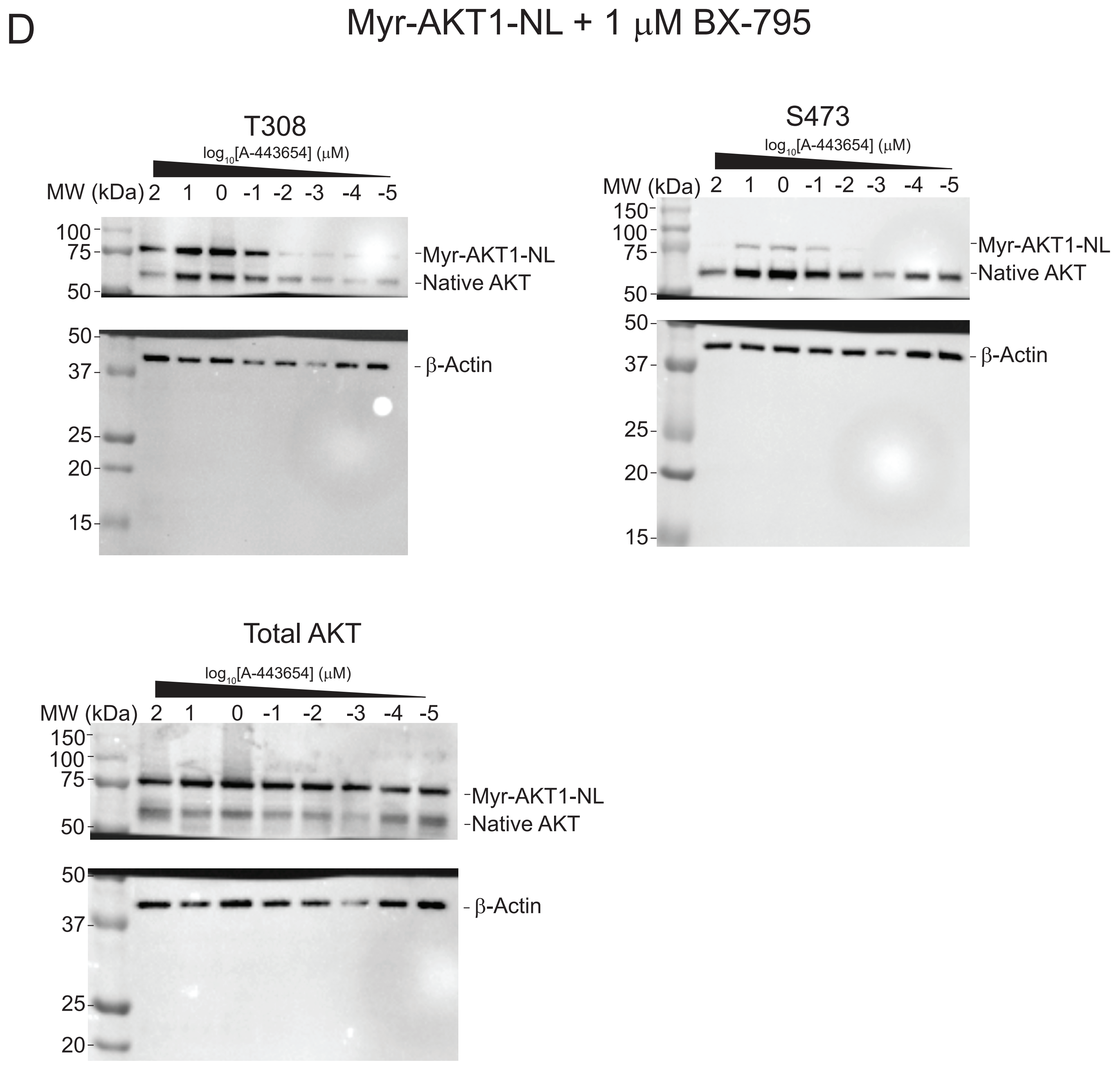


**Figure S7**- Uncropped blots associated with Figure 5D and E. Blots indicate T308 and S473 phosphorylation, and Total AKT in the presence of a A-443654 dose-response curves. Each blot has an accompanying β-actin blot associated with the same SDS-PAGE gel. **(A)** Over-expressed AKT1-NL. **(B)** Over-expressed AKT1-NL in the presence of 1 μM BX-795. **(C)** Over-expressed Myr-AKT1-NL. **(D)** Over-expressed Myr-AKT1-NL in the presence of 1 μM BX-795.

A

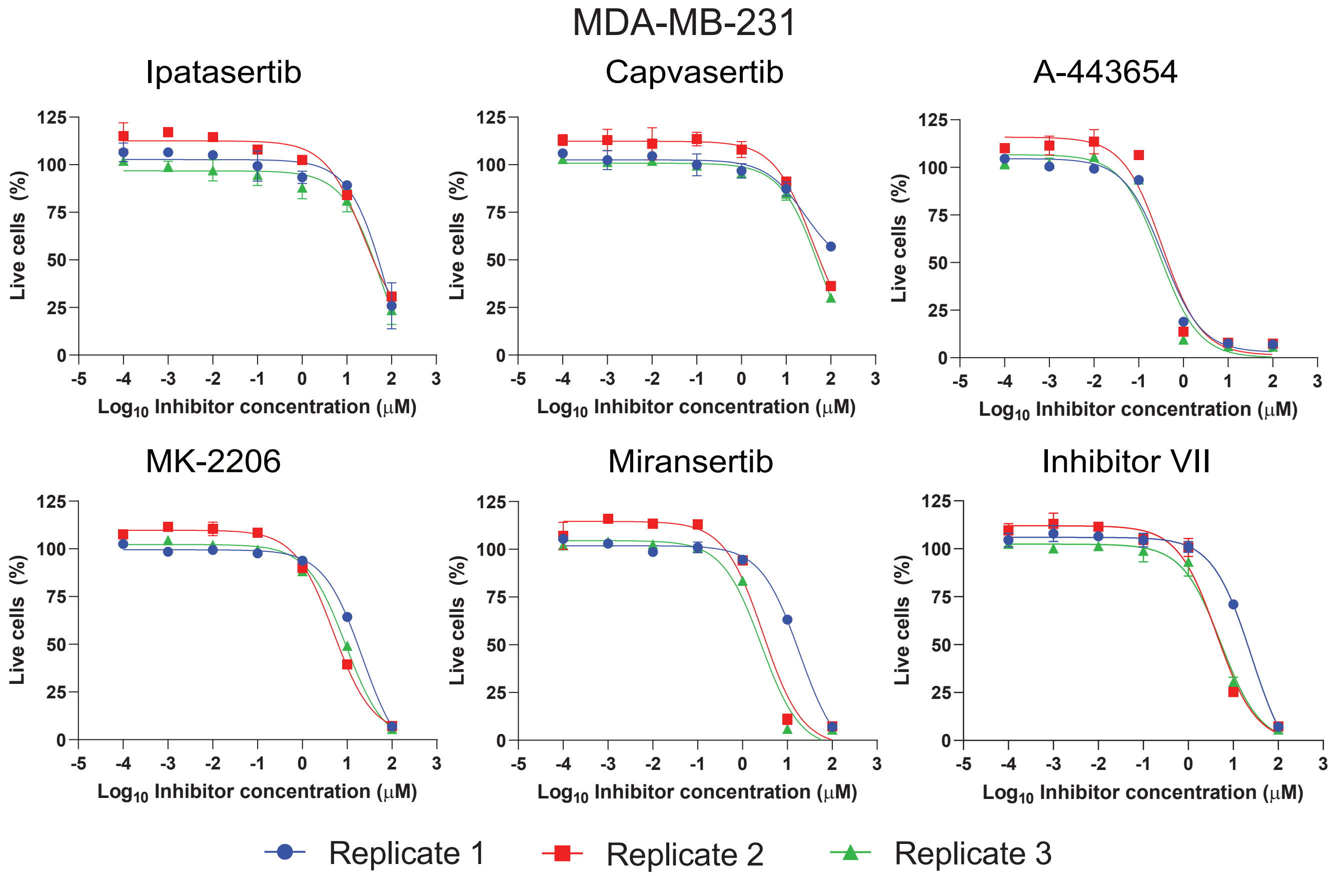


$IC_{50}$ values

| MDA-MB-231 | | | | | |
|---|---|---|---|---|---|
| Drugs | Replicate 1 (μM) | Replicate 2 (μM) | Replicate 3 (μM) | Average (μM) | Standard Error |
| Ipatasertib | 90.31 | 24.03 | 60.19 | 58.18 | 19.16 |
| Capivasertib | 25.71 | 39.38 | 57.57 | 40.89 | 9.23 |
| A-443654 | 0.35 | 0.34 | 0.30 | 0.33 | 0.01 |
| MK-2206 | 21.86 | 4.96 | 9.27 | 12.03 | 5.07 |
| Miransertib | 18.79 | 2.89 | 2.62 | 8.10 | 5.35 |
| Inhibitor VIII | 25.28 | 4.28 | 5.37 | 11.64 | 6.83 |

B

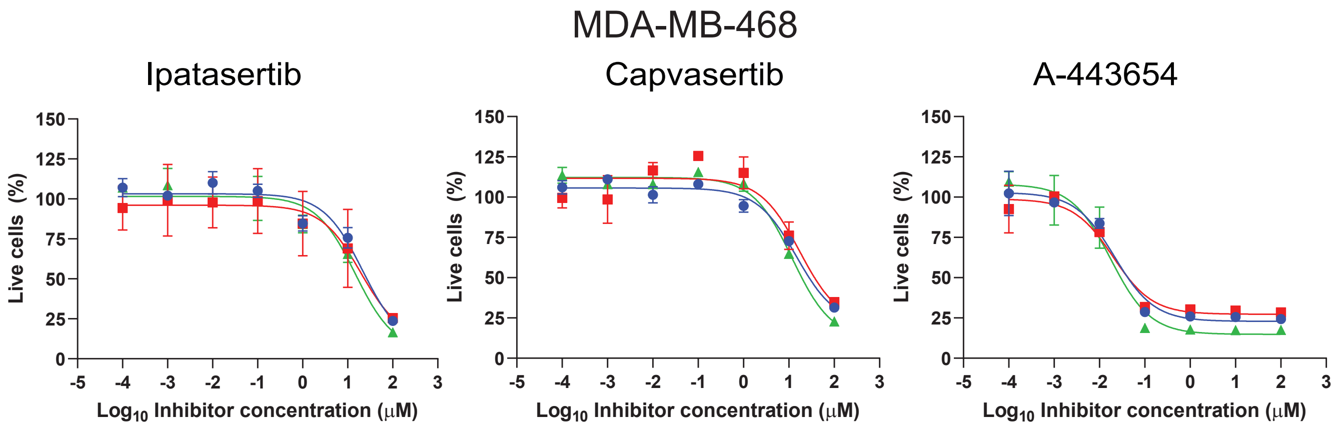

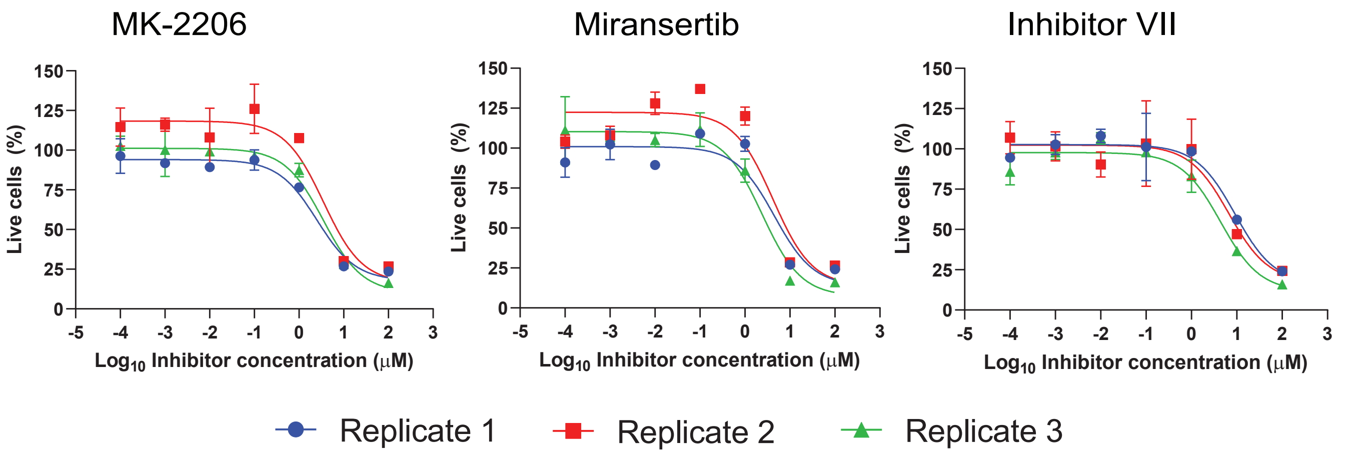


IC$_{50}$ values

| MDA-MB-468 | | | | | |
|---|---|---|---|---|---|
| Drugs | Replicate 1 (μM) | Replicate 2 (μM) | Replicate 3 (μM) | Average (μM) | Standard Error |
| Ipatasertib | 21.50 | 19.33 | 14.80 | 18.54 | 1.97 |
| Capivasertib | 14.61 | 17.28 | 11.84 | 14.58 | 1.57 |
| A-443654 | 0.022 | 0.020 | 0.018 | 0.020 | 0.001 |
| MK-2206 | 2.53 | 3.53 | 3.53 | 3.20 | 0.33 |
| Miransertib | 4.56 | 4.19 | 2.39 | 3.71 | 0.67 |
| Inhibitor VIII | 9.21 | 6.71 | 4.41 | 6.78 | 1.39 |

**Figure S8-** Replicate Cell Viability Curves and fitted IC$_{50}$ values for MDA-MB2-31 **(A)** and MDA-MB-468 **(B)** cell lines. Data correspond to Figure 6A.

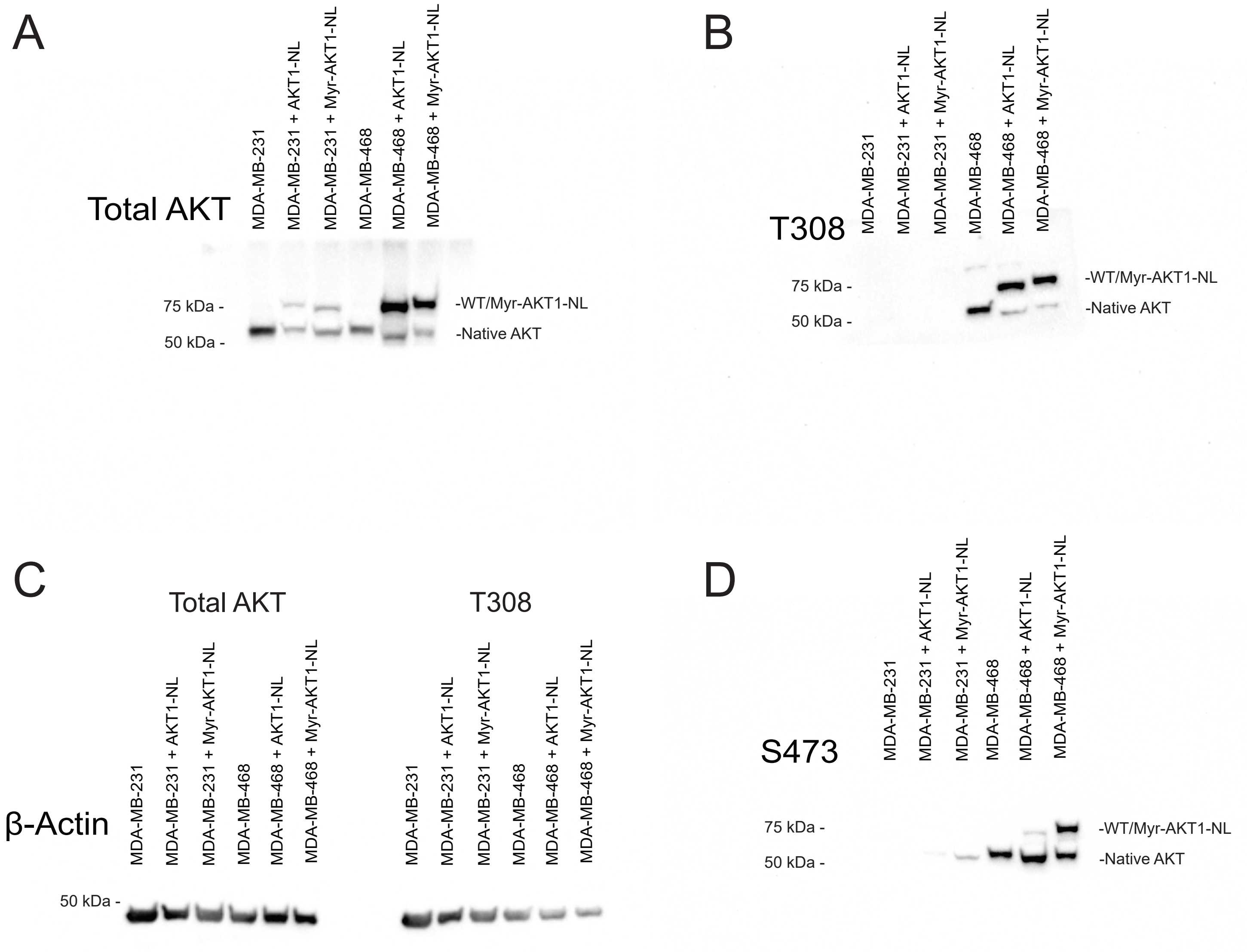

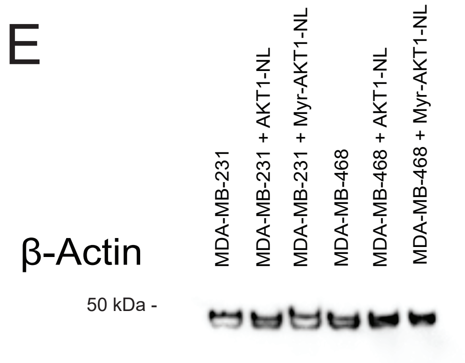


**Figure S9-**Uncropped blots for Figure 6B.
Total AKT **(A)**. T308 phosphorylaton **(B)**. β-Actin corresponding to Total AKT and T308 blots **(C)**.
S473 phosphorylation (D). β-Actin corresponding to the S473 blot **(E)**.

A

#### AKT1-NL

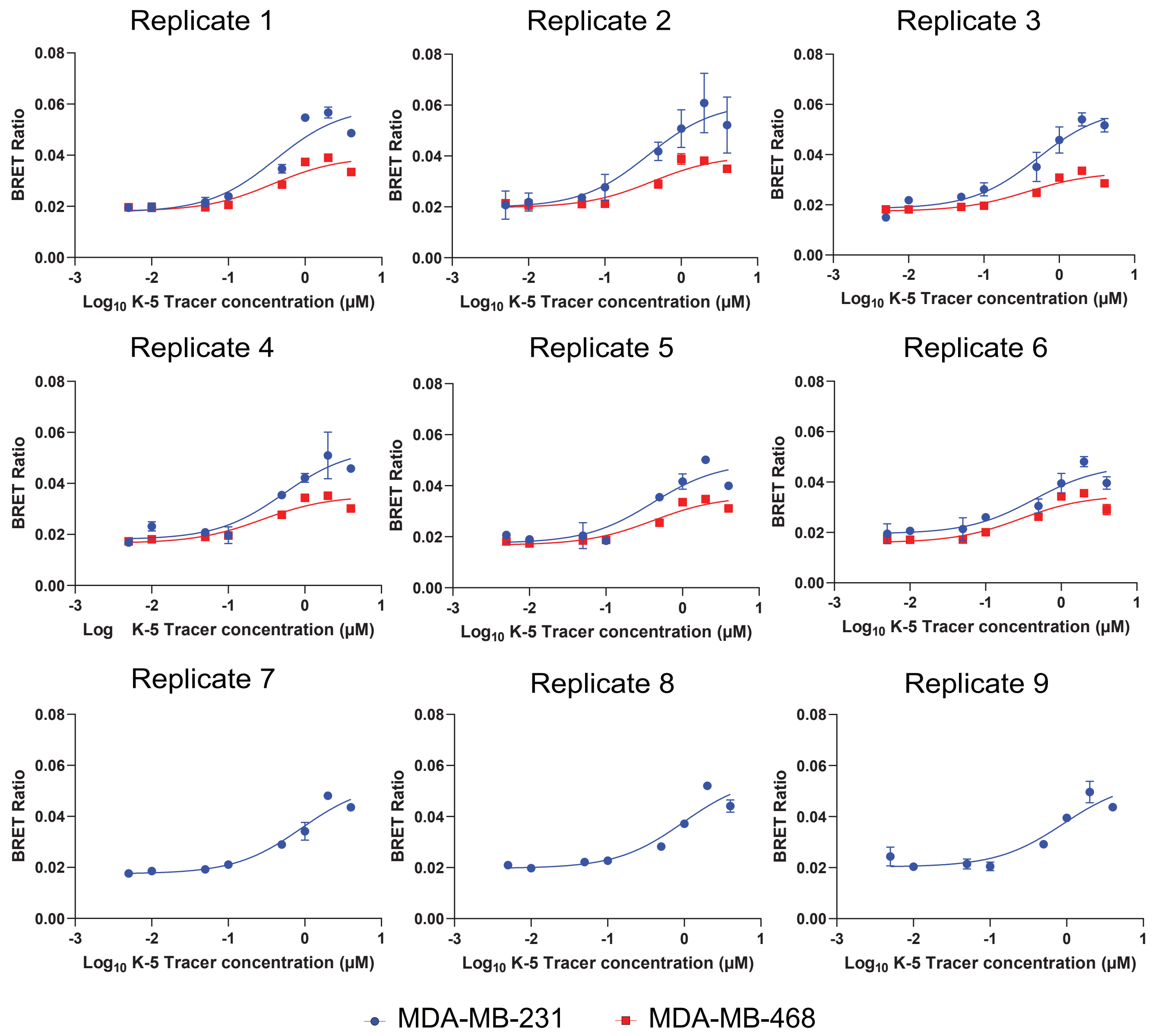


| AKT1-NL | Replicate 1 (μM) | Replicate 2 (μM) | Replicate 3 (μM) | Replicate 4 (μM) | Replicate 5 (μM) | Replicate 6 (μM) | Replicate 7 (μM) | Replicate 8(μM) | Replicate 9 (μM) | Average (μM) | Standard Error |
|---|---|---|---|---|---|---|---|---|---|---|---|
| MDA-MB-231 | 0.41 | 0.37 | 0.52 | 0.53 | 0.41 | 0.44 | 0.91 | 0.97 | 0.91 | 0.61 | 0.08 |
| MDA-MB-468 | 0.38 | 0.41 | 0.38 | 0.28 | 0.28 | 0.29 | | | | 0.33 | 0.03 |

#### Myr-AKT1-NL

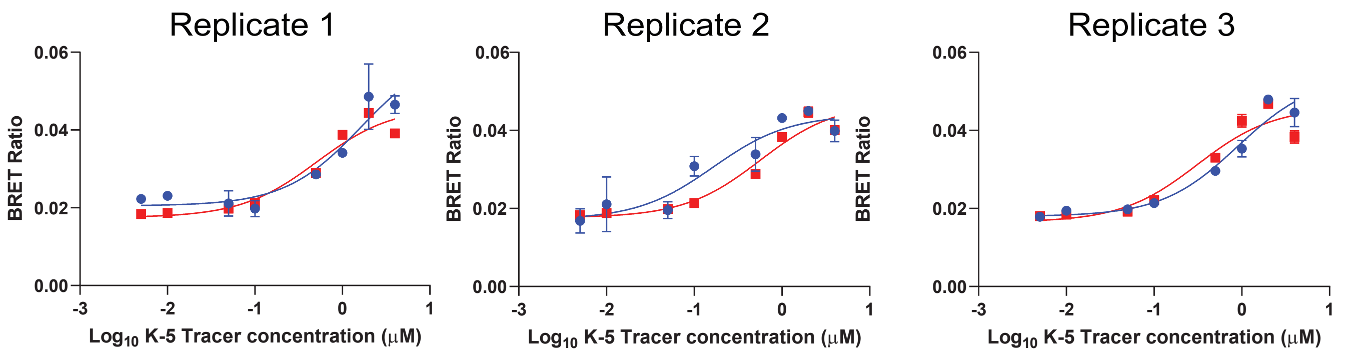

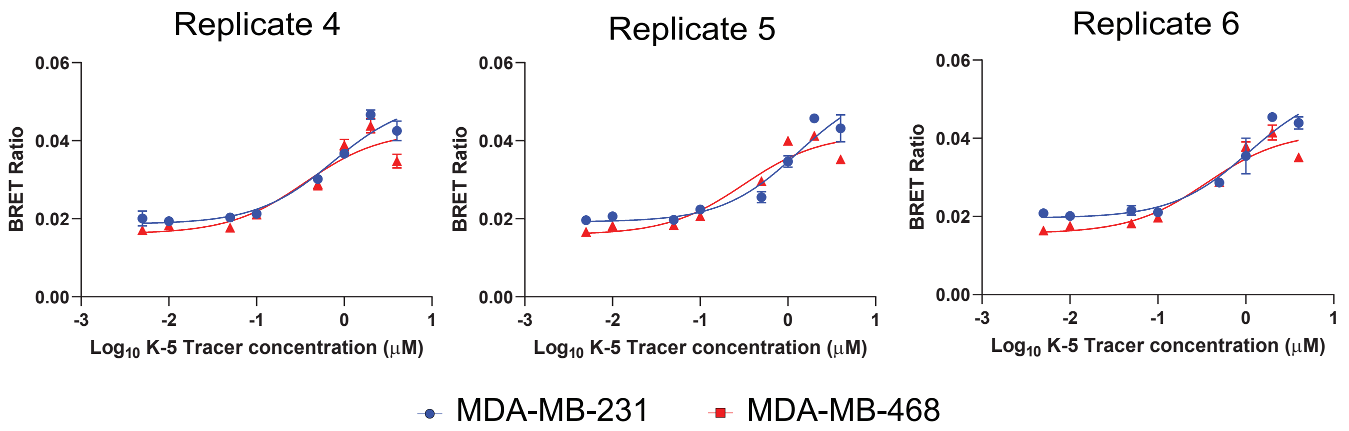


| Myr-AKT1-NL | Replicate 1 (μM) | Replicate 2 (μM) | Replicate 3 (μM) | Replicate 4 (μM) | Replicate 5 (μM) | Replicate 6 (μM) | Average (μM) | Standard Error |
|---|---|---|---|---|---|---|---|---|
| MDA-MB-231 | 1.63 | 0.16 | 0.90 | 0.74 | 1.29 | 1.09 | 0.97 | 0.20 |
| MDA-MB-468 | 0.51 | 0.57 | 0.31 | 0.36 | 0.31 | 0.36 | 0.40 | 0.04 |

B

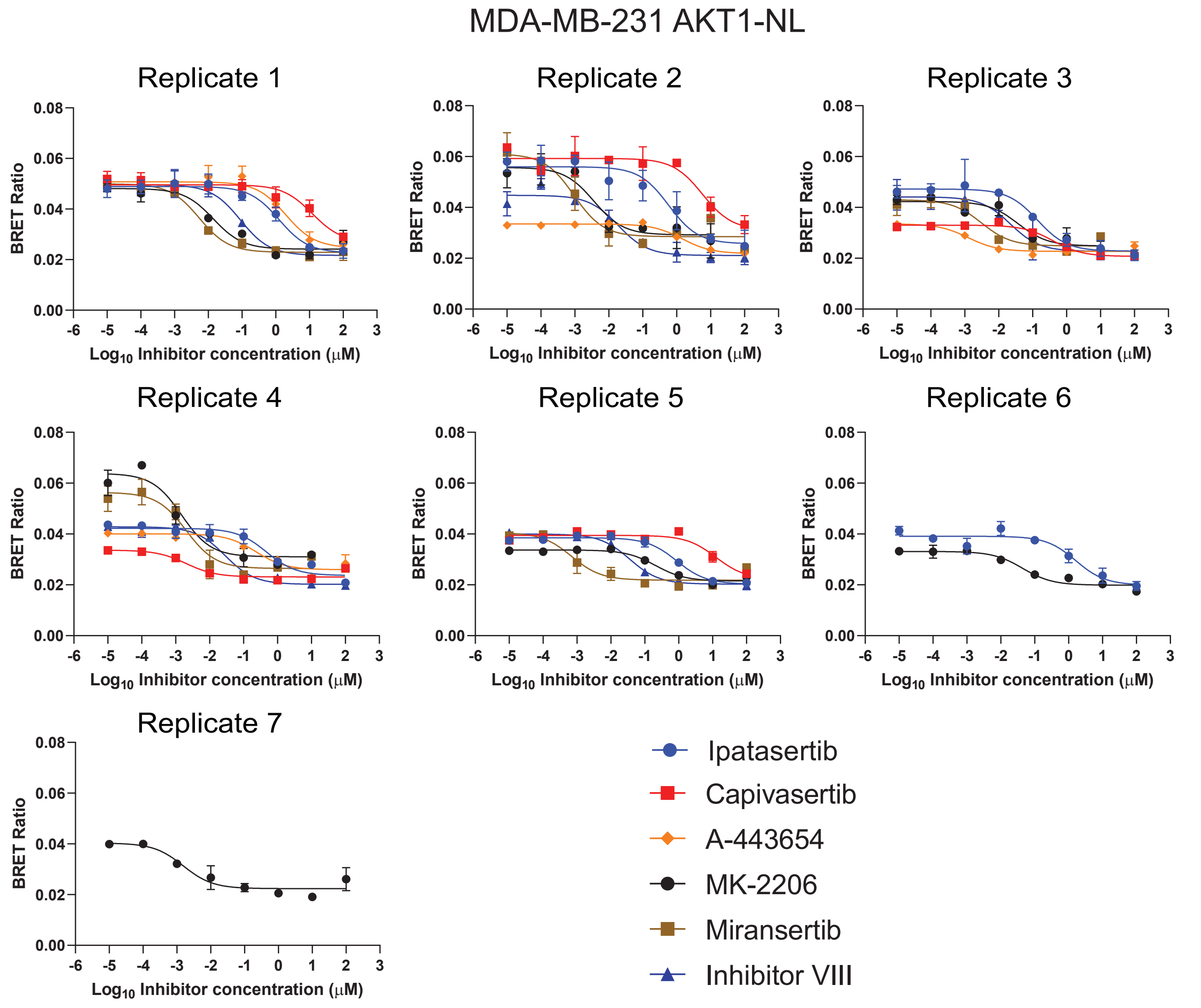

#### Apparent $IC_{50}$ values

| MDA-MB-231 | | | | | | | | | |
|---|---|---|---|---|---|---|---|---|---|
| AKT1-NL | Replicate 1 (μM) | Replicate 2 (μM) | Replicate 3 (μM) | Replicate 4 (μM) | Replicate 5 (μM) | Replicate 6 (μM) | Replicate 7 (μM) | Average (μM) | Standard Error |
| Ipatasertib | 1.26 | 0.58 | 0.14 | 0.47 | 0.88 | 1.63 | | 0.83 | 0.22 |
| Capivasertib | 12.050 | 6.422 | 0.400 | 0.002 | 12.570 | | | 6.289 | 2.710 |
| A-443654 | 1.9910 | 1.4560 | 0.0013 | 0.2323 | | | | 0.9201 | 0.4788 |
| MK-2206 | 0.0141 | 0.0036 | 0.0374 | 0.0008 | 0.2076 | 0.0441 | 0.0015 | 0.0442 | 0.0280 |
| Miransertib | 0.0054 | 0.0009 | 0.0031 | 0.0016 | 0.0007 | | | 0.0024 | 0.0009 |
| Inhibitor VIII | 0.0901 | 0.0189 | 0.0223 | 0.0287 | 0.0358 | | | 0.0391 | 0.0131 |

#### Calculated $K_i$

| MDA-MB-231 | | | | | | | | | |
|---|---|---|---|---|---|---|---|---|---|
| AKT1-NL | Replicate 1 (nM) | Replicate 2 (nM) | Replicate 3 (nM) | Replicate 4 (nM) | Replicate 5 (nM) | Replicate 6 (nM) | Replicate 7(nM | Average (nM) | Standard Error |
| Ipatasertib | 475.36 | 219.52 | 52.29 | 177.65 | 330.38 | 616.20 | | 311.90 | 84.38 |
| Capivasertib | 4549.74 | 2424.77 | 150.92 | 0.77 | 4746.08 | | | 2374.45 | 1023.17 |
| A-443654 | 751.75 | 549.74 | 0.47 | 87.71 | | | | 347.42 | 180.79 |
| MK-2206 | 5.31 | 1.37 | 14.13 | 0.30 | 78.38 | 16.65 | 0.58 | 16.68 | 10.58 |
| Miransertib | 2.05 | 0.36 | 1.17 | 0.62 | 0.26 | | | 0.89 | 0.33 |
| Inhibitor VIII | 34.02 | 7.12 | 8.40 | 10.83 | 13.50 | | | 14.77 | 4.93 |

C

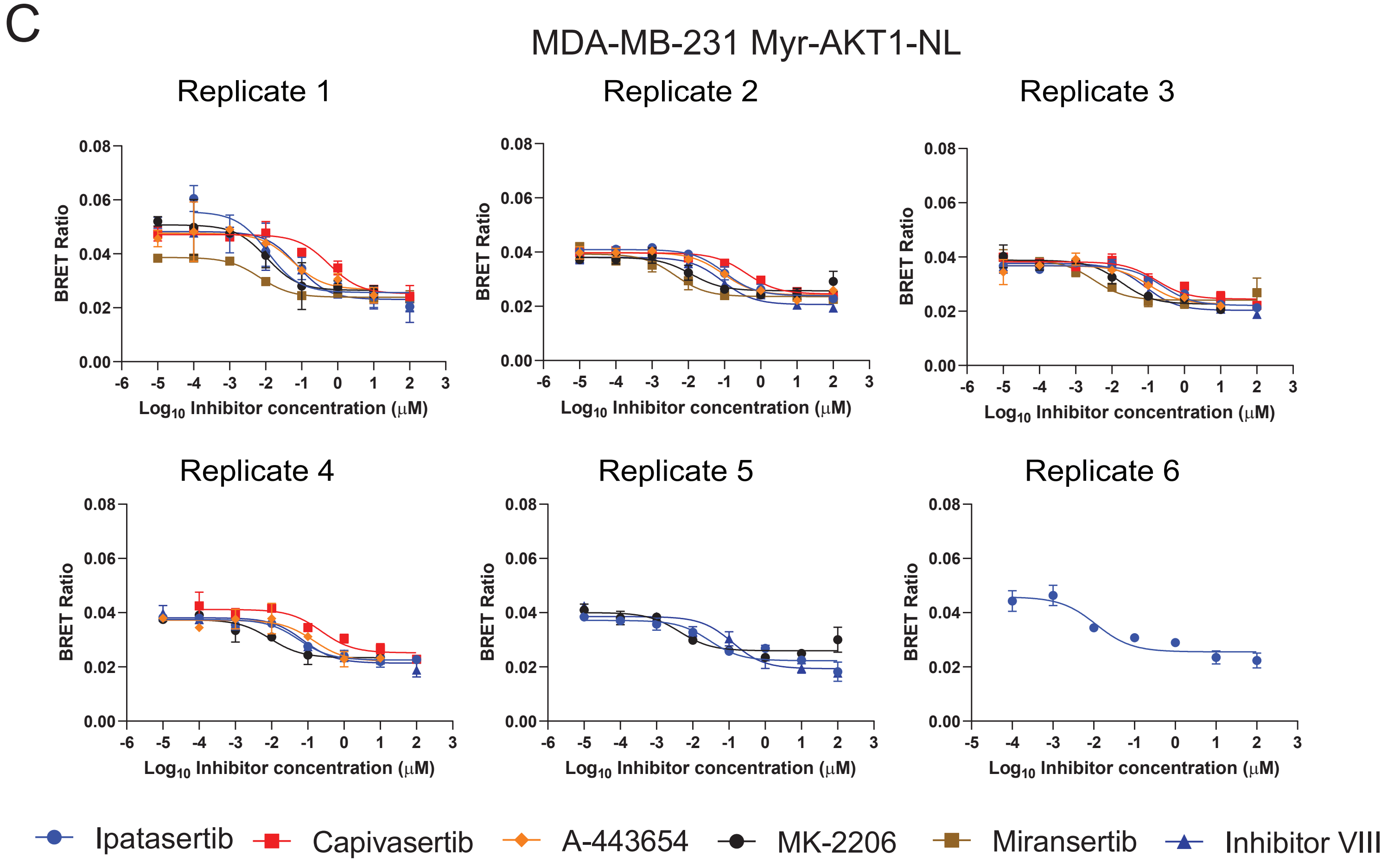

#### Apparent $IC_{50}$ values

| MDA-MB-231 | | | | | | | | |
|---|---|---|---|---|---|---|---|---|
| Myr-AKT1-NL | Replicate 1 (nM) | Replicate 2 (nM) | Replicate 3 (nM) | Replicate 4 (nM) | Replicate 5 (nM) | Replicate 6 (nM) | Average (nM) | Standard Error |
| Ipatasertib | 0.0114 | 0.1013 | 0.2363 | 0.0573 | 0.0290 | 0.0114 | 0.0744 | 0.0352 |
| Capivasertib | 0.5733 | 0.4038 | 0.1711 | 0.2348 | | | 0.3458 | 0.0904 |
| A-443654 | 0.0557 | 0.0920 | 0.1036 | 0.1454 | | | 0.0992 | 0.0185 |
| MK-2206 | 0.0105 | 0.0119 | 0.0178 | 0.0092 | 0.0043 | | 0.0108 | 0.0022 |
| Miransertib | 0.0068 | 0.0043 | 0.0032 | | | | 0.0048 | 0.0011 |
| Inhibitor VIII | 0.0757 | 0.0872 | 0.0845 | 0.0755 | 0.1263 | | 0.0898 | 0.0094 |

#### Calculated $K_i$ values

| MDA-MB-231 | | | | | | | | |
|---|---|---|---|---|---|---|---|---|
| Myr-AKT1-NL | Replicate 1 (μM) | Replicate 2 (μM) | Replicate 3 (μM) | Replicate 4 (μM) | Replicate 5 (μM) | Replicate 6 (μM | Average (μM) | Standard Error |
| Ipatasertib | 5.59 | 49.83 | 116.23 | 28.18 | 14.24 | 5.58 | 36.61 | 17.33 |
| Capivasertib | 282.00 | 198.63 | 84.16 | 115.50 | | | 188.26 | 57.35 |
| A-443654 | 27.41 | 45.25 | 50.96 | 71.52 | | | 41.21 | 7.09 |
| MK-2206 | 5.18 | 5.87 | 8.77 | 4.51 | 2.12 | | 5.29 | 1.07 |
| Miransertib | 3.35 | 2.11 | 1.59 | | | | 2.73 | 0.62 |
| Inhibitor VIII | 37.22 | 42.90 | 41.55 | 37.12 | 62.13 | | 44.18 | 4.63 |

D

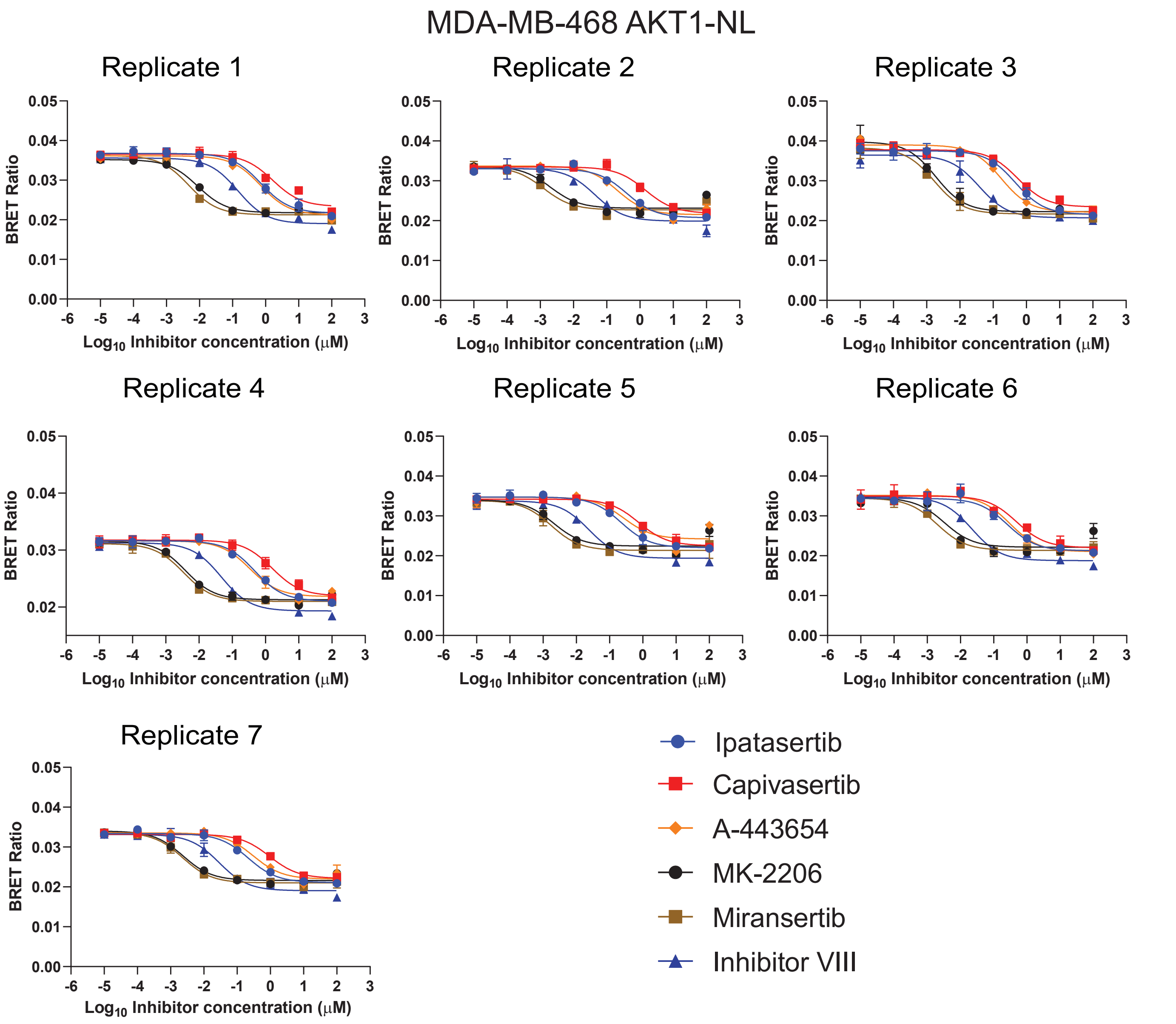

#### Apparent $IC_{50}$ values

| MDA-MB-468 | | | | | | | | | |
|---|---|---|---|---|---|---|---|---|---|
| AKT1-NL | Replicate 1 (μM) | Replicate 2 (μM) | Replicate 3 (μM) | Replicate 4 (μM) | Replicate 5 (μM) | Replicate 6 (μM) | Replicate 7 (μM) | Average (μM) | Standard Error |
| Ipatasertib | 0.714 | 0.400 | 0.468 | 0.478 | 0.221 | 0.259 | 0.216 | 0.394 | 0.068 |
| Capivasertib | 1.657 | 1.456 | 0.550 | 1.563 | 0.751 | 0.489 | 0.949 | 1.059 | 0.186 |
| A-443654 | 0.707 | 0.208 | 0.176 | 0.339 | 0.267 | 0.305 | 0.310 | 0.330 | 0.067 |
| MK-2206 | 0.009 | 0.002 | 0.002 | 0.004 | 0.002 | 0.003 | 0.002 | 0.003 | 0.001 |
| Miransertib | 0.005 | 0.001 | 0.002 | 0.003 | 0.002 | 0.002 | 0.002 | 0.002 | 0.0005 |
| Inhibitor VIII | 0.146 | 0.044 | 0.038 | 0.047 | 0.024 | 0.022 | 0.029 | 0.050 | 0.016 |

#### Calculated $K_i$ values

| MDA-MB-468 | | | | | | | | | |
|---|---|---|---|---|---|---|---|---|---|
| AKT1-NL | Replicate 1 (nM) | Replicate 2 (nM) | Replicate 3 (nM) | Replicate 4 (nM) | Replicate 5 (nM) | Replicate 6(nM) | Replicate 7(nM | Average (nM) | Standard Error |
| Ipatasertib | 187.78 | 105.11 | 122.96 | 125.72 | 58.04 | 68.13 | 56.69 | 103.49 | 17.92 |
| Capivasertib | 435.73 | 382.87 | 144.68 | 411.01 | 197.35 | 128.62 | 249.50 | 278.54 | 49.03 |
| A-443654 | 185.94 | 54.59 | 46.15 | 89.14 | 70.08 | 80.12 | 81.54 | 86.80 | 17.51 |
| MK-2206 | 2.31 | 0.54 | 0.49 | 1.02 | 0.54 | 0.86 | 0.61 | 0.91 | 0.25 |
| Miransertib | 1.24 | 0.33 | 0.44 | 0.90 | 0.46 | 0.52 | 0.60 | 0.64 | 0.12 |
| Inhibitor VIII | 38.50 | 11.60 | 9.97 | 12.29 | 6.21 | 5.74 | 7.53 | 13.12 | 4.34 |

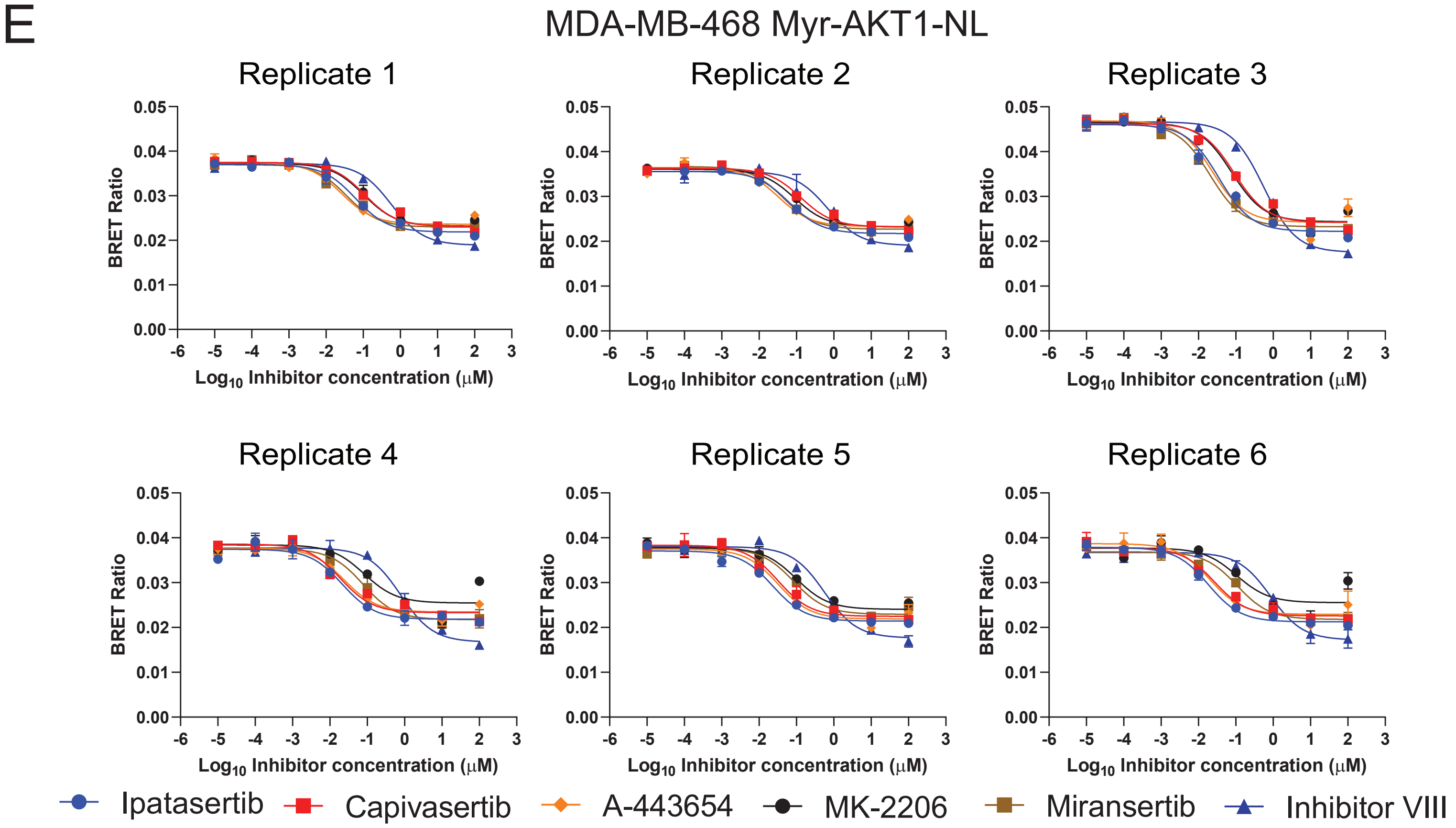


#### Apparent $IC_{50}$ values

| MDA-MB-468 | | | | | | | | |
|---|---|---|---|---|---|---|---|---|
| Myr-AKT1-NL | Replicate 1 (μM) | Replicate 2 (μM) | Replicate 3 (μM) | Replicate 4 (μM) | Replicate 5 (μM) | Replicate 6 (μM) | Average (μM) | Standard Error |
| Ipatasertib | 0.0622 | 0.0636 | 0.0353 | 0.0216 | 0.0234 | 0.0186 | 0.0375 | 0.0084 |
| Capivasertib | 0.1090 | 0.1310 | 0.0904 | 0.0192 | 0.0323 | 0.0266 | 0.0681 | 0.0196 |
| A-443654 | 0.0261 | 0.0330 | 0.0235 | 0.0240 | 0.0320 | 0.0197 | 0.0264 | 0.0021 |
| MK-2206 | 0.1037 | 0.0885 | 0.0772 | 0.0829 | 0.0982 | 0.1046 | 0.0925 | 0.0046 |
| Miransertib | 0.0348 | 0.0444 | 0.0205 | 0.0771 | 0.0839 | 0.1081 | 0.0615 | 0.0137 |
| Inhibitor VIII | 0.5721 | 0.6862 | 0.5597 | 0.7862 | 0.5679 | 0.8550 | 0.6712 | 0.0517 |

Calculated $K_i$ values

| MDA-MB-468 | | | | | | | | |
|---|---|---|---|---|---|---|---|---|
| Myr-AKT1-NL | Replicate 1 (nM) | Replicate 2 (nM) | Replicate 3 (nM) | Replicate 4 (nM) | Replicate 5 (nM) | Replicate 6(nM) | Average (nM) | Standard Error |
| Ipatasertib | 17.88 | 18.28 | 10.14 | 6.21 | 6.71 | 5.33 | 10.76 | 2.41 |
| Capivasertib | 31.31 | 37.63 | 25.97 | 5.50 | 9.28 | 7.64 | 19.56 | 5.63 |
| A-443654 | 7.49 | 9.46 | 6.75 | 6.90 | 9.18 | 5.65 | 7.57 | 0.61 |
| MK-2206 | 29.79 | 25.43 | 22.17 | 23.81 | 28.21 | 30.05 | 26.58 | 1.33 |
| Miransertib | 10.01 | 12.75 | 5.89 | 22.14 | 24.11 | 31.05 | 17.66 | 3.92 |
| Inhibitor VIII | 164.33 | 197.11 | 160.77 | 225.83 | 163.12 | 245.59 | 192.79 | 14.85 |

**Figure S10**- Replicate data associated with Figure 6C and D. **(A)** Signal buildup curves and fitted $EC_{50}$ values using the K5 tracer on MDA-MB-231 or MDA-MB-468 cells transfected with AKT1-NL or Myr-AKT1-NL mutants. **(B)** Replicate binding curves for AKT inhibitors on AKT1-NL overexpressed in MDA-MB-231 cells. Fitted $IC_{50}$ values from binding curves. Calculated $K_i$ values using fitted $IC_{50}$ values and $EC_{50}$ values from panel (A). **(C)** same as (B) for overexpressed Myr-AKT1-NL in MDA-MB-231 cells. **(D)** same as (B) for overexpressed AKT1-NL in MDA-MB-468 cells. **(E)** same as (B) for overexpressed Myr-AKT1-NL in MDA-MB-468 cells.
